# Functional omics of ORP7 in primary endothelial cells

**DOI:** 10.1101/2024.03.19.585674

**Authors:** Juuso H. Taskinen, Minna Holopainen, Hanna Ruhanen, Reijo Käkelä, Vesa M. Olkkonen

## Abstract

**Background:** Many members of the oxysterol binding protein related protein (ORP) family have been characterized in detail over the past decades, but the lipid transport and other functions of ORP7 still remain elusive. What is known about ORP7 points toward an endoplasmic reticulum and plasma membrane-localized protein, which also interacts with GABARAPL2 and unlipidated LC3B, suggesting a further autophagosomal/lysosomal association. Functional roles of ORP7 have been suggested in cholesterol efflux, hypercholesterolemia, and macroautophagy. We performed a hypothesis-free omics analysis of chemical ORP7 inhibition utilizing transcriptomics and lipidomics as well as proximity biotinylation interactomics to characterize ORP7 functions in a primary cell type, human umbilical vein endothelial cells (HUVECs). Moreover, assays on metrics such as angiogenesis, cholesterol efflux and lipid droplet quantification were conducted.

**Results:** Pharmacological inhibition of ORP7 lead to an increase in gene expression related to lipid metabolism and inflammation, while genes associated with cell cycle and cell division were downregulated. Lipidomic analysis revealed increases in ceramides, lysophosphaditylcholines, as well as saturated and monounsaturated triacylglycerols. Significant decreases were seen in all cholesteryl ester and in some unsaturated triacylglycerol species, compatible with the detected decrease of mean lipid droplet area. Along with the reduced lipid stores, ABCG1-mediated cholesterol efflux and angiogenesis decreased. Interactomics revealed an interaction of ORP7 with AKT1, a central metabolic regulator.

**Conclusions:** The transcriptomics results suggest an increase in prostanoid as well as oxysterol synthesis, which could be related to the observed upregulation of proinflammatory genes. We envision that the defective angiogenesis in HUVECs subjected to ORP7 inhibition could be the result of an unfavorable plasma membrane lipid composition and/or reduced potential for cell division. To conclude, the present study suggests multifaceted functions of ORP7 in lipid homeostasis, angiogenic tube formation and gene expression of lipid metabolism, inflammation and cell cycle in primary endothelial cells, possibly through AKT1 interaction.

## 1. Introduction

Current understanding of the Oxysterol-binding protein-related protein (ORP) family is extensive, but the functions of some of its members still elude researchers. A good example is ORP7 which has been left with little attention, whereas OSBP, from which the family takes its name, has been extensively studied. The primary role of ORPs is to establish membrane contact sites (MCS) where they facilitate the transfer of different lipids between intracellular membranes. A well-studied MCS is formed by OSBP between the ER and *trans*-Golgi, where OSBP mediates the transport of ER cholesterol reciprocally to *trans*-Golgi PI(4)P which is transported to the ER (1). Another well-examined family member is ORP2, which ferries cholesterol from endosomes to the plasma membrane and PI(4,5)P_2_ to endosomes (2–4). Which lipids ORP7 transports and between which membranes remain unclear, as do its other cellular functions. Notably, PubMed searches for ORP7 and OSBPL7 yield 5 and 8 results, respectively, whereas searches for ORP2 and OSBPL2 generate 47 and 51 results, and those for OSBP yield 350 results. The sparse knowledge on ORP7 suggests that it localizes to the PM and ER (5) and interacts with GABARAPL2, formerly GATE16 (Zhong et al., 2011), which has been shown to localize to ER, Golgi and autophagosomes (7,8). More recent studies have unveiled that inhibition of ORP7 leads to an increase in ABCA1 dependent cholesterol efflux to apolipoprotein A1 (ApoA1) in kidney podocytes, while ORP7 inhibition had a weaker impact on cholesterol efflux to high-density lipoprotein (HDL) particles (9). A whole genome sequencing study in a Malaysian cohort showed that participants with a ORP7 c.651_652del variant had 17 times higher odds of hypercholesterolemia than subjects without the variant (10). A large-scale study of the human macroautophagy pathway revealed that ORP7 among other ORPs binds to the unlipidated form of LC3B and is required for macroautophagy (11). A database mining study on pancreatic ductal adenocarcinoma (PDCA) data performed by Chou et al. showed that *OSBPL7* had the highest positive correlation with *OSBPL2* and -*3* and highest negative correlations in expression with *OSBPL8* and -*11* (Chou et al., 2021). Expression of *OSBPL7* was also shown to be positively associated with the infiltration of immune cells. Single nucleotide variations in *OSBPL7* have also been associated with eczema (13) as well as with the uptake of amyloid beta in the brain of a Korean cohort (14). A schematic overview of the aforementioned knowledge on ORP7 has been encapsulated into Figure 1. This figure contains a considerable of speculation as the lipid trafficking properties of ORP7 are unknown and it is unclear if ORP7 traffics between the ER and PM although it localizes to both. Even though ORP7 interacts with both LC3B and GABARAPL2 individually, it is unknown if all three form a complex, as LC3B and GABARAPL2 also interact with each other (15).

**Figure 1.**
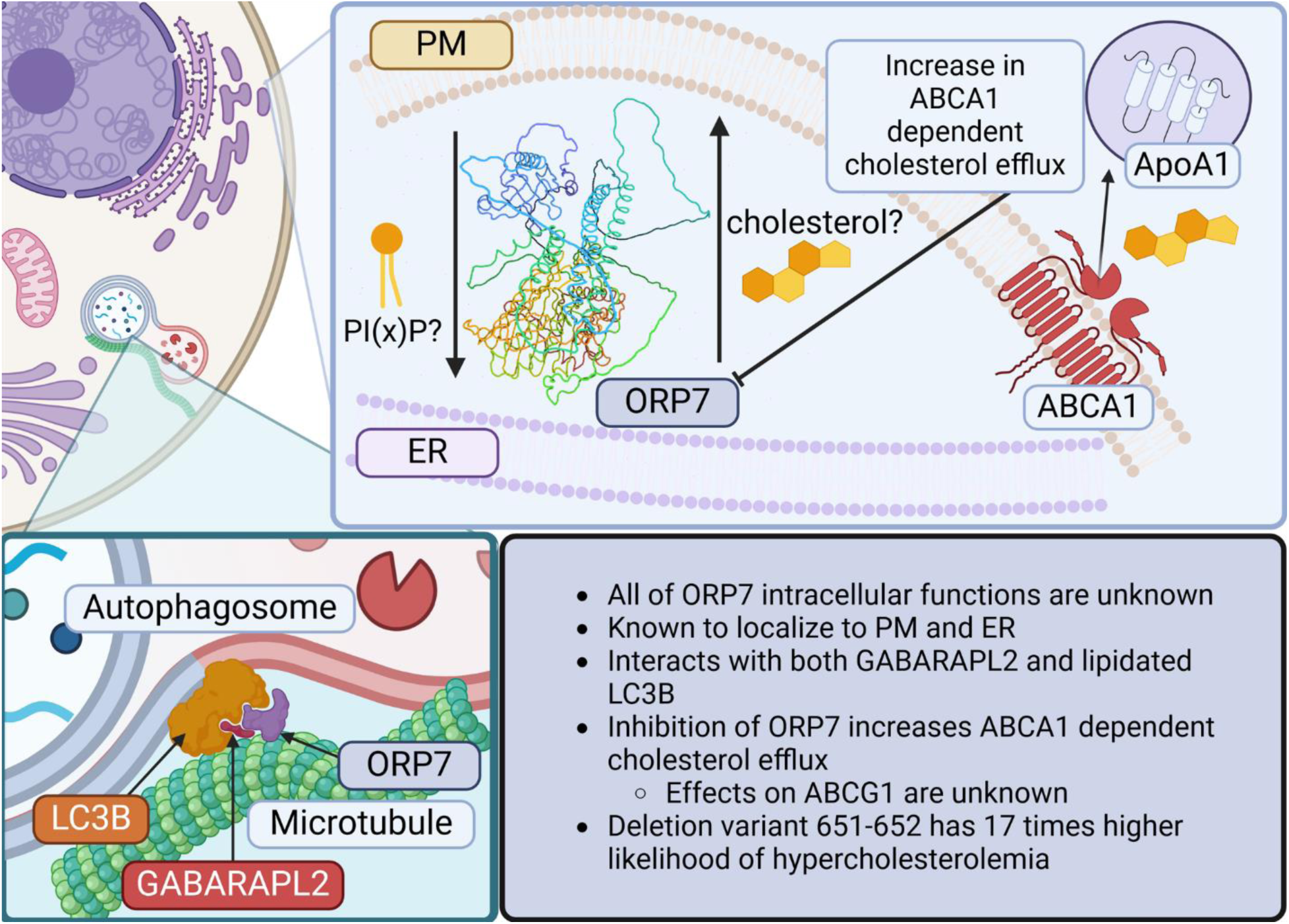
Condensed schematic of current knowledge on ORP7. It is important to note that this figure is highly speculative since many of the processes shown have not been proven by experimental evidence, while parts have been shown to be true *in vitro*, such as the increase in ABCA1 cholesterol efflux (9), interactions of ORP7 between LC3B (15) and GABARAPL2 (6) and its localization to the PM and ER. (Created with BioRender.com)

Given the sparse information on ORP7, we set out to investigate how the manipulation of ORP7 affects the macromolecule pools of HUVECs. To this end we used hypothesis-free omics methods such as transcriptomics, lipidomics and interactomics to get a systemwide view of the changes in manipulated HUVECs. We used a recently discovered 5-arylnicotinamide ORP7 inhibitor CpdG (9) and an ORP7 proximity biotinylation overexpression construct with a mycBirA domain to study these omics in HUVECs. To our knowledge, this is to date the first study to thoroughly investigate the effects of ORP7 manipulation by using multiomics in any cell or animal model.

## 2. Methods

The following section will outline the methods and reagents used during this study; most of the workflows have been condensed and illustrated in Figure 2. The aim was to study the macromolecule profile of ppharmacologicaly ORP7 inhibited HUVECs, to gain a better understanding on the role of ORP7 and MCS in endothelial cells.

**Figure 2.**
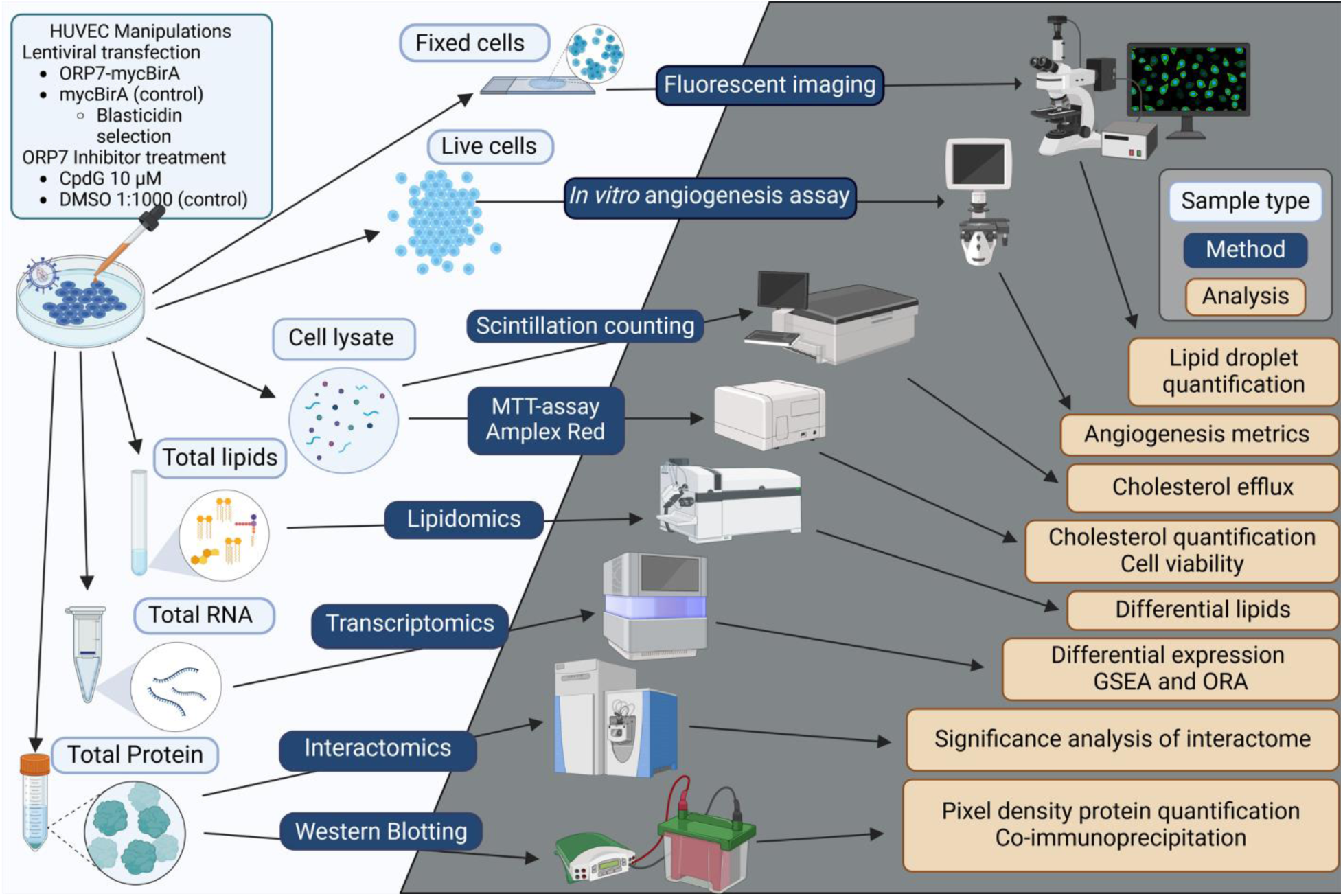
Simplified illustration of the experimental workflow used in this study. Light blue text boxes depict sample types, whereas dark blue text boxes show methods used. (Created with BioRender.com)

### 2.1 cDNA constructs and mRNA quantification by quantitative real-time PCR (qPCR)

cDNA constructs were made in brief as follows: Inserts were produced by PCR using Dynazyme II according to manufacturer’s instructions, using primers 1 and 2 in Table 1.

**Table 1.**
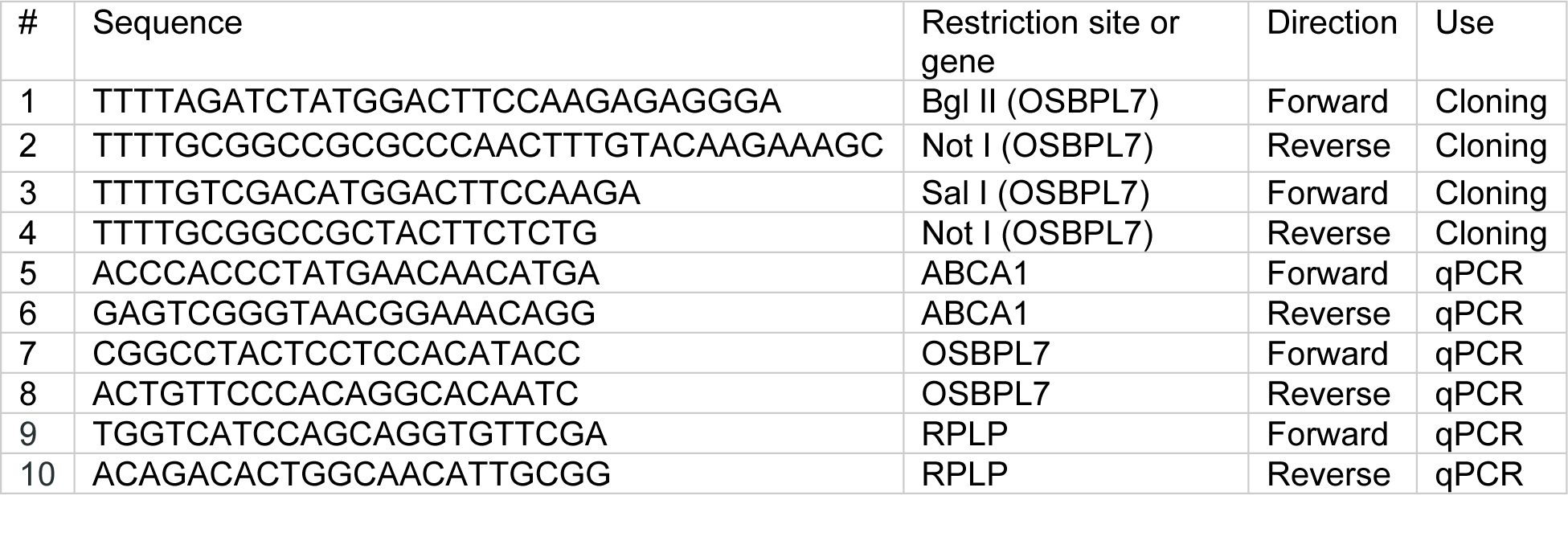
Cloning and qPCR primers used in this study.

An ORP7 cDNA construct in pEGFP-C1 vector and pMycBirA (kind gift from Johan Peränen, Univ. of Helsinki) were digested using Bgl II and Not I and the ORP7 ORF was ligated into pMycBirA. (Froger and Hall, 2007) The obtained plasmids were validated by Sanger sequencing at the Institute for Molecular Medicine Finland FIMM Genomics unit supported by HiLIFE and Biocenter Finland. Sequence reads were mapped to a reference sequence of each plasmid generated with Benchling using UniPro UGENE software (16). Correct constructs where then moved to pENTR2B with Sal I and Not I/Xho I, in a similar manner using primers 3 and 4 in Table 1. The constructs were transferred to pLenti6.3/V5-DEST by Gateway (Thermo Fisher Scientific, Waltham, MA) recombination at the Helsinki University Genome Biology Core (HiLIFE Helsinki and Biocenter Finland).

qPCR validation of select RNAseq observations was carried out on Roche Lightcycler^TM^ 480 II instrument by using Sybr Green chemistry, Roche qPCR Master mix and primers specified in Table 1. *RPLP* was used as housekeeping control mRNA. Fold-changes in gene expression were calculated by employing the -DDC_T_ method.

### 2.2 Cell culture, inhibitor treatment and lentiviral transduction

HUVECs were grown in EGCM2 media (PromoCell cat #: C-22111) without antibiotics, supplemented with endothelial growth mix 2 (PromoCell cat #: C-39216) to passage 6. HUVECs at passage 2, were transduced on 6-well plates using lentiviral particles packaged by the Helsinki Biomedicum Virus Core supported by HiLIFE and the Faculty of Medicine, University of Helsinki, and Biocenter Finland. Cells were incubated with the lentiviruses encoding ORP7-mycBirA (oexORP7) or mycBirA (oexMYC = control) for 48 hours with an approximate MOI of 15. After lentiviral transduction, the cells were selected using media with (1:4000 of 10 mg/mL Blastidicin, Gibco) for five days. Transfected cells were then expanded from 6-well plates to 10 cm plates without antibiotics to passage 6 and used for experiments. For ORP7 inhibition, cells were treated with 10 µM concentration of CpdG (Medchemexpress, cat #: HY-143200, OSBPL7-IN-1) or with 1:1000 dilution of DMSO unless otherwise specified.

### 2.3 MTT assays

Metabolic activity of CpdG or DMSO treated and oexORP7 or oexMYC cells was measured using CellTiter 96® AQueous One Solution Cell Proliferation Assay (Promega cat #: G358C), according to the manufacturer’s instructions. Absorbance at 490 nm was measured using a PerkinElmer EnSpire Multimode Plate Reader. Percentage of metabolic activity was calculated by the following formula: 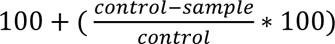, where control is the absorbance of normal HUVECs or oexMYC and samples oexORP7 cells or cells treated with CpdG at 1,5,10,25,50,75,100, 250 and 500 µM concentrations for 24 h.

### 2.4 Western Blotting

Oex and CpdG or DMSO treated cells (24 h) were washed twice with ice cold PBS, then lysed to 200 µL lysis buffer [150 mM NaCl, 0.5 mM MgCl_2_, 0.5 % Triton X-100, 0.5% sodium deoxycholate (v/v)] with 1X EDTA free Protease inhibitor cocktail (Merck cat #: 04693159001) and with or without 1X phosphatase inhibitor (Roche, cat #: REF 04 906 845 001, lot #: 37590700) for 20 min at +4°C. Lysates were collected to a fresh tube and spun for 15 min at 13,000 x g at +4°C, after which supernatant was moved to a fresh tube. Sample protein concentrations were measured with Thermo Fisher Pierce™ BCA Protein Assay Kit (cat #: 23225), and the samples were blotted with a BioRad Trans-Blot Turbo Transfer System. Membranes were blocked for 1 h at room temperature (RT) in 5% bovine serum albumin (BSA) for phosphor-specific antibodies or for 15 min in BioRad EveryBlot solution and then probed overnight at +4°C with antibodies shown in Table 2. Membranes were washed thrice for 10 min with 1 X TBST at RT and then incubated with 1:2000 dilutions of HRP-conjugated secondary antibodies in Milk-TBST or EveryBlot for 1 h at RT. Membranes were washed as above and detected using Thermo Fisher Pierce ECL Western blotting substrate (cat #: 32106, lot #: WJ335099) according to the manufacturer’s instructions.

**Table 2.** Antibodies used in this study.

| Protein | Dilution | Manufacturer | Catalog number | Lot number |
| --- | --- | --- | --- | --- |
| ORP7 | 1:1000 | Sigma Aldrich | HPA036076 | NA |
| pFAK | 1:500 | Invitrogen | 44-626G | NA |
| FAK | 1:1000 | Invitrogen | AHO0502 | NA |
| pAKT1 | 1:1000 | Cell Signalling | 4051S | NA |
| AKT1 | 1:1000 | Invitrogen | AH01112 | YG373683 |
| ACTB | 1:5000 | Sigma Aldrich | A2066-2ml | 106m4770V |
| cMYC | 1:1000 | Santa Cruz Biotechnology | sc-40 | NA |

### 2.5 Co-immunoprecipitation

Oex cells or normal HUVECs were grown on 10 cm plates until 80% confluent, after which they were washed twice with 1X cold PBS on ice. Cells were lysed to 1 mL of the lysis buffer described in section 2.3 that included both protease and phosphatase inhibitors and incubated for 30 min at + 4°C. Lysates were spun for 15 at 13,000 x g at +4 °C and supernatant was collected to a fresh tube. Magnetic beads were washed twice with 1 X TBST, after which 25 µL were added to 500 µL of cell lysates and mixed on a roller for 1 h at +4 °C. Either 5 µL of Mouse HRP antibody or 10 µL of AKT1 antibody (see section 2.3) were added to 25 µL of washed beads, and mixed on a roller for 2h at +4 °C. Beads were removed from each mix by incubating on a magnetic rack for 2 min at +4 °C, after which antibody coupled beads were added to the pre-cleared lysates and incubated o/n at +4 °C. Lysates were removed to a fresh tube by incubating on a magnetic rack and beads were washed thrice for 5 min with lysis buffer at +4 °C. To the washed beads 40 µL of 2 X Laemmli buffer (BioRad) was added and incubated for 10 min at +50 °C. Magnetic beads were removed as described before and to each elution 1 μL of 2-β-mercaptoethanol was added, and samples were boiled for 5 min at +100 °C. Samples were spun down and Western blotting was performed as described in section 2.3.

### 2.6 Angiogenesis assay

Angiogenesis assay (Millipore cat #: ECM625, lot #: 3183813) was performed on oex cells or cells treated for 24 h with CpdG or DMSO at 10 µM or 1:1000 concentrations, respectively. Oex cells were incubated in normal EGCM2 media. Cells were imaged by using a bright-field set-up of EVOS M5000 (Invitrogen) microscope after 8 h. Cell junctions were counted from binary images using Fiji (17) Angiogenesis analyzer plugin (18).

### 2.7 Measurement of cholesterol concentration

Amplex Red^TM^ cholesterol assay (Invitrogen, cat #: A12216, lot #: 2422702) was performed according to manufacturer’s instructions from undiluted samples of oex cell lysates or 1:4 dilutions of CpdG or DMSO treated cell lysates, made as in section “*2.3 Western Blotting”*. Assays were performed for total cholesterol (TC) with cholesterol esterase treatment or free cholesterol (FC) without esterase treatment, and cholesterol concentrations were normalized to protein concentrations measured as in section “*2.3 Western Blotting”*.

### 2.8 Cholesterol efflux to ApoA1 and HDL

Normal HUVECs or oex cells were plated at 50,000 cells per 12-well plate and left to adhere for 24 h, after which the cells were washed twice with 1 PBS and then 1 mL of media with 0.2 µCi/mL of ^3^H-cholesterol and 5 mg/mL Sandoz 58-035 SOAT inhibitor was added. After a 24 h incubation media was removed and cells were washed thrice with 1 X PBS, after which 2 mL of FBS-free EGCM (Sigma Aldrich, cat #: 211F-500) media with 1 mg/mL of BSA was added. To the HUVEC media either 10 µM of T0901317, 10 µM CpdG or 1:1000 (v/v) DMSO was added and incubated for 24 h. After incubation, a 1 mL aliquot was taken from each well and spun at 800 x g for 5 min, and the supernatant was collected to a scintillation tube. For ApoA1 mediated efflux 100 µL of FBS free media with 50 µg/mL ApoA1 was added and for HDL mediated efflux 111 µL of 450 µg/mL HDL isolated from human plasma and dialyzed to PBS was added and incubated for 24 h. Finally, media was collected and spun as before, and supernatant was collected to scintillation tubes. Cells were lysed to 300 µL of 1% SDS and collected to scintillation tubes. To each scintillation tube 3 mL of scintillation cocktail (Ultima Gold, cat #: 6013326, lot #: 23151, Perkin Elmer) was added and tubes were mixed briefly before they were subject to liquid scintillation counting with a Wallac Winspectral 1414 counter (Perkin-Elmer, Waltham, MA).

### 2.9 Immunofluorescence staining, fluorescence microscopy and lipid droplet quantification

HUVECs were seeded at 10,000 cells/coverslip and left to adhere for 24 h in 50 µL of EGCM2. Oex cells and normal HUVECs were left to adhere for 24 h, after which the oex cells were fixed and normal HUVECs were incubated for another 24 h in 100 µL of media containing 10 µM of ORP7 inhibitor or 1:1000 DMSO. Cells were fixed with 4% PFA in PBS and blocked in 1% BSA in PBS. Lipid droplet staining was performed by incubating fixed cells for 1 hour at RT with 1:1000 of BODIPY493/503 stock (50 mM) and mounted with Mowiol:Dabco:DAPI solution. Imaging of the specimens was carried out on a Zeiss Observer.Z1 fluorescence microscope (Serial #: 3834002509, Zeiss Group), with a 63 X immersion oil objective. Lipid droplets were quantified as specified in (19).

### 2.10 Lipidomics

Lipids from oex and CpdG or DMSO treated cells were extracted according to Folch *et al.* (Folch, Lees and Sloane Stanley, 1957). Solvents were evaporated and the lipid extracts immediately dissolved in chloroform/methanol 1:2 (by volume). Internal standard mixture (SPLASH® LIPIDOMIX® and Cer 18:1;O2/17:0, both from Avanti Polar Lipids, Alabaster, AL, USA) were added to the samples, which were analysed with LC-MS/MS as previously described (21). In brief, samples were analysed with acetonitrile/water/isopropanol-based solvent system (22) by employing Agilent 1290 Infinity HPLC (Agilent Technologies, Santa Clara, CA) equipped with a Luna Omega C18 100 Å (50 x 2.1 mm, 1.6 μm) column (Phenomenex) and Agilent 6490 Triple Quad LC/MS with iFunnel Technology. The lipids species were identified and quantified using lipid class-specific detection modes, as previously described(21) Retrieved spectra were processed by Mass Hunter Workstation qualitative analysis software (Agilent Technologies, Inc.), and individual lipid species were quantified using the internal standards and LIMSA software (23).

### 2.11 Transcriptomics

RNA was extracted from oex, CpdG or DMSO treated cells with the RNAeasy Mini kit (Qiagen cat #: 74104) according to the manufacturer’s instructions, with the following modifications: Samples were homogenized by passing lysate through a 200 µL pipette tip 10 times, on-column DNAse 1 digestion was performed according to manufacturer’s instructions, and samples were eluted to 30 µL of RNAse free H_2_O. RNA integrity, library preparation and RNA sequencing were performed according to the following instructions by GENEWIZ Germany GmbH (Leipzig, Germany). RNA samples were quantified using Qubit 4.0 Fluorometer (Life Technologies, Carlsbad, CA, USA) and RNA integrity was checked with RNA Kit on Agilent 5300 Fragment Analyzer (Agilent Technologies, Palo Alto, CA, USA). RNA sequencing libraries were prepared using the NEB Next Ultra RNA Library Prep Kit for Illumina following manufacturer’s instructions (NEB, Ipswich, MA, USA). Briefly, mRNAs were first enriched with Oligo(dT) beads. Enriched mRNAs were fragmented for 15 minutes at 94 °C. First strand and second strand cDNAs were subsequently synthesized. cDNA fragments were end repaired and adenylated at 3’ends, and universal adapters were ligated to cDNA fragments, followed by index addition and library enrichment by limited-cycle PCR. Sequencing libraries were validated using NGS Kit on the Agilent 5300 Fragment Analyzer (Agilent Technologies, Palo Alto, CA, USA), and quantified by using Qubit 4.0 Fluorometer (Invitrogen, Carlsbad, CA). The sequencing libraries were multiplexed and loaded on the flow cell on the Illumina NovaSeq 6000 instrument according to manufacturer’s instructions. The samples were sequenced using a 2×150 Pair-End (PE) configuration v1.5. Image analysis and base calling were conducted by the NovaSeq Control Software v1.7 on the NovaSeq instrument. Raw sequence data (.bcl files) generated from Illumina NovaSeq was converted into fastq files and de-multiplexed using Illumina bcl2fastq program version 2.20. One mismatch was allowed for index sequence identification.

### 2.12 Biotin proximity labeling interactomics

Interactomics was performed on HUVECs transfected with oexORP7-mycBirA or control oexmycBirA vectors detailed in section “2.1. cDNA constructs”. Transfected cells on 6-well plates were selected for approximately 5 days with 10 µg/mL of blasticidin until all wild type HUVECs were dead, and then left to grow until confluent, after which cells were expanded to passage 6. Samples were created as described in (24) section “*Procedure*” steps 47 – 60, from two 10 cm plates of transfected and selected cells for each sample. LC-MS/MS was performed by University of Helsinki Viikki Proteomics Unit (Helsinki, Finland) as described in (24).

### 2.13 Bioinformatics and statistics

RNA-seq analysis was performed according to the following pipeline using the Chipster suite (25): Pair ended reads were clipped with Triommatic (phred >= 30) (26), then aligned to the human genome (GRCh38) with STAR (27), aligned reads were counted using HTSeq (28), and differential expression-analysis was performed using DESeq2 (29) and log2 fold change shrinkage was estimated using apeglm (30). Overrepresentation analysis (ORA) was performed using all genes that had an adjusted p-value of less than 0.05, gene set enrichment analysis (GSEA) was performed using the whole RNAseq data set and ranked according to log2 fold change, ties were resolved at random. Both ORA and GSEA were performed using the R package ClusterProfiler (31). Lipidomic data analysis was performed using the Bioconductor package LipidR (32) with the following caveats: Lipid concentrations were normalized to a normalization factor calculated by dividing the protein concentrations of each sample by the median of all sample protein concentrations. Concentrations were then multiplied by 1000 and log2 transformed for further analysis. To decrease false discovery rate, all p-values in omics results were corrected by using the Benjamini-Hochberg correction. T-test and ANOVA comparisons between groups were made using R package ggpubr (stat_compare_means function). PC-plots were generated using R package factoextra. Interactomics analysis was performed using the Crapome server for SAINT (33).

## 3. Results

As we describe in the method section, we performed all, except interactomics, analyses on both treated and oex cells, but we have chosen to focus on the results acquired for CpdG-treated cells. Data from the oex cells are discussed when they are relevant. Results from the oex cells showed little to no significant changes in most metrics measured, therefore we have chosen to relegate those data to the supplementary material in order to keep the present article more concise.

### 3.1 HUVECs exhibit a high tolerance for ORP7 inhibition and overexpression

When inhibitors are used in a new cell model, it is vital to study how they affect cellular health to discover a concentration where the cells show little to no toxicity and the inhibitor shows strong inhibition of its target. To this end, we performed the well-established MTT assay, to measure metabolic activity of HUVECs treated with CpdG or the vehicle control DMSO as well as overexpression of ORP7-mycBirA construct (oexORP7) compared to the mycBirA control (oexMYC). These results are exhibited in Figure 3A and B respectively.

**Figure 3.**
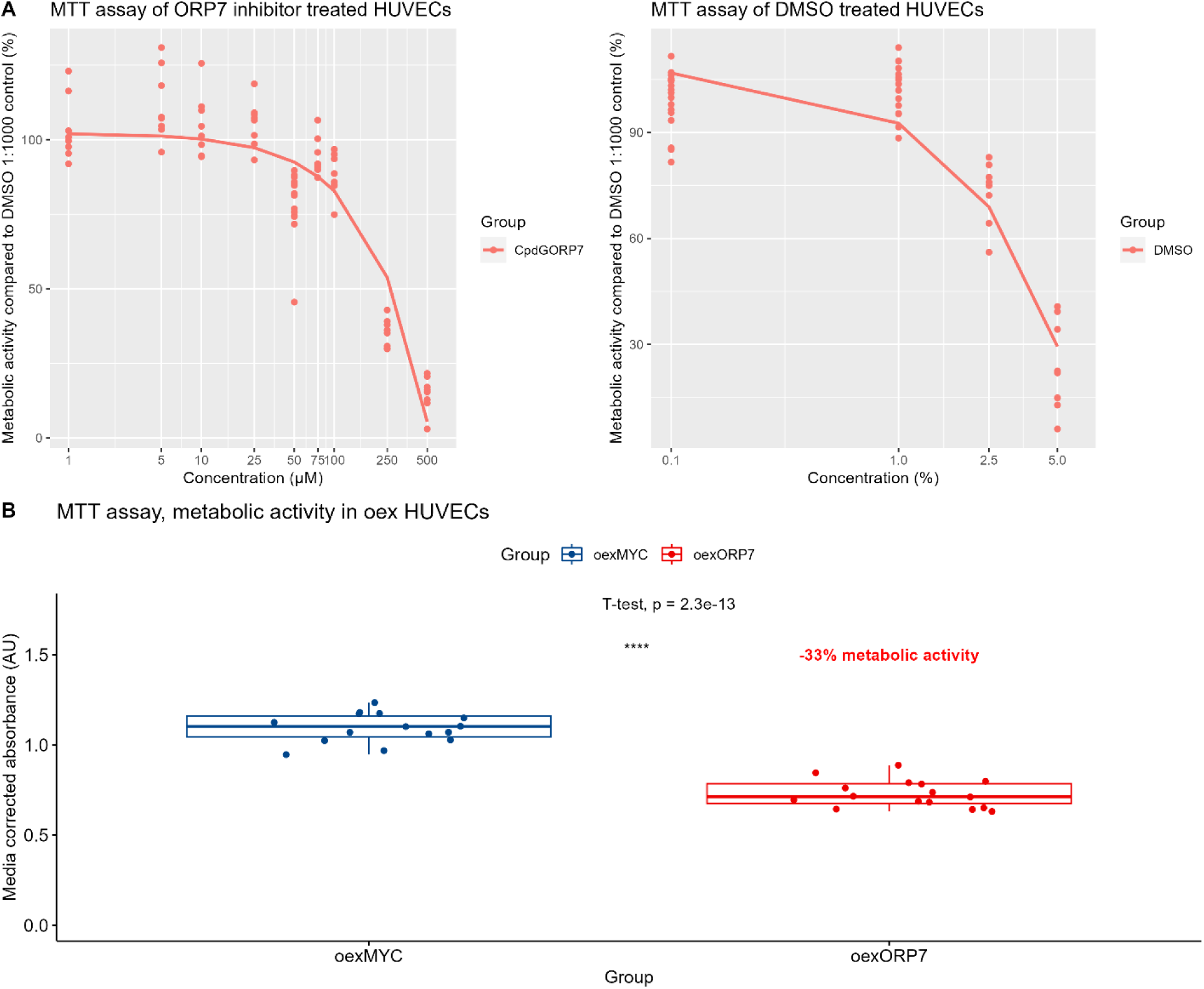
Metabolic activity (MTT-assay) in CpdG or DMSO treated and oex cells. A) Y-axis represents the % of metabolic activity compared to vehicle control (1:1000 DMSO), and the X-axis depicts either the concentration of CpdG in µM or DMSO in v/v-%. N = 8 per concentration. B) Metabolic activity in oexORP7 or oexMYC HUVECs, Y-axis represents absorbance in arbitrary units, and X-axis each group measured. In red is displayed the calculated reduction in metabolic activity in oexORP7 cells as compared to oexMYC control cells. N = 14 per group. P-values were determined using Student’s T-test.

CpdG was well tolerated by HUVECs and a 50% reduction in metabolic activity was reached at 250 µM concentration, whereas the third highest concentration of 100 µM displayed similar reductions in metabolic activity to all the lower concentrations of CpdG (Figure 3A). oexORP7 cells exhibited a small but statistically significant 33% decrease in the metabolic activity compared to oexMYC control (Figure 3B). We chose to continue further experimentation with 10 µM concentration of CpdG as it did not affect HUVEC metabolism in a deleterious manner, and this concentration has been shown to be biologically relevant in THP1 cells (Wright et al., 2021).

### 3.2. Quality control of overexpression treatment, and omics results

To ensure the quality of results from the experiments detailed in further sections, we performed rigorous quality control of both CpdG-treated and oexORP7 cells. We examined the mRNA levels of *ABCA1* and *OSBPL7* in CpdG-treated cells with qPCR, as *ABCA1* expression has been shown to increase with ORP7 inhibition (9). The CpdG-treated cells had statistically significantly higher ABCA1 expression (fold change 1.36) than the DMSO-treated cells (Figure 4A), but the parallel slight increase of OSBPL7 was not significant (Figure 4B). At the protein level, the western blot analysis verified the overexpression of oexORP7 and oexMYC in the corresponding transduced cells (Figure 4C). The level of ORP7-mycBirA overexpression was approximately 16-fold as compared to the endogenous ORP7 in wild type cells.

**Figure 4.**
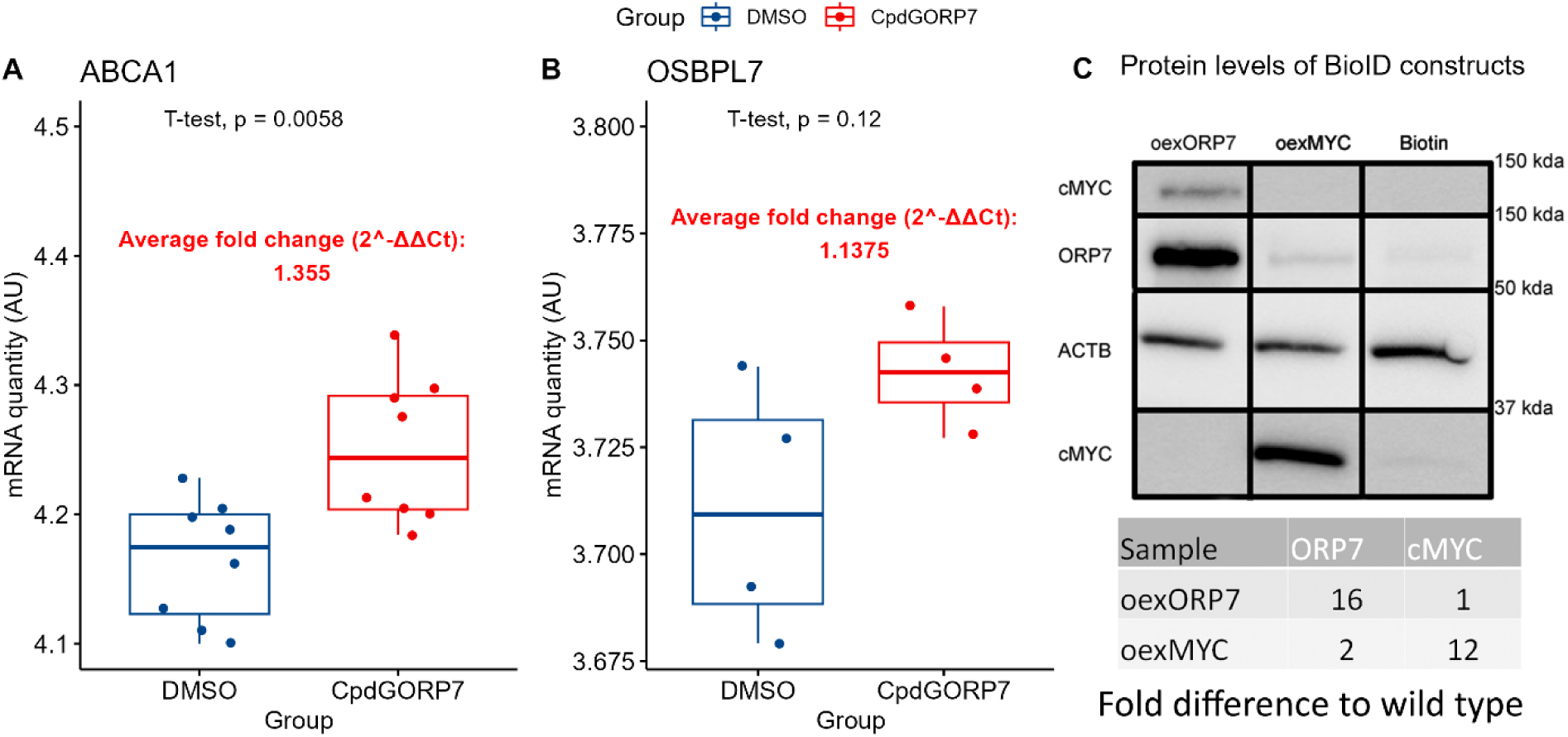
Results of qPCR from CpdG and DMSO treated cells and Western blotting of ORP7 or MYC overexpressing cells. A) mRNA quantity of *ABCA1* in arbitrary units. N = 8 per group B) mRNA quantity of *OSBPL7* in arbitrary units, showing a statistically non-significant average fold change of 1.13. N = 4 per group C) Western blotting results of overexpressing cells. The rows from top to bottom are; protein levels of cMYC levels in oexORP7 cells, ORP7, β-Actin (ACTB) and cMYC. The left-most lane is for oexORP7 cells, the second lane displays the same in oexMYC cells and the last lane depicts the same but in wild type cells treated with 200 mM biotin (Biotin). Fold differences of proteins shown in the table were calculated from β-Actin normalized pixel densities compared to wild type.

To ensure the quality of our omics results, we performed principal component analysis (PCA) to verify if our manipulations resulted in variation between sample groups in all omics experiments performed (Figure 5).

**Figure 5.**
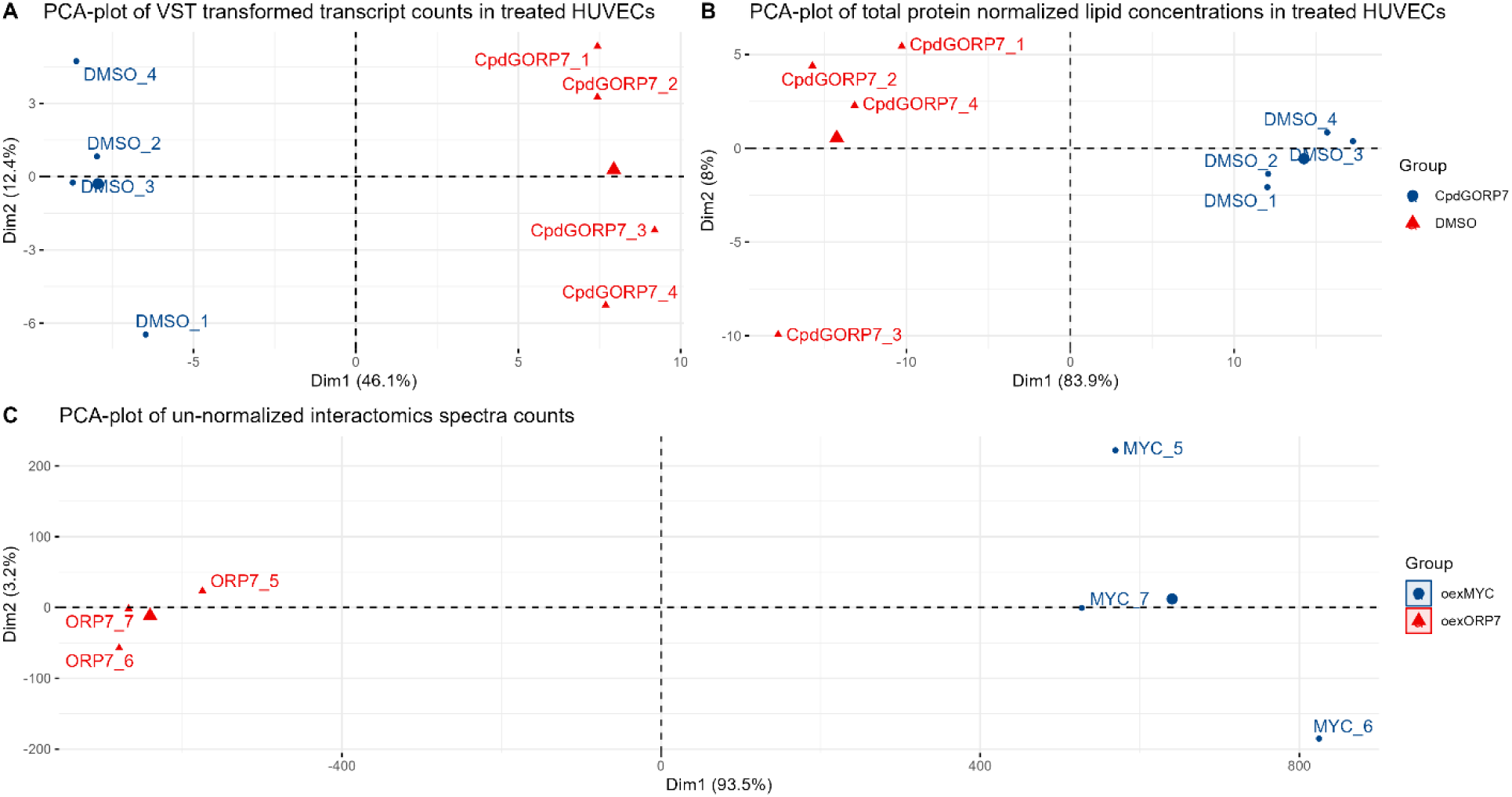
PCA-scores plots of omics results used in this study. All plots show controls (DMSO or oexMYC) in blue and CpdG inhibition or ORP7 overexpression in red. Each axis shows the two largest components where percentage in brackets is the amount of variation explained by each dimension. A) PCA-plot of variance stabilizing transformed transcript counts in CpdG- or DMSO-treated HUVECs. B) Total protein normalized lipid concentration in CpdG- or DMSO-treated HUVECs. C) PCA-plot of un-normalized unique spectra counts in proximity based interactomics samples. In all plots, samples are shown to group together along the largest dimension, with more spread along the second largest dimension.

When using as PCA loadings either transcript counts, lipid concentrations or interactome unique spectra counts, in each experiment all samples resembled the samples of their own treatment group, and treatment or overexpression groups separated along the dimension, which has the largest explanatory power (Figure 5). This indicates that the variation between the biochemical profiles of samples was mostly due to treatments or overexpression.

### 3.3 Transcriptomics reveals multifaceted changes in CpdG-treated HUVECs

Transcriptomics is a convenient method to study the intermediate steps from genotype to phenotype. It can give insights into how cells compensate for the effects of a test variable. We performed a comprehensive gene set analysis on CpdG-treated differential expression data compared to DMSO control, in order to reach a better understanding of how HUVECs compensate for the CpdG treatment (Figure 6). Wikipathways gene sets, that had overrepresentation of downregulated genes, were related to: cell cycle, DNA repair, cancer, and nucleic acid metabolism (highlighted with blue in the left facet of Figure 6). Overrepresentation in upregulated genes were mostly related to lipid metabolism gene sets, such as cholesterol biosynthesis, eicosanoid metabolism, synthesis of n-3 and n-6 polyunsaturated fatty acids, and oxysterols derived from cholesterol. Inflammation-related sets were also present, which include sets such as IL-18 signaling pathway, platelet-mediated interactions and Netrin-UNC5B signaling pathway. Gene set related to mitochondria were also present and these included gene sets such as Oxidative stress response, Electron transport chain OXPHOS system in mitochondria, Mitochondrial complex 1 assembly model OXPHOS system and NRF2 pathway (highlighted with red in the right facet of Figure 6). Similar analysis done on other databases and on data from overexpressing cells is shown in Supplementary Figures S14 - S29.

**Figure 6.**
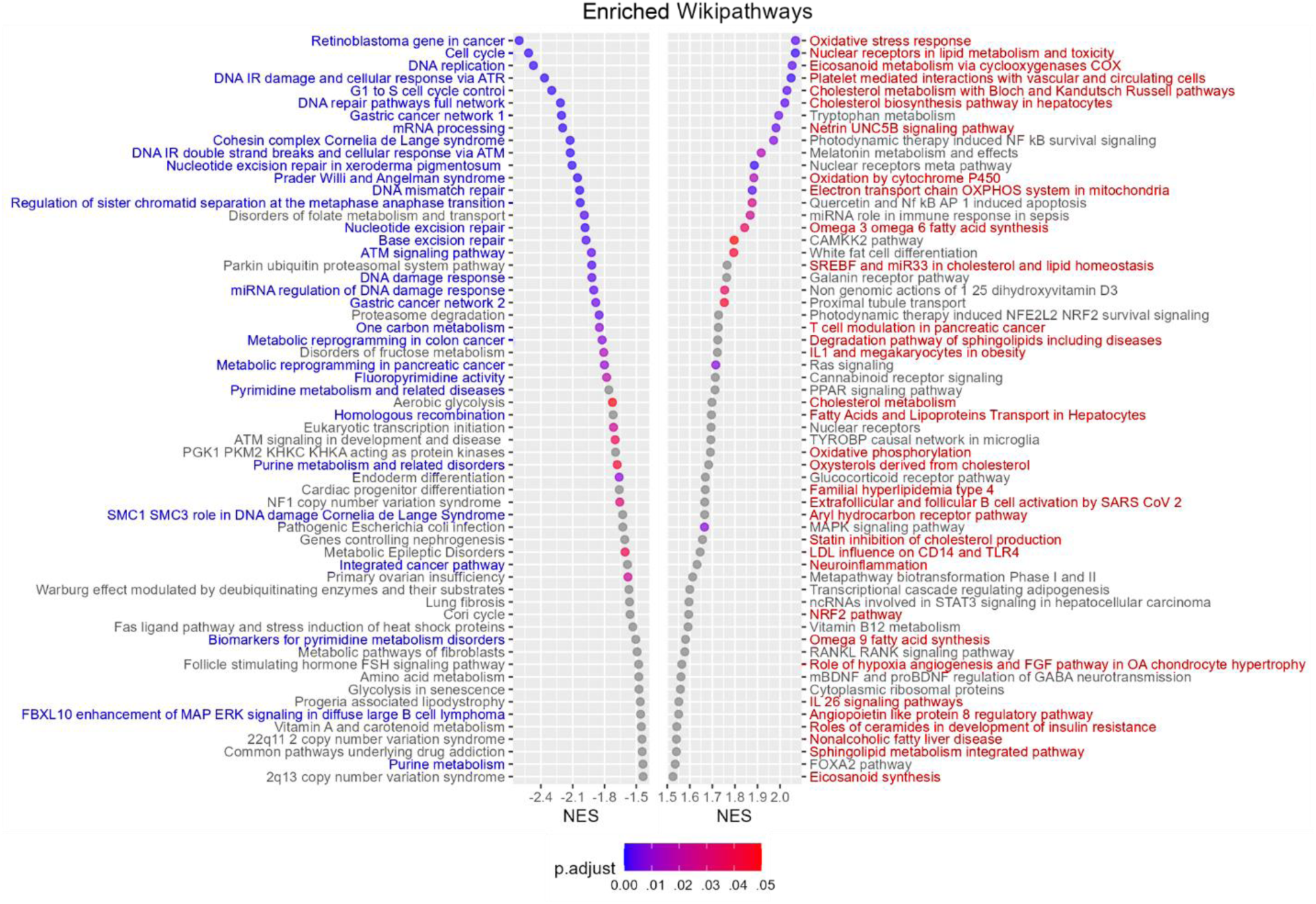
Subset of GSEA results of differentially expressed genes from CpdG-treated using Wikipathways database. Y-axis represents enriched gene sets, and X-axis the normalized enrichment score (NES) for each gene set, where a negative value indicates overrepresentation in highly downregulated and positive values highly upregulated genes. The color of each dot represents the Benjamini-Hochberg adjusted p-value for each gene set. Highlighted with blue font are the gene sets related to cell cycle, cell division, cancer, DNA repair and nucleotide metabolism. Highlighted with red font are the gene sets related to lipid metabolism, mitochondrial function, and inflammation.

Gene set analysis is a convenient and robust way to produce systematic understanding of the basis of functional changes, but understanding gene level changes provides a more detailed and precise understanding into a differential expression data set. Examining the changes in individual genes can reveal variations in, for example, key pathway genes or rate limiting enzymes that affect whole pathways. The same goes for transcriptional factors, which when differentially expressed can have significant downstream effects. Therefore, we also plotted the differentially expressed genes with a conventional volcano plot (Figure 7).

**Figure 7.**
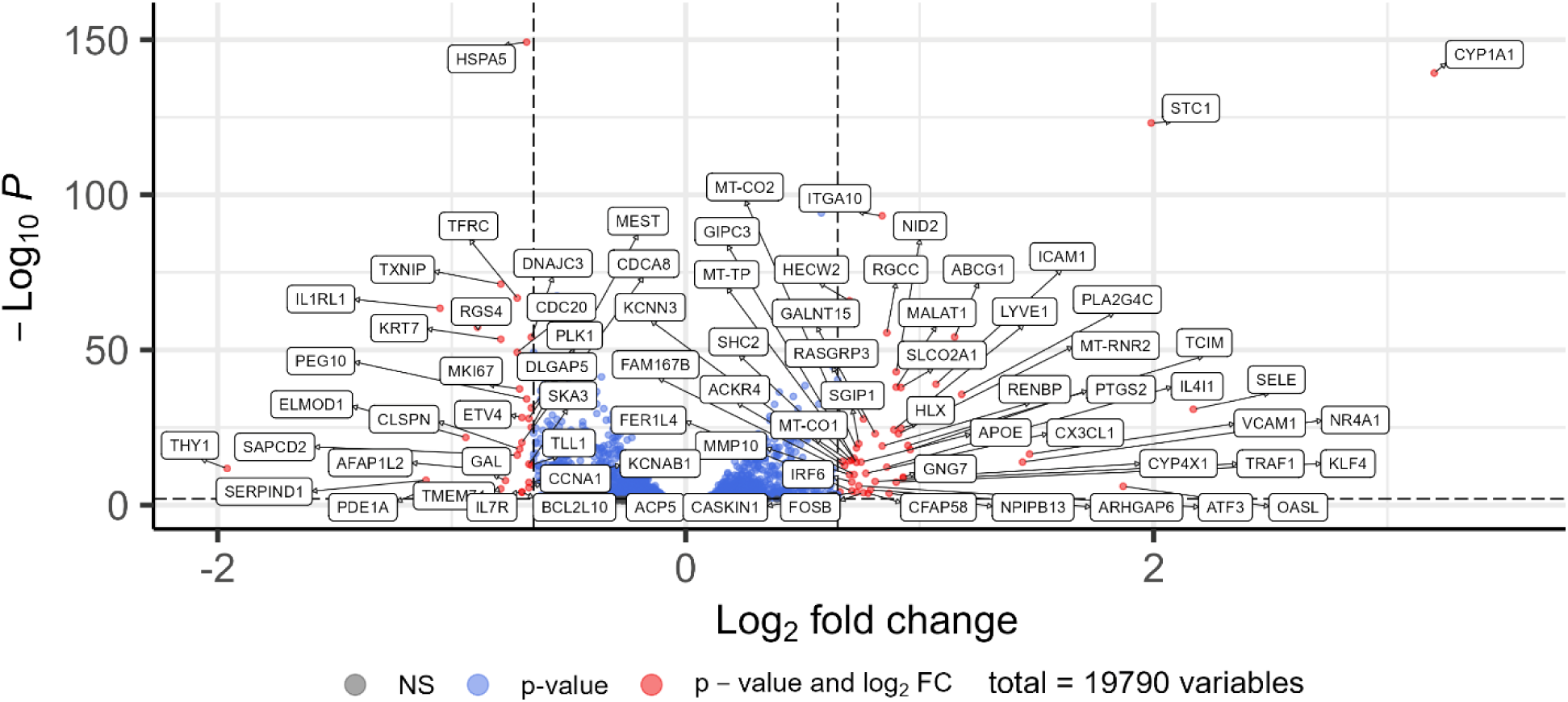
Volcano plot of differential gene expression results in CpdG-treated cells compared to DMSO control. X-axis represents log2 fold change and Y-axis -log10 adjusted p-value. Annotated red dots show genes that have an absolute log2 fold change of 0.75 or higher and are significantly altered, blue dots have an absolute log2 fold change which is lower than 0.75 and are significantly altered, grey dots represent genes that have not been significantly altered.

Interestingly, many genes related to prostaglandin metabolism were upregulated. These included *CYP1A1*, *SLOC2A1* and *PTGS2* (34–36) as well as phospholipase, lysophospholipase and O-acyltransferase PLA2G4C, which release arachidonic acid (AA) from phosphatidylcholine (37–39). Other notable upregulated genes were *SELE*, *VCAM1* and *ICAM1*, which are related to endothelial cell inflammation and early atherogenesis (40–43). A similar expression patter is visible in other inflammation and atherosclerosis related genes such as *APOE*, *KLF4*, *CXC3L1*, *STC1*, *CH25H*, and *ABCG1* (44–53). Downregulation was evident in inflammation related genes such as, *THY1*, *IL1RL1*, *IL7R* (54,55), as well as in cell adhesion related genes such as *FERMT1* (56). Log2 fold change and statistics for the aforementioned genes and other genes related to, for example cholesterol synthesis, autophagy and inflammation can be found in table 3.

**Table 3.**
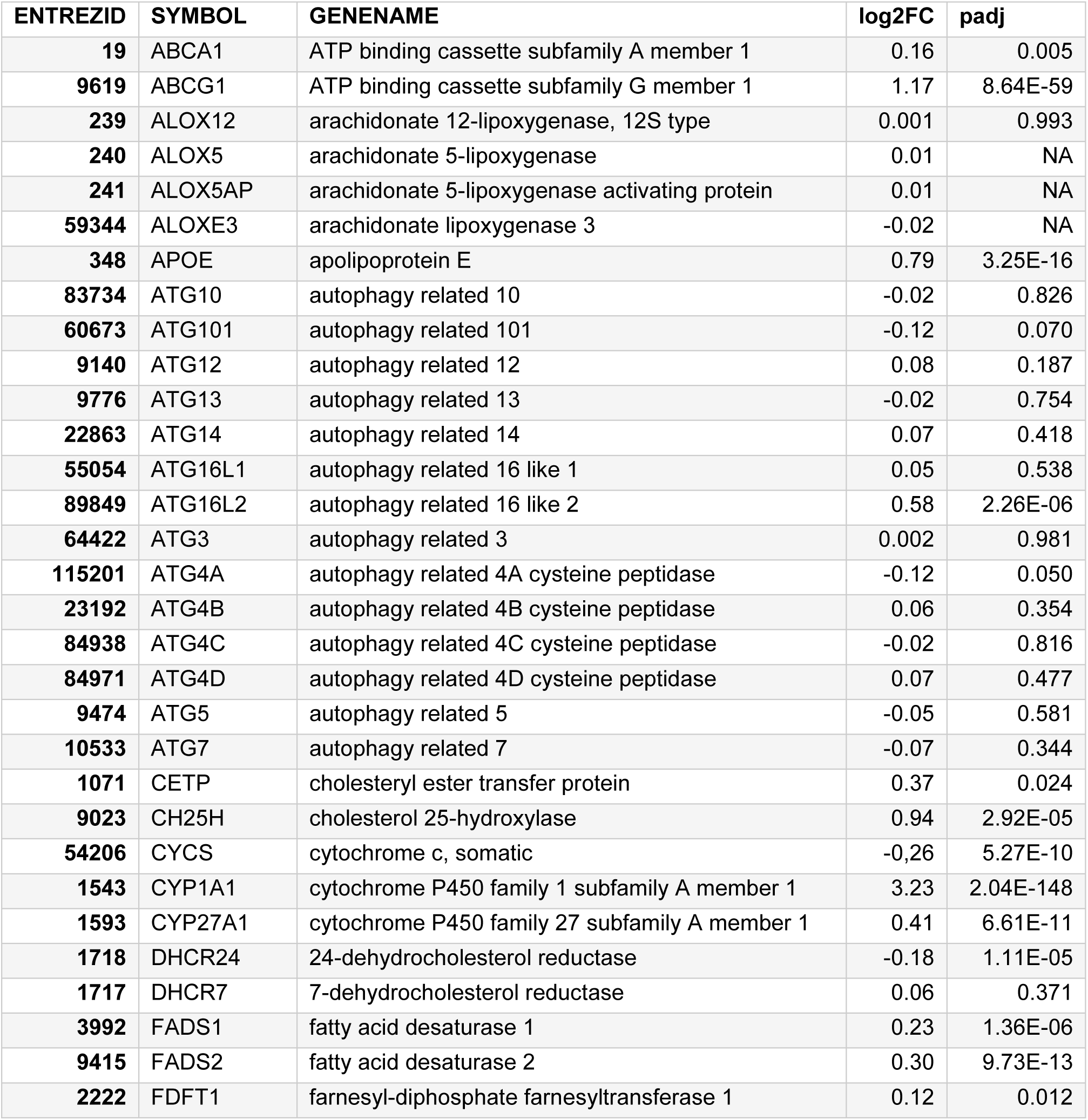
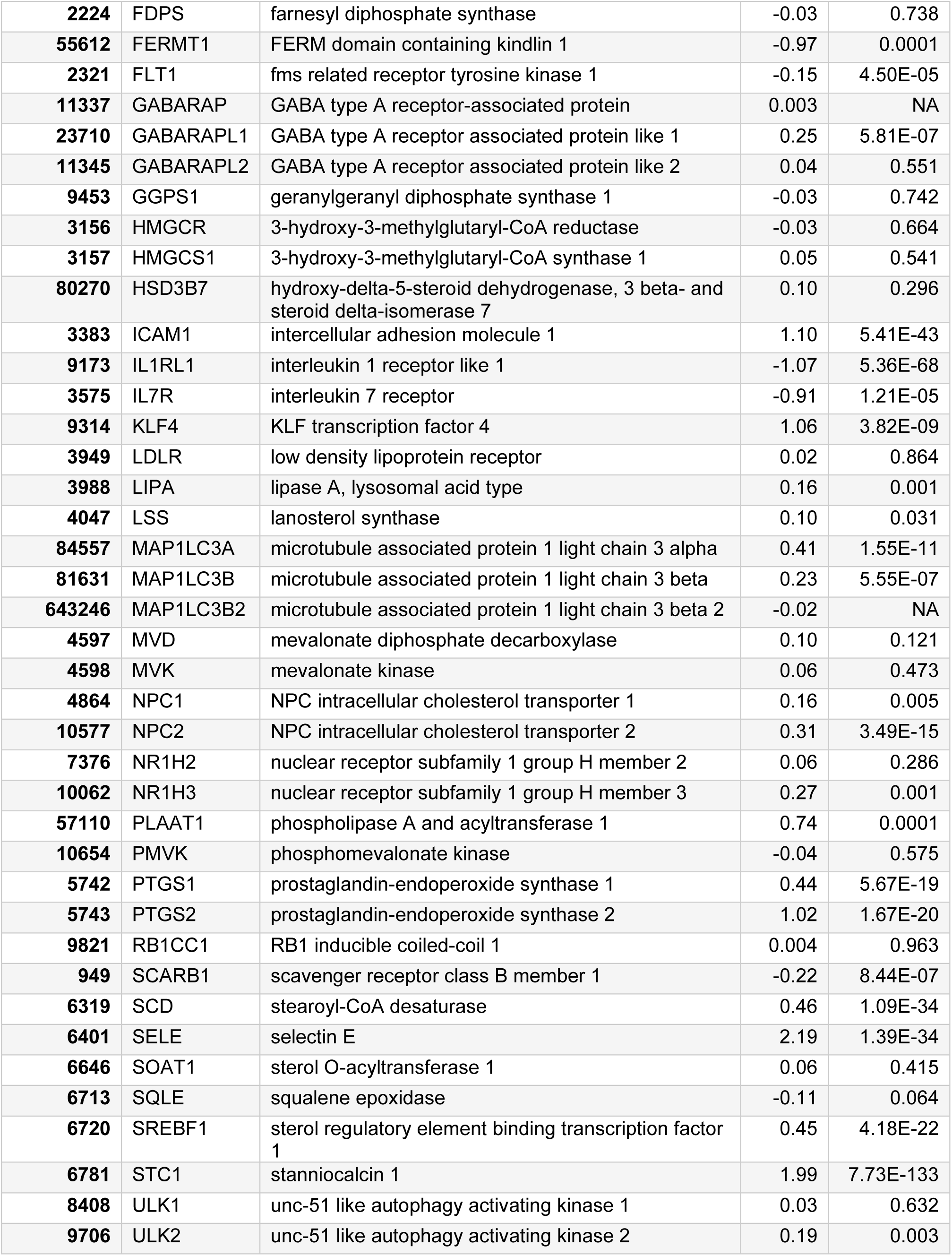
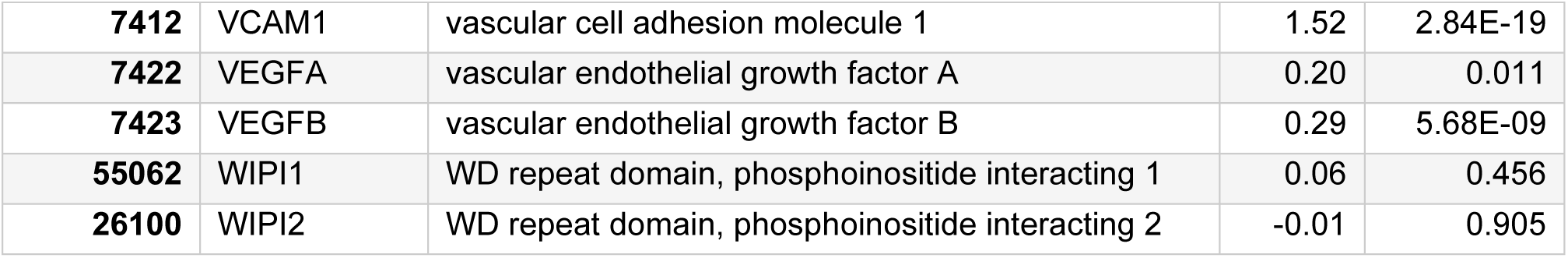
Differential expression metrics for selected genes related to lipid metabolism, cholesterol metabolism, inflammation, mammalian autophagy (57) and endothelial cell function. Log2 fold change (log2FC) and Benjamini-Hochberg adjusted p-values (padj) are shown.

### 3.4 Manipulated HUVECs have reduced angiogenic capacity

Endothelial cells have multiple crucial functions such as angiogenesis, i.e. the formation of new blood vessels from existing ones (58). We used a matrix-based angiogenesis assay to measure how CpdG treatment affects angiogenesis in HUVECs. Inhibition of ORP7 reduced angiogenetic capacity in all metrics measured but did not block it entirely. The number of junctions, branching length and segment length of the newly formed vessels were all statistically significantly decreased in the CpdG-treated cells compared to the DMSO controls (Figure 8). The definitions of each angiogenic metric are detailed in the documentation of Angiogenesis Analyzer (18).

**Figure 8.**
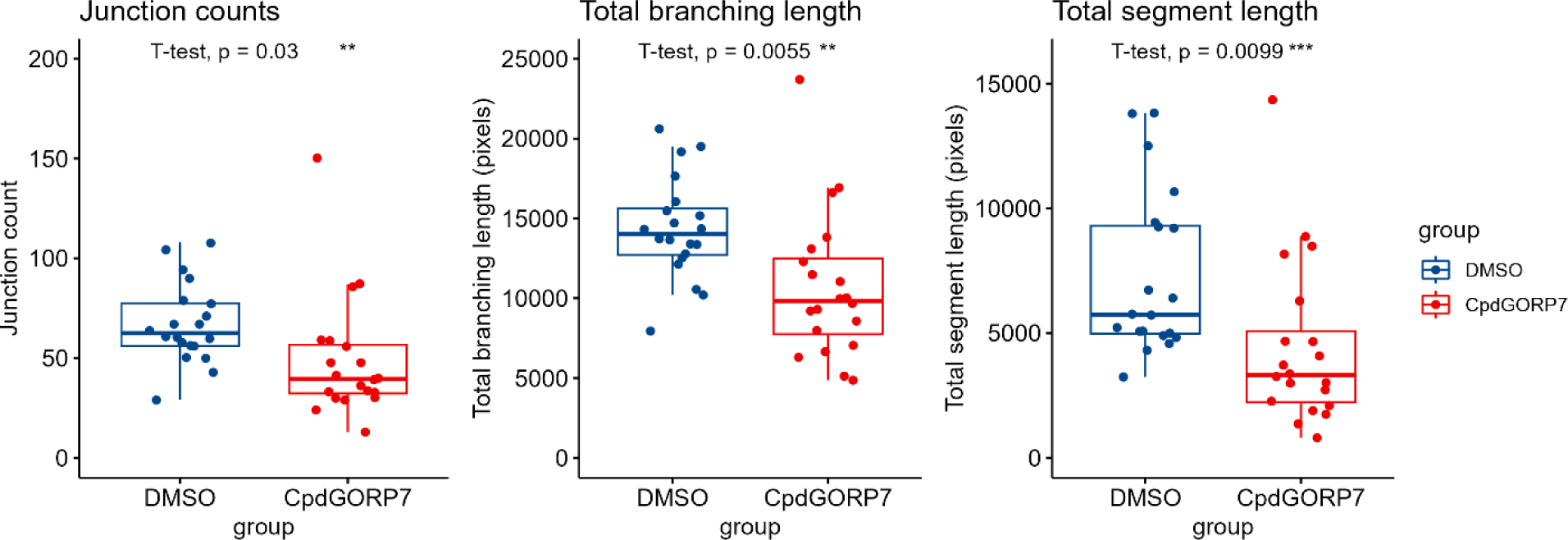
Angiogenic metrics from CpdG or DMSO treated cells grown on angiogenesis inducing matrix. In all box plots the Y-axis represents each angiogenic metric and X-axis the group measured. Blue represents the DMSO control and red indicates CpdG-treated cells, length is represented in pixels. As shown, slight reductions in all angiogenetic metrics CpdG treated cells are clearly visible. N = 20 per group.

### 3.5 Lipidome of manipulated HUVECs is significantly altered

As the OSBP family has been shown to transport multiple types of lipids (59), we decided to investigate how the manipulation of ORP7 affects the lipidomic profile of HUVECs and examined the cellular cholesterol levels using Amplex Red assay. We found no significant changes in the protein normalized concentrations of total or free cholesterol levels in the CpdG-treated cells compared to the DMSO group, but an increasing tendency in the mean free cholesterol levels was seen (Figure 9).

**Figure 9.**
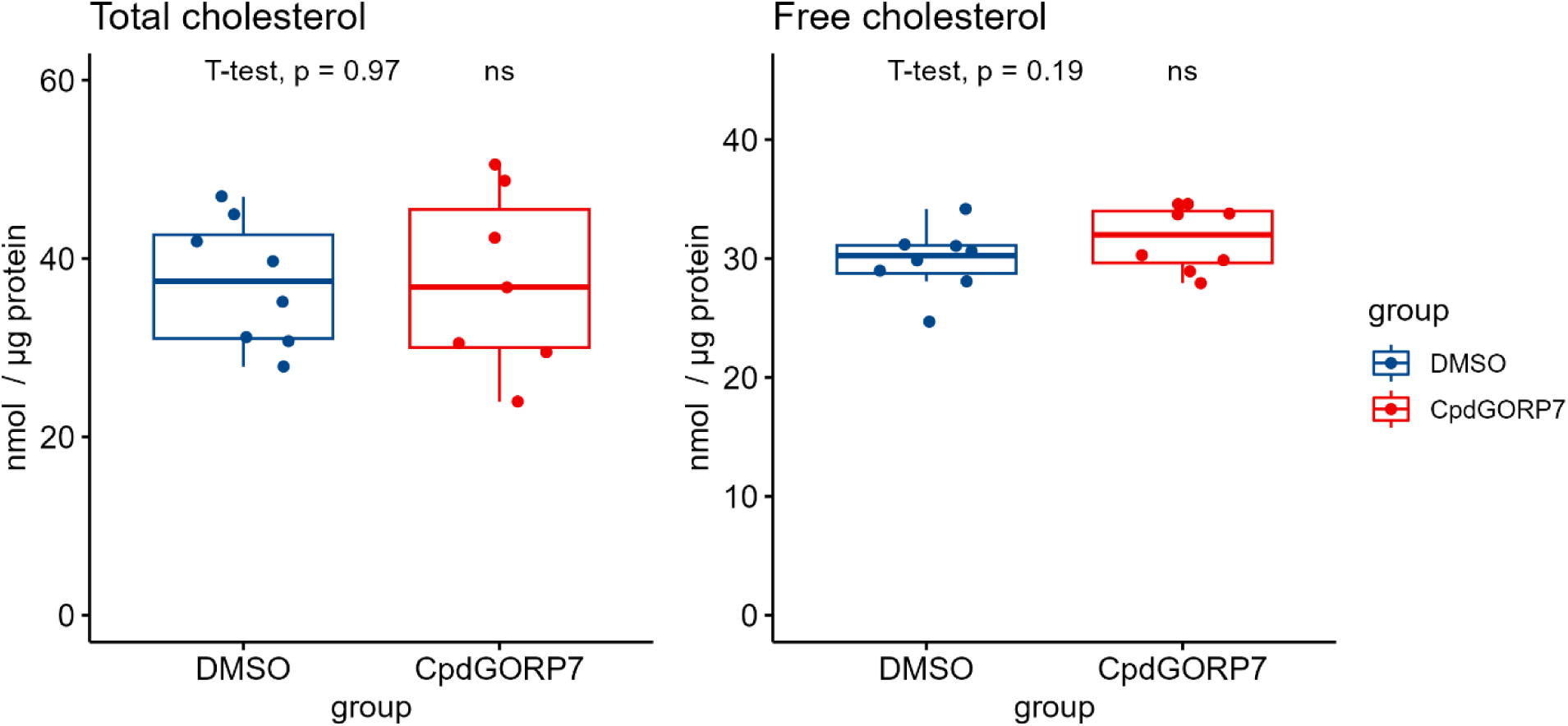
Total and free cholesterol levels in CpdG and DMSO treated cells. Box plots show on the Y-axis protein normalized cholesterol concentrations and on the X-axis each group measured. Blue represents DMSO control and red depicts CpdG-treated cells. Although no significant changes can be seen a slight increase in mean free cholesterol levels is visible. P-values were determined using Student’s T-test and N = 7 Although we did not see significant changes in the cholesterol pools of manipulated HUVECs, we also wanted to reproduce the already published increase in cholesterol efflux in CpdG-treated cells (9). To this end we performed a cholesterol efflux assay using ^3^H-labeled cholesterol and human ApoA1 or HDL as acceptors (Figure 10). These experiments showed a significant increase in ApoA1 dependent cholesterol efflux with the positive control T091317, a known LXR agonist that induces ABCA1 mediated cholesterol efflux (60), but not with CpdG. However, a significant decrease in HDL dependent cholesterol efflux was observed in CpdG-treated cells, which is mediated through ABCG1 (61) (Figure 10).

**Figure 10.**
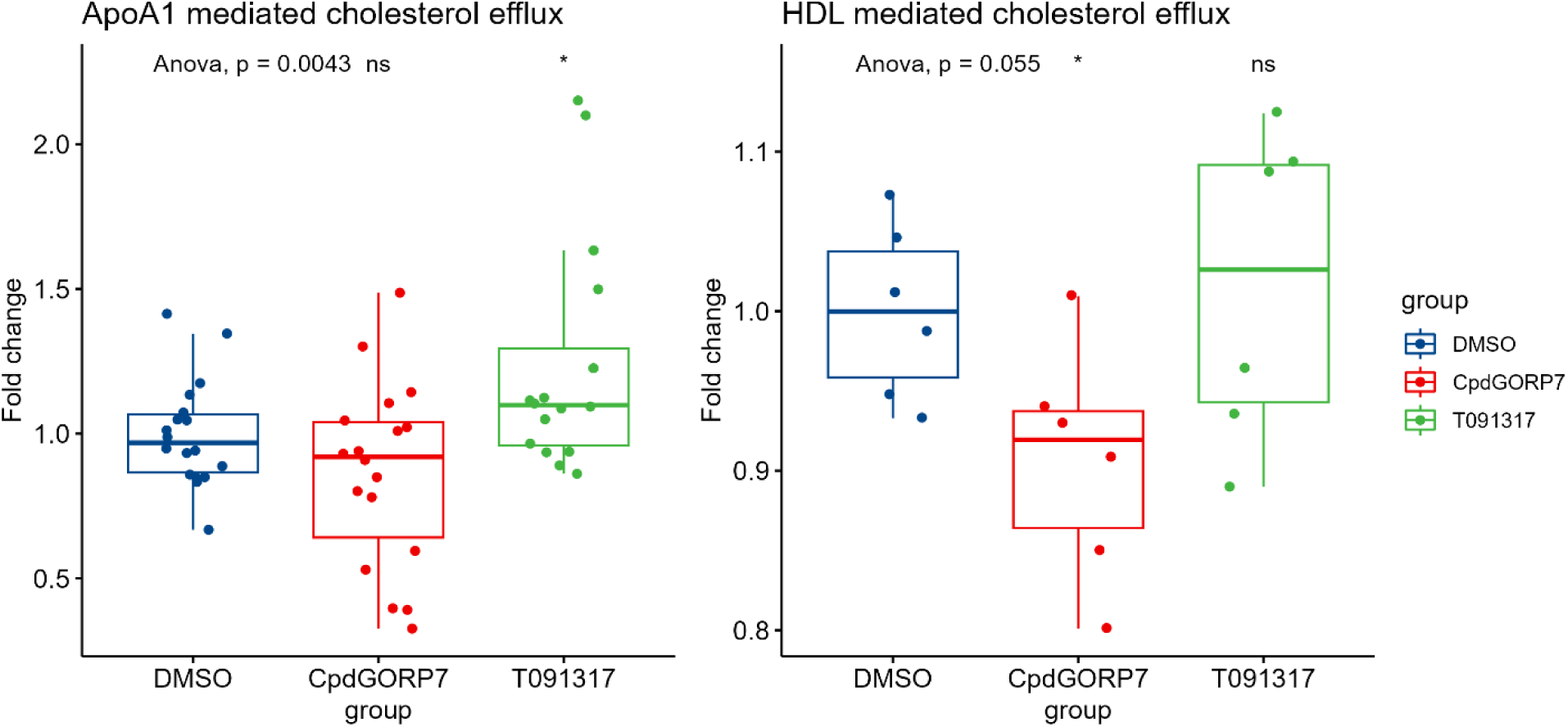
Cholesterol efflux measured in fold change in CpdG, DMOS or T091317 treated HUVECs. Box plots are shown where Y-axis represents the fold change compared to the mean of DMSO control and the X-axis each group measured. Controls are shown in blue. The left facet shows a significant increase in cholesterol efflux to ApoA1 in T091317-treated cells whereas the CpdG treatment did not show any significant increase, compared to DMSO control. On the right facet a significant decrease in efflux to HDL in the CpdG-treated cells can be seen, whereas no notable change is visible in the T091317 or DMSO group. Pairwise p-values were determined using Student’s T-test against DMSO group and groupwise ANOVA was calculated using DMSO as reference. N = 18 for ApoA1, N = 6 for HDL per group.

We next investigated how the inhibition of ORP7 affects the lipidome of HUVECs. We performed LC-MS/MS mass spectrometric analysis to get a systemwide view of the total lipidome changes in CpdG-treated cells (Figure 11). When looking at total lipid class levels, the CpdG treatment induced significant reduction of cholesteryl esters (CE) and phosphatidylserines (PS), and significant increases in ceramides (Cer), lysophosphatidylcholines (LPC) and triacylglycerols (TG) (Figure 11).

**Figure 11.**
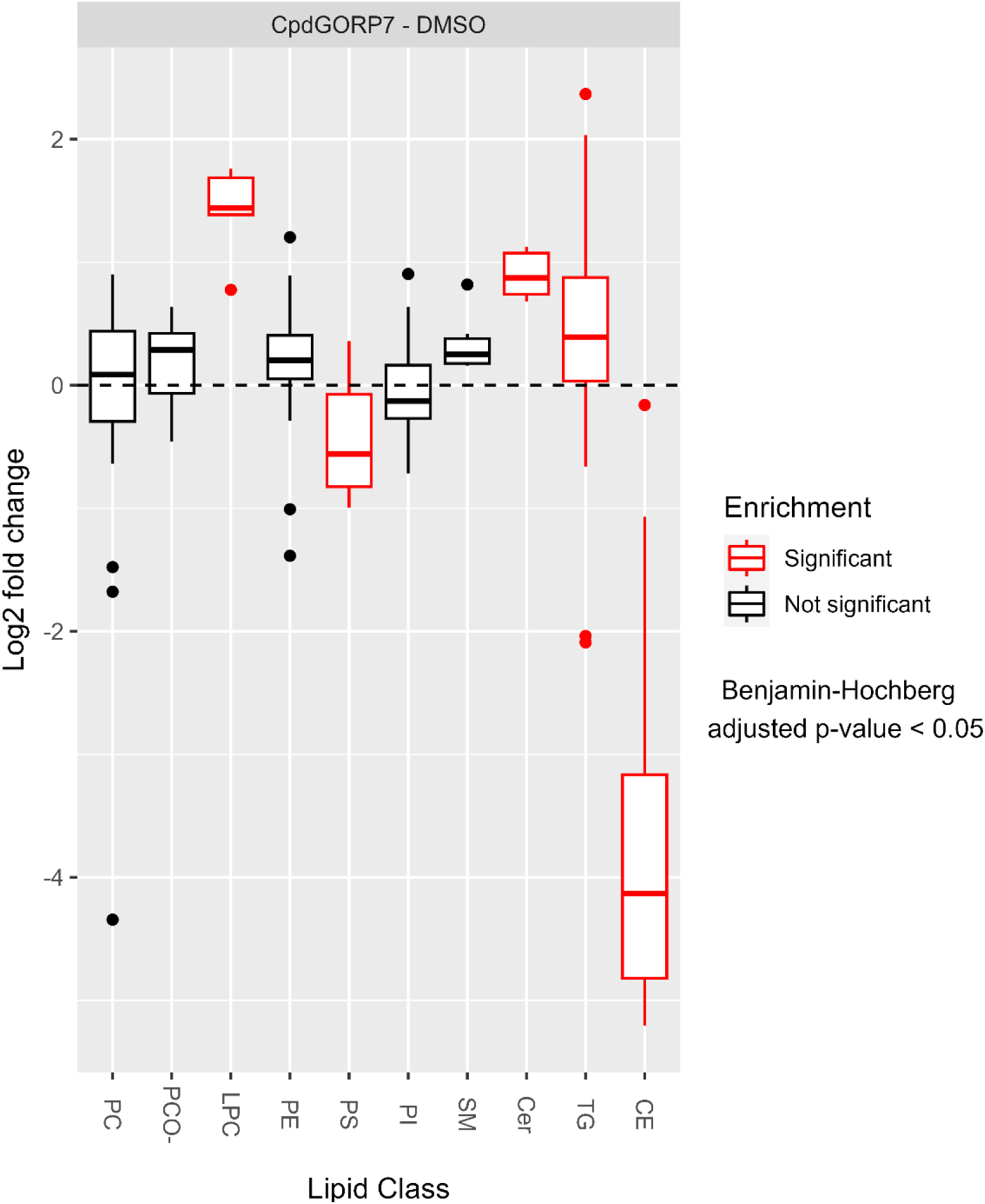
Log2 fold change distribution of different lipid classes in CpdG-treated HUVECs as compared to DMSO-treated controls. The Y-axis represents log2 fold change and X-axis each lipid class, significantly altered classes are shown in red, according to Benjamini – Hochberg adjusted p-values. The CpdG-treated HUVECs show significant changes in CE, Cer, LPC, PS and TG.

Even though the total levels of certain lipid classes had not changed significantly, there were multiple differences in the molecular species in each lipid class. Therefore, we plotted the log2 fold change of individual lipid species in each lipid class using a tile plot (Figure 12.).

**Figure 12.**
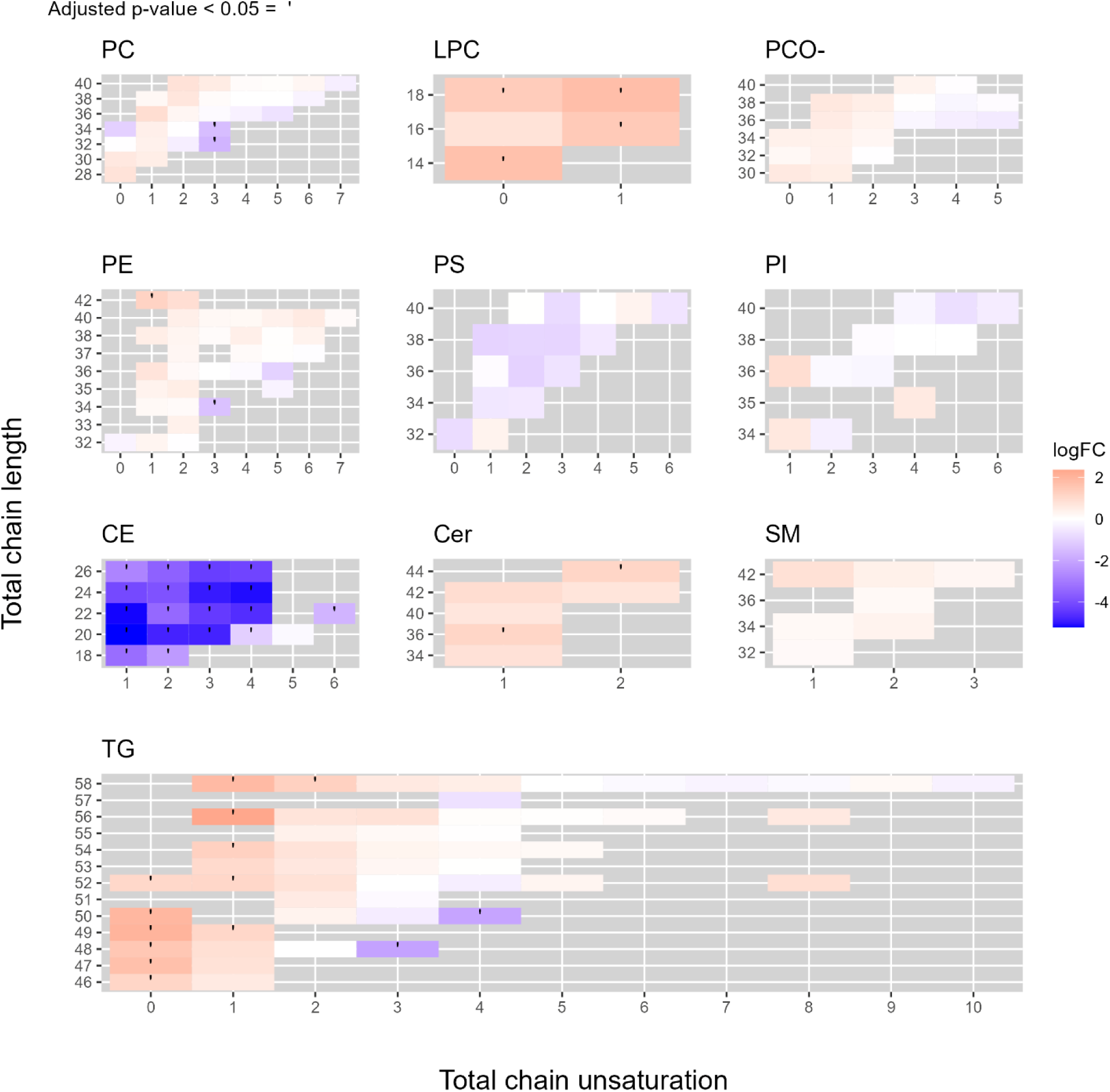
Lipid species specific changes in CpdG-treated cells as compared to DMSO-treated controls. Each facet shows a different lipid class, where the Y-axis represents the total chain length, the X-axis total chain unsaturation and each tile depicts a different lipid species. Each tile is colored according to the log2 fold change of each species, where orange represents an increase and blue a decrease, each statistically significantly altered species has been marked with a dot on the tile.

ORP7 inhibition by CpdG treatment reduced CEs and PSs, and increased LPCs and Cers across all molecular species, whereas other classes showed targeted lipid species-specific increases and decreases (Figure 12). For example, saturated and monounsaturated TGs increased upon the ORP7 inhibition, whereas a decrease was found in relatively short polyunsaturated species (TGs 48:3 and 50:4). The intermediates between those two types of TGs showed a slight increase or no change at all. Phosphatidylcholines (PCs), phosphatidylcholine-alkyls (PCO-) and phosphatidylethanolamines (PEs), exhibited a general pattern where ORP7 inhibition caused increases for most saturated and monounsaturated species (except PC 34:0). The clearest decreases due to ORP7 inhibition were found in few relatively short polyunsaturated species (significant for PC 32:3, PC 34:3 and PE 34:3), whereas the longest highly unsaturated species showed mild, non-significant responses. The responses of SM, PS and PI species to ORP7 inhibition remained non-significant; all SM species and a majority of phosphatidylserine (PS) and phosphatidylinositol (PI) species showed a decreasing tendency as compared to the control DMSO treatment.

As ORP7 inhibition by CpdG had decreased CEs drastically but simultaneously increased TGs, we investigated how microscopic occurrence of lipid droplets in the HUVECs had changed, CEs and TGs being the main ingredients of the droplets. We used BODIPY 493/503 staining to visualize and quantify lipid droplets in the CpdG- and DMSO-treated cells. Even though the nuclei normalized lipid droplet number had not changed, we observed a drop in the mean area of lipid droplets in the CpdG-treated HUVECs (Figure 13).

Pairwise p-values were determined using Student’s T-test against DMSO group and N = 13 per group.

**Figure 13.**
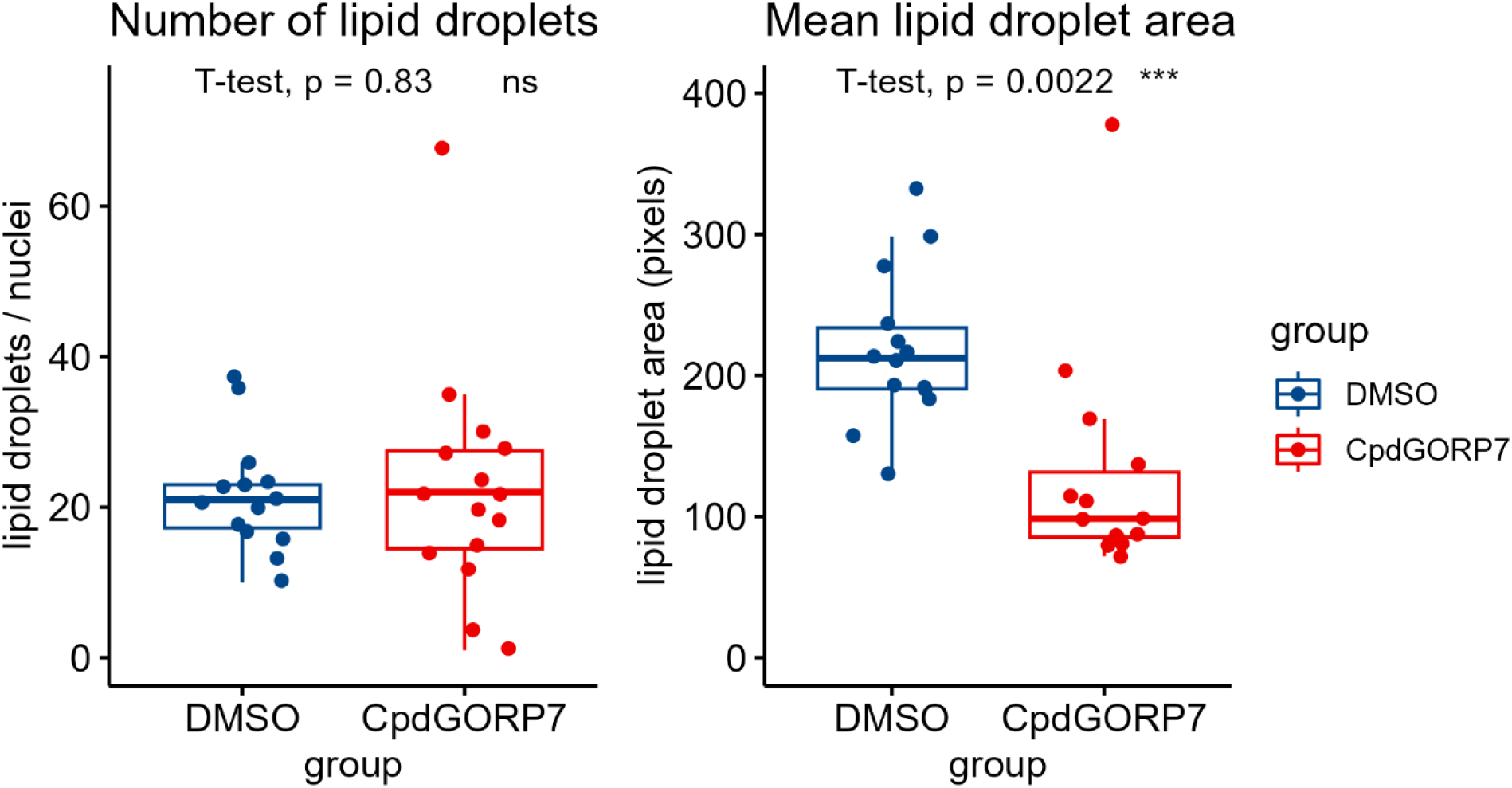
Lipid droplet metrics in measured from CpdG-treated HUVECs using BODIPY 493/503 stanning. Left panel displays mean lipid droplets per nuclei and right panel the mean lipid droplet area in pixels on the Y-axis, whereas the X-axis shows each group measured. No notable change can be seen in the number of lipid droplets, but the mean area has decreased significantly in CpdG-treated cells.

### 3.6 Proximity biotinylation interactomics shows interaction between ORP7 and basement membrane components

One of the best ways to understand the biological role of a protein is to study its interactome, which can not only reveal the biological process it is involved in, but also its subcellular location. To this end, we performed proximity biotinylation interactomics analysis by using an ORP7 with biotinylation tag as prey, which resulted in a total of 1632 proteins being captured, most of which were also captured from cells expressing the mycBirA bait. These data were then subject to the SAINT analysis, where the top 10 candidate ORP7 interaction partners were indicated (Table 4). Interestingly, these partners included multiple extracellular matrix (ECM) components: Fibronectin (FN1) Collagen Type IV Alpha 4 A 1 & 2 as well as 18A1 (COL4A1/2, COL18A1), along with Perlecan (HSPG2) and Multimerin 1 (MMRN1). Other proteins in the top 10 were RAC-alpha serine/threonine-protein kinase (AKT1) and Chaperonin Containing TCP1 subunit 7 (CCT7). Interactomics results for VAPA and VAPB, ER proteins acting as membrane anchors for several ORPs, did not produce any results from the SAINT analysis, but their unique spectrum counts were added in Table 4.

**Table 4.**
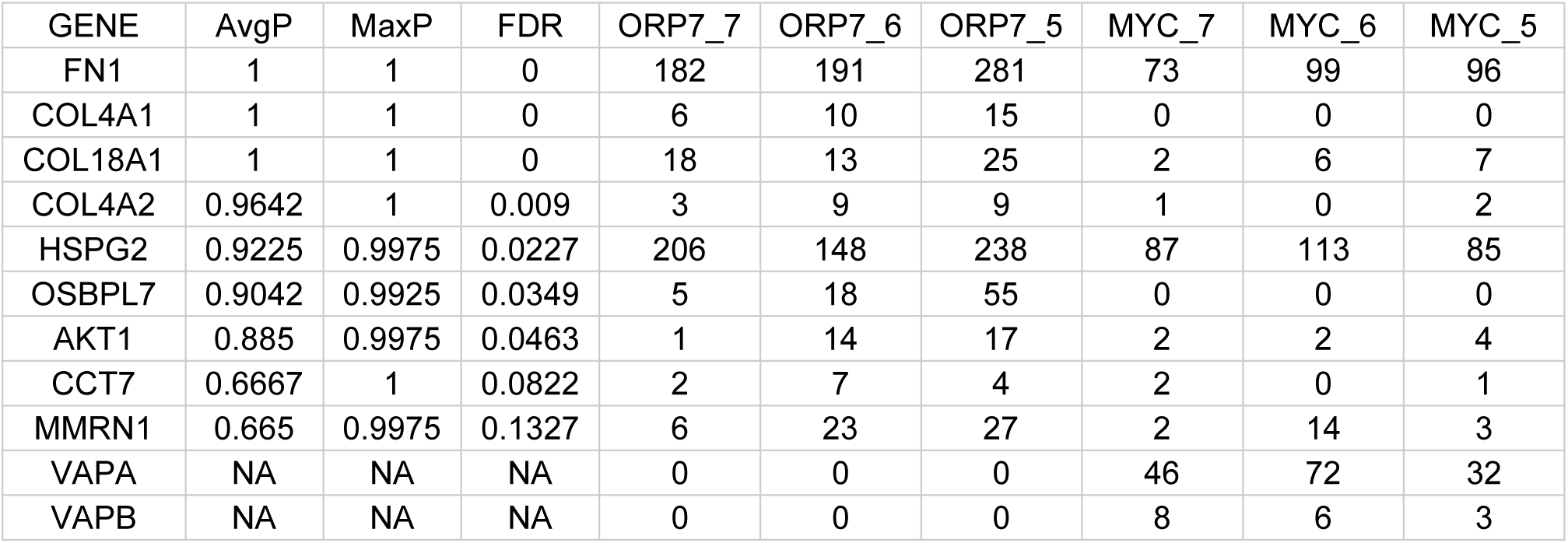
SAINT interactomics results. As statistical metrics, the average probability (AvgP), maximum probability (MaxP), and false discovery rate (FDR) are shown for the top 10 proteins predicted to interact with ORP7, as well as the unique spectrum counts in each sample on the last 6 columns.

| GENE | AvgP | MaxP | FDR | ORP7_7 | ORP7_6 | ORP7_5 | MYC_7 | MYC_6 | MYC_5 |
| --- | --- | --- | --- | --- | --- | --- | --- | --- | --- |
| FN1 | 1 | 1 | 0 | 182 | 191 | 281 | 73 | 99 | 96 |
| COL4A1 | 1 | 1 | 0 | 6 | 10 | 15 | 0 | 0 | 0 |
| COL18A1 | 1 | 1 | 0 | 18 | 13 | 25 | 2 | 6 | 7 |
| COL4A2 | 0.9642 | 1 | 0.009 | 3 | 9 | 9 | 1 | 0 | 2 |
| HSPG2 | 0.9225 | 0.9975 | 0.0227 | 206 | 148 | 238 | 87 | 113 | 85 |
| OSBPL7 | 0.9042 | 0.9925 | 0.0349 | 5 | 18 | 55 | 0 | 0 | 0 |
| AKT1 | 0.885 | 0.9975 | 0.0463 | 1 | 14 | 17 | 2 | 2 | 4 |
| CCT7 | 0.6667 | 1 | 0.0822 | 2 | 7 | 4 | 2 | 0 | 1 |
| MMRN1 | 0.665 | 0.9975 | 0.1327 | 6 | 23 | 27 | 2 | 14 | 3 |
| VAPA | NA | NA | NA | 0 | 0 | 0 | 46 | 72 | 32 |
| VAPB | NA | NA | NA | 0 | 0 | 0 | 8 | 6 | 3 |

Given the overrepresentation of ECM components, and the fact that ORP7 has been shown to localize to the cytoplasmic face of the plasma membrane (5), we suspected that the interaction between ORP7 and these ECM components could be driven by AKT1, as it is unlikely that ORP7 would come into proximity with secreted or extracellular matrix proteins given its known localization in the cytoplasmic compartment. To confirm the suggested interaction of ORP7 with AKT1, we performed co-immunoprecipitation using ProteinG coupled magnetic beads and AKT1 antibody (Figure 1). ORP7 and AKT1 were found co-precipitating from both ORP7 overexpressing and wild-type HUVECs, while the proteins were absent in precipitates pulled down with from negative control lysate incubated in the absence of the AKT1 antibody (Figure 14).

**Figure 14.**
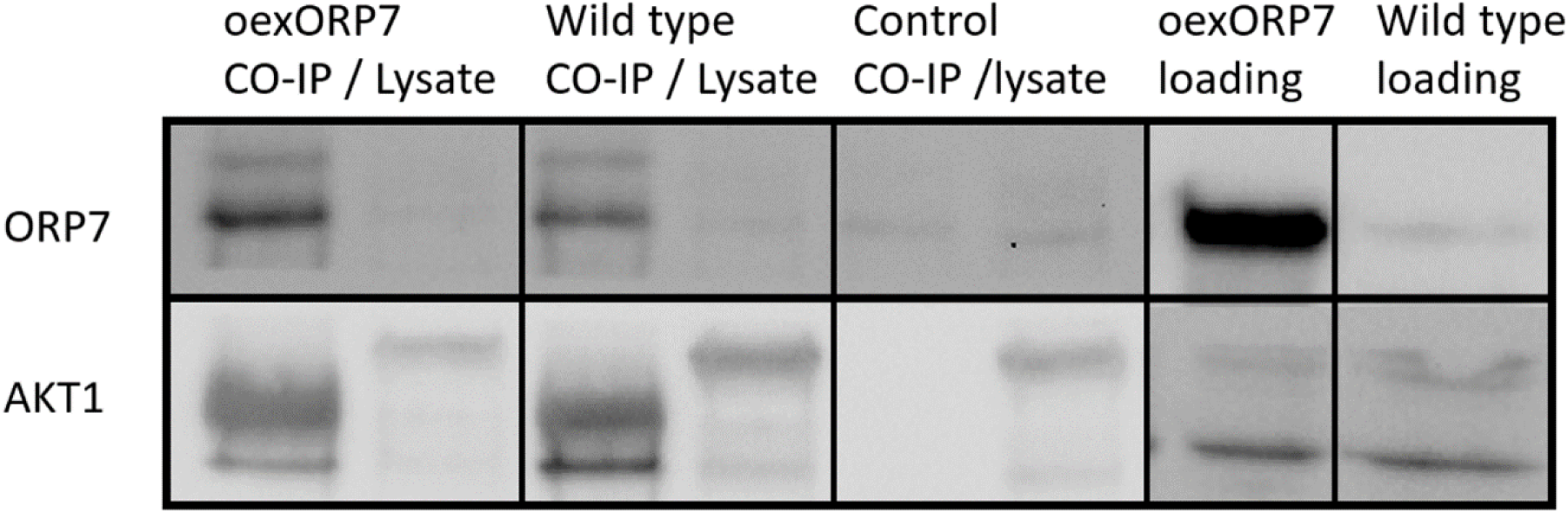
Western blotting results of co-immunoprecipitation using Protein-G coupled magnetic beads incubated with AKT1 or mouse-HRP antibody used as a negative control. Each wide box shows a different sample type, and the two lanes in each box show elution from beads (CO-IP) and lysates after incubation with beads (Lysate). The first box shows these results in oexORP7 samples, the second box shows lysates from HUVEC without any treatment or overexpression (Wild type=, and the third box oexORP7 sample lysates incubated with Protein-G coupled magnetic beads incubated with mouse HRP antibody. The rows show antibody staining for ORP7 in the top lane and AKT1 in the bottom lane.

As AKT1 has been shown to interact with and activate FAK at focal adhesions (62,63), we also examined by western blotting if overexpression of ORP7 activates either of these kinases (Figure 15).

**Figure 15.**
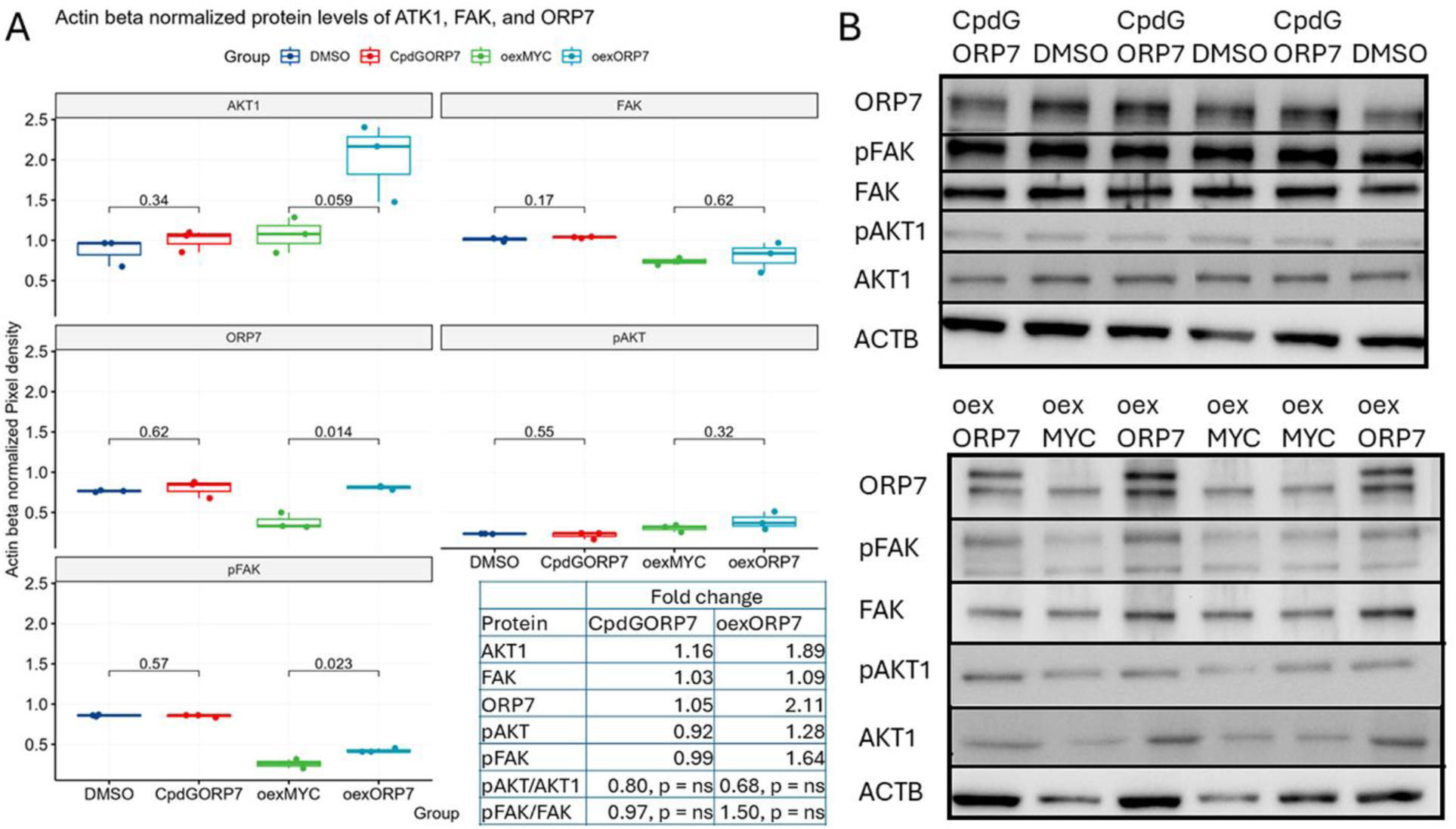
Western blotting results of both unphosphorylated and phosphorylated FAK and AKT1 as well as β-actin loading control. A) Box plots and table quantifying protein levels of AKT1, FAK and ORP7 and their fold change, where each facet shows the levels for different proteins, and pairwise t-test p-values shown above brackets as a comparison to DMSO and oexMYC in CpdG treated and ORP7 overexpressing cells, respectively. N = 3. B) Western blots where sample types are shown on top of each lane and each row shows the protein detected. The top panel shows these results for CpdG and DMSO treated cells and bottom panel for oexORP7 and oexMYC cells. Table shows a significant pAKT1/AKT1 ratio in oexORP7 cells compared to oexMYC control.

The levels of phosphorylated FAK and AKT1 (pFAK, pAKT1) as well as those of unphosphorylated AKT1 appeared to be increased in the oexORP7 cells, but only the pFAK increase was statistically significant. No change in unphosphorylated FAK could be seen. In the CpdG-treated cells, the levels of both phosphorylated AKT1 and FAK or their unphosphorylated forms showed small statistically insignificant changes. The pAKT1/total AKT1 ratio decreased significantly upon ORP7 overexpression but was not affected by ORP7 inhibition, while a statistically insignificant increasing trend in pFAK/FAK ratio was seen in the ORP7 overexpressing cells.

## 4. Discussion

As we have shown, inhibition of ORP7 leads to a plethora of complex outcomes in endothelial cells, many of which are unlikely to be a direct consequence of CpdG-inhibition of ORP7. These downstream effects ranged from the upregulation of lipid biosynthesis, oxidative stress, and inflammatory processes to reductions in cell division, proliferation, angiogenesis *in vitro* and cholesterol efflux. We suspect that these outcomes are the result of dysfunctional lipid traffic from the ER to the plasma membrane, mitochondria or autophagosomes, as will be discussed in the sections to come.

### 4.1 Systemwide omics results point toward changes in inflammatory processes, lipid metabolism, and oxidative stress

Gene set analysis of our transcriptomic data showed decreases in mRNAs related to cell cycle, cell division and nucleotide synthesis, as well as increases in genes related to inflammation, oxidative stress and lipid metabolism. Lipidomic analysis, on the other hand, exhibited increases in lipid classes known to increase inflammation and endothelial cell dysfunction, such as ceramides (64,65), TGs (66,67) and LPCs (68).

#### Inflammation

The relative upregulation of the inflammatory process in the transcriptomics data can most likely be attributed to genes encoding pro-inflammatory proteins such VCAM1, ICAM1 and SELE, which are shown to be elevated in the early stages of atherosclerosis (41). A vital anti-inflammatory molecule also affected is the transcription factor *KLF4*, which controls the expression of multiple inflammation-associated genes (69), of which *CH25H* is a pivotal example in our transcriptomics data set (70). Another anti-inflammatory gene upregulated upon CpdG treatment is *STC1*, which has been shown to mediate inflammation by inhibiting the tissue infiltration of leukocytes (50). Moreover, downregulation can be observed in pro-inflammatory genes such as *THY1*, *IL1RL1*, and *IL7R* (54,55). Other evidence suggesting that CpdG-treated HUVECs show inflammatory activation is the induction of genes of arachidonic acid (AA)-derived lipid mediator synthesis i.e., *PTGS2*, *SLOC2A1* and *CYP1A1.* Lipid mediators are auto- and paracrine signals produced from membrane phospholipids (Underwood et al., 1998; Yamashita et al., 2005, 2009,Lu *et al.*, 1996; Chan *et al.*, 2002; Mesaros, Lee and Blair, 2010), especially from the hydrolysis of PC that produces LPC species, which had decreased and increased respectively, in our lipidomic data. *PTGS2* may convert phospholipid-derived AA to pro-inflammatory prostanoids, and *CYP1A1* may convert AA to epoxyeicosatrienoic acids and hydroxyeicosatetraenoic acids, having both anti- and proinflammatory properties. It is possible that the CpdG-treated cells are trying to produce more epoxyeicosatrienoic and hydroxyeicosatetraenoic acids by upregulating *CYP1A1* (71,72), as its log2 fold change is almost three times that of *PTGS1/2*. Since we did not see significant changes in lipoxygenase genes (*ALOX*), we suspect that neither leukotriene nor lipoxin synthesis had changed (73). The observed increase in Cers and LPCs, and upregulation of biosynthetic genes of AA-derived lipid mediators together suggest that inflammatory signaling is activated in the CpdG-treated cells.

As the oxidized AAs, 25-hydroxycholesterol (25-HC) has diverse functions, and we saw a relatively large increase in *CH25H* expression, which suggests that cholesterol is being turned into oxysterols (74). This observation is supported by the increase in *CYP27A1* expression as increase in CYP27A1 protein levels should increase the synthesis of 27-hydroxycholesterol from cholesterol (75). We did not see significant changes in the cholesterol biosynthesis pathway genes but did see an increase in liver X receptor (LXR) expression activating cholesterol efflux by ABCA1. Since cholesterol biosynthesis genes were not changed and if ORP7 traffics cholesterol from the ER, it is likely that the CpdG-treated cells could have turned excess FC into oxysterols (76); Thus, excess FC at the ER may not be the main culprit for increased inflammatory gene expression. Instead, inflammation could be the result of excess oxysterol products derived from FC.

#### Raft accumulation of accessible cholesterol

Why CpdG-treated cells are consuming CEs from lipid droplets, as is evident from the reduced mean area of lipid droplets and drastic reductions in all CE species, is unknown. CEs could be used to balance the plasma membrane level of chemically available cholesterol, which might be reduced given the fact that ABCG1 mediated cholesterol efflux was reduced, and no change ABCA1 mediated cholesterol efflux was evident. We also observed increases in the expression of *NPC1* and -*2*, which traffic FC from lysosomes to the PM (77). This could be seen as compensatory mechanisms to add more FC to the PM. The increase in *CH25H* expression is likely the result of overexpression of *KLF4* (70), but the underlying reason remains unclear.

Increased 25-HC levels should initiate synthesis of CEs (Nguyen et al., 2023). Thus, why the CpdG-treated cells show such drastic reductions in CEs and no notable induction of cholesterol synthesis genes is puzzling. One possibility could be that the relatively short 24-hour incubation time with CpdG has not been sufficient to induce CE synthesis or, what we think is a more likely explanation, that FC is accumulating to some membrane compartment or domain where it is inaccessible for CE synthesis, as we did not detect any significant change in total cholesterol levels. Decrease in ABCG1 mediated cholesterol efflux seems peculiar considering that the reduction in CEs should make more FC available for efflux. The increase in Cers and reduction of cholesterol efflux could suggest that FC is accumulating in raft domains (79)where it is unavailable for both efflux and other cellular functions. The increase in LPC could also be the result of Cer and FC accumulation in raft domains, as it can make the membrane environment more hospitable for rigid raft domains (80). Whether or not intracellular levels of 25-OHC have increased in the CpdG-treated cells is not known, but if this were the case, the consequences would not be limited to cholesterol synthesis alone, as 25-HC can also bind to the oxysterol binding pocket of multiple ORPs (81) and therefore competes with FC for ORP-mediated traffic.

#### Oxidative stress response and possible association of ORP7 with mitochondria associated membrane (MAM)

The observed reductions in mRNAs related to cell cycle, cell proliferation and nucleotide synthesis are a puzzling phenomenon. We suspect that this reduction could be due to mitochondrial dysfunction. Given that transcriptomics data showed increased NES in lipid metabolism and oxidation related gene sets, such as the oxidative stress response, the occurrence of increased oxidative stress is likely. Oxidative stress can result in the release of cytochrome C from mitochondrial cardiolipin (82), which in turn downregulates cellular division and proliferation through caspase activation (83). Some evidence pointing toward this conclusion is present in our transcriptomics data set as, for example, the NF-κB/AP-1 induced apoptosis pathway showed a positive NES. GABARAPL2 and LC3B have been shown to be necessary for mito- and autophagy (84–86) and have high affinities for membranes with cardiolipin and Cers (87), such as those seen at both mitochondrial outer and inner membranes or MAM (88,89). MAM lipid raft-like domains are also necessary for autophagy (90), which suggests that GABARAPL2 and LC3B could be associated with these raft-like domains as MAM is enriched with cardiolipin (88). It has also been shown that ceramides are necessary for mitophagy, especially C-18 ceramide (Cer 18:1;O2/18:0, Cer 36:1 in Figure 12) (Sentelle et al., 2012; Jiang and Ogretmen, 2013). We would like to speculate that Cers and FC may be accumulating in both PM and MAM lipid raft like domains, since Cers and cholesterol are known to be enriched at both the PM and MAM, and MAM raft-like domains are needed for autophagy (88,90). Therefore, this speculated increase of Cers and FC at MAM and a subsequent increase in MAM raft-like domains could be a marker for increased autophagic potential. Given that OPR7 has been shown to interact with both GABARAPL2 and LC3B independently, and both proteins interact with each other, we suspect that ORP7 may play a role in MCSs between MAM, mitochondria or autophagosomes. ORP7 could also play a more autophagy focused role at MAM, and instead its main traffic targets could be the ER and PM as it has been shown to localize to both intracellular structures (5). Our transcriptomic data showed inconsistent patterns in autophagy related genes (Table 3), but we did see an increase in LC3A/B (MAP1LC3A/B) and GABARAPL1, but not in GABARAPL2. Which lipids ORP7 might transfer at these MCSs still remains elusive, but modulating cholesterol concentration there would be likely as several of the proteins in the ORP family have been shown to traffic cholesterol (Wang, Ma, Qi, Dong, Du, Rae, Wang, W.-F. Wu, et al., 2019; Koponen et al., 2020; Nakatsu and Kawasaki, 2021).

### 4.2 Interaction of ORP7 with AKT1

As was evident from our interactomics data, we were unable to confirm the ORP7-VAPA or - VAPB interaction. This is quite puzzling as ORP7 does contain a FFAT motif, and the interaction between VAPA and ORP7 has been confirmed with GST-pull down (95), BIFC (96) and high throughput methods (97). We were also unable to confirm the interaction between ORP7 and GABARAPL2 or LC3B through proximity biotinylation interactomics, as no unique spectrum counts for these proteins were shown in either oexORP7 or oexMYC control samples. The reasons for the lack of these interaction partners in our dataset could be multiplex. It is unlikely that the reason is low protein concentrations in our pull-down samples, as endothelial cells have 6 to 7 times higher expression of GAPARAPL2 (231 nTPM), VAPA (197.5 nTPM), and LC3B (176 nTMP) as compared to AKT1 (31.6 nTPM), according to Human Protein Atlas (98). Other possibilities for the lack of these interactions in our dataset could be that the mycBirA epitope of the ORP7 construct does not come to sufficient proximity to these proteins. It is also possible that ORP7 is compartmentalized in a differential manner in endothelial cells compared to the cell models used in the aforementioned studies, and therefore ORP7 does not come into proximity with these proteins. Another possibility is that the pull-down conditions used were not suitable for GABARAPL2 or LC3B. As we alluded to in the results section, we suspect that the interaction between the ECM components and ORP7 could be driven through the confirmed interaction between AKT1, as AKT1 has been shown to interact and regulate focal adhesions (63). If the interaction between AKT1, ORP7 and focal adhesions is “sticky” enough, excess biotin from the cells can biotinylate ECM components when cells are lysed during sample preparation, as ORP7 is not known to be secreted outside the cell. Moreover, the overexpression of ORP7 did moderately increase the phosphorylation of FAK, but the underlying mechanism remains unknown. However, the impact of ORP7 on the activity of AKT1 could provide a hypothetical explanation for this. Since COL4A1, COL4A2, HSPG2 and COL18A1 are all endothelial basement membrane components (99), validating the direct interaction of ORP7 with these components could confirm whether or not ORP7 specifically binds to the endothelial basement membrane or if this interaction is driven by AKT1. The interaction between ORP7 and AKT1 could also take place at MAMs or autophagosomes as AKT1 localizes also to microtubules according to Human Protein Atlas (98). Of note, ORP7 would not be the only ORP that interacts with AKT1 as also ORP2 and ORP9 are shown to bind to AKT1 (100–102).

### 4.3. Reduced angiogenic capacity as a result of systemic changes

Results of the angiogenesis assay clearly revealed an angiogenic defect as the CpdG-treated cells exhibited reductions in all angiogenic metrics measured. Our transcriptomic data set included very little evidence pointing towards a compensation for the reduced angiogenic potential, as only VCAM1, SELE and ICAM1 were upregulated, which have been shown to promote angiogenesis (40,103). Although not shown in human endothelial cells, increase in PTGS2 activity has been shown to upregulate the expression of VEGF and therefore increase angiogenesis in tumors (104,105). Our data revealed a slight increase in both *VEGFA* and *VEGFB*, and a slight decrease in *FLT1*, which could point toward a similar mechanism in endothelial cells. The observed angiogenic defect can be the result of an unfavorable lipid composition, such as the reduction in FC available at the PM, which has been shown to reduce angiogenesis (106). This would make sense if FC and Cer are accumulating at MAM raft like domains instead of the PM, as it has been shown that PM cholesterol and sphingolipid rich lipid rafts are vital for normal angiogenesis (107,108). Other reasons for the reduction in angiogenic capacity could be the downregulation cell cycle and cell division genes by cytochrome c release and activation of caspases, which could hamper angiogenic tube formation *in vitro*.

## 5. Conclusions

This study has shown that ORP7 inhibition leads to a systematic increase in inflammation due to an increase in pro-inflammatory lipid classes such as Cers and LPCs, and to the upregulation of pro-inflammatory genes such as *VCAM1*, *ICAM1* and *SELE*. Our data also revealed that endothelial cells upregulate genes responsible for the synthesis of oxysterols, epoxyeicosatrienoic and hydroxyeicosatetraenoic acids, as well as prostanoids, putatively to regulate inflammation. Inhibition of ORP7 also leads to a decrease in both ABCG1 mediated cholesterol efflux as well as angiogenesis but has no effect on ABCA1 mediated cholesterol efflux in endothelial cells. We were also able to prove an interaction between ORP7 and AKT1 through two independent methods. This study represents the first of its kind to characterize the results of ORP7 inhibition and overexpression in a comprehensive manner in any cell type. We believe this study will serve as the basis for a larger body of work on ORP7 functions in endothelial cells as well as in other cell and animal models. More work is needed to study whether and how ORP7 inhibition affects the levels of oxysterols and prostanoids along with inflammation, as well as in the formation of rafts and raft-like domains. Moreover, it remains to be studied whether and how ORP7 manipulations may induce mitochondrial dysfunction due to oxidative stress. The lipid transport functions of ORP7 also remain unknown, but our work suggests that ORP7 could play a role in autophagosome formation at MAMs, for example by trafficking cholesterol between ER, mitochondria and autophagosomes.

## Supporting information

Supplementary text and images

## Declarations

### Ethics approval and consent to participate

Not applicable

### Consent for publication

Not applicable.

### Availability of data and materials

Data has been added to Open Science Framework and can be found from the following link https://osf.io/jy5vu/. All omics data has been uploaded to Gene expression omnibus under the accession number GSE261689. Any missing data or materials are made available upon reasonable request.

### Author contributions

**JHT** was involved in Conceptualization, formal analysis, validation, investigation, writing the original draft, visualization, funding acquisition. **MH** participated in formal analysis, investigation, review, and editing. **HR** took part in formal analysis and investigation. **RK** contributed to conceptualization, review, and editing. **VMO** partook in conceptualization, review, editing, supervision, project administration and funding acquisition.

### Footnotes

Limitations of the study are related to the compound CpdG, which has been shown to display a low affinity for cannabinoid receptor 1 (CNR1) (9). Therefore, CpdG treatment may in principle have off target effects driven through CNR1. However, our transcriptomics analysis of CpdG- or DMSO-treated HUVECs detected no transcripts for CNR1 or CNR2.

## Acknowledgements

Ms. Riikka Kosonen is acknowledged for her expert technical assistance. Human HDL was a kind gift from Adjunct professor Matti Jauhiainen (Minerva Foundation Institute for Medical Research, Helsinki).

