## Supplementary text and images for "Functional omics of ORP7 in primary endothelial cells"

Supplementary File 1

As we explained in the main article, we decided to keep most of the results we obtained for over expression samples separate from the inhibitor treated cell results. We will use the same numbering scheme for subsections and figures in order to keep cross-referencing as easy as possible. Therefore, there might not only be gaps in the subsection numbering, but similar gaps can also appear in figure numbering.

3.2. Quality control of over expression treatment, and omics results.

We also performed PCA on oex-cells omics data and these results are shown in Figure S5. Transcriptomics data show clear separation along the largest dimensions, whereas more separation is evident along the second largest dimension. Lipidomics data shows more random spread along both dimensions, even though the group centers for each group separate nicely along the diagonal, the fact that oexMYC_3 and oexORP7_4 cluster closer together compared to the other samples from their group, clearly shows that there is less variation between the groups in our lipidomics experiment.

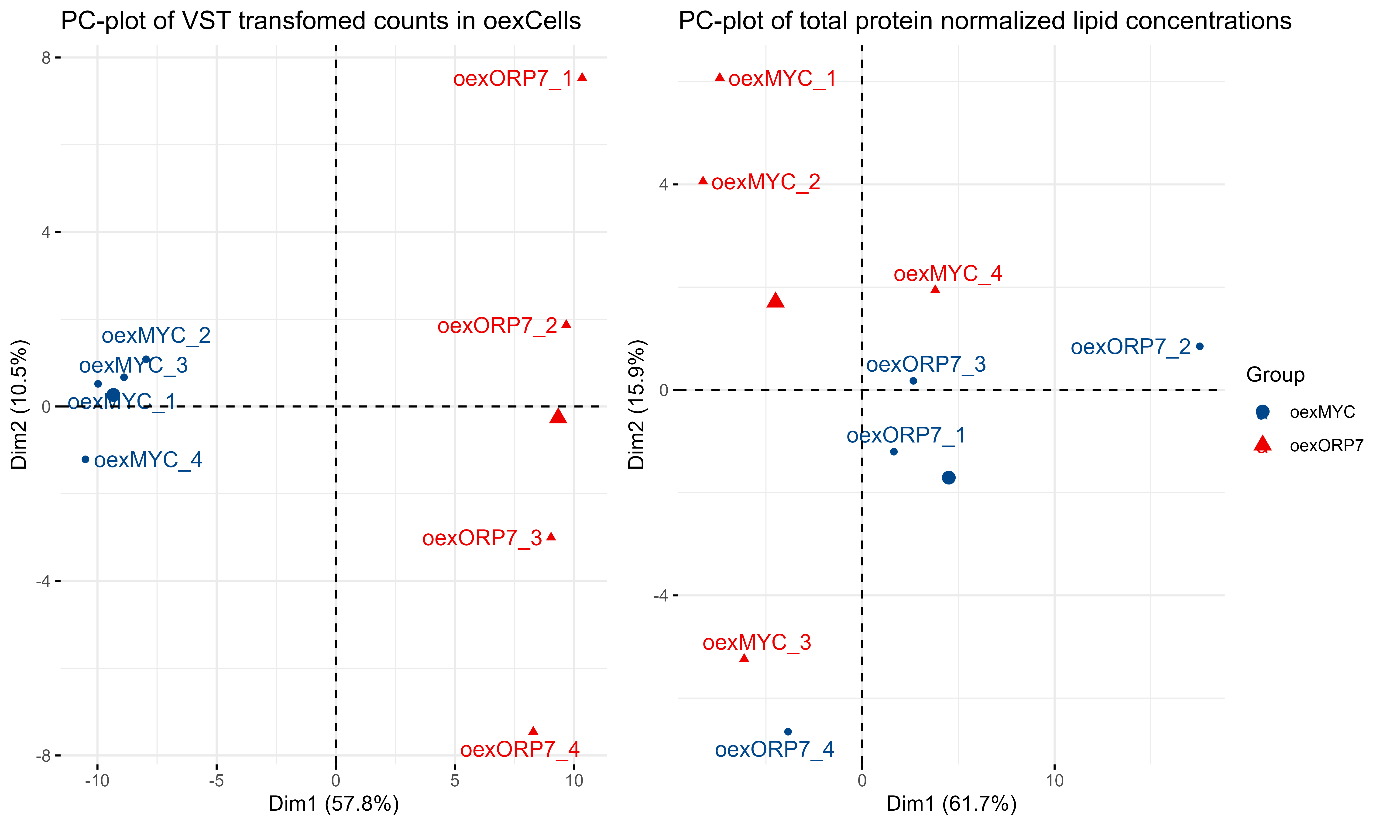

Figure S5. PC-plots of oex-cell omics results, all plots control is show in blue and manipulation show in red. Each axsis shows the two largest components. PC-plots of VST transformed transcript counts and total protein normalized lipid concentration in oex HUVECs are shown, where samples show clear separation along the largest dimension in transcriptomics samples, but a more random separation of lipidomics samples is seen.

3.3 Transcriptomics reveals multifaceted changes in inhibitor treated HUVECs.

We performed gene set analysis on oex-cell samples, but the results showed relatively little significant changes in any gene sets, suggesting that expression changes are somewhat random. For those who are not well initiated in gene set analysis, a simplified explanation of how gene sets are considered significant is as follows: Significance of a gene set is based on how genes within a gene set cluster in a, for example, log2 fold change ranked list of genes. I.e. if genes in a gene set cluster more towards top, middle or bottom of the list the gene set is considered significant, if genes are clustered randomly in the list, the set is not significant. In Figure S6 a dot plot of enriched gene sets from Wikipathways is shown, which demonstrates on the X-axis the normalized enrichment score (NES) of each gene set shown on the Y-axis. The color of each dot represents the Benjamini Hochberg adjusted p-value for each gene set. Figure S6, shows some similarities to the results in treated cell although the amount of significantly altered pathways is much lower. For example, on the left facet pathways related to cancer are shown and on the right facet pathways related to cholesterol synthesis are visible. Most notably inflammation related sets are shown in the left facet as well as cancer related sets in the right facet which is opposed to what we showed in treated cells. Other databases show very few or no significantly altered pathways and these figures are shown in Figures S22-S29, which depict similar dot plots to Figure S6.

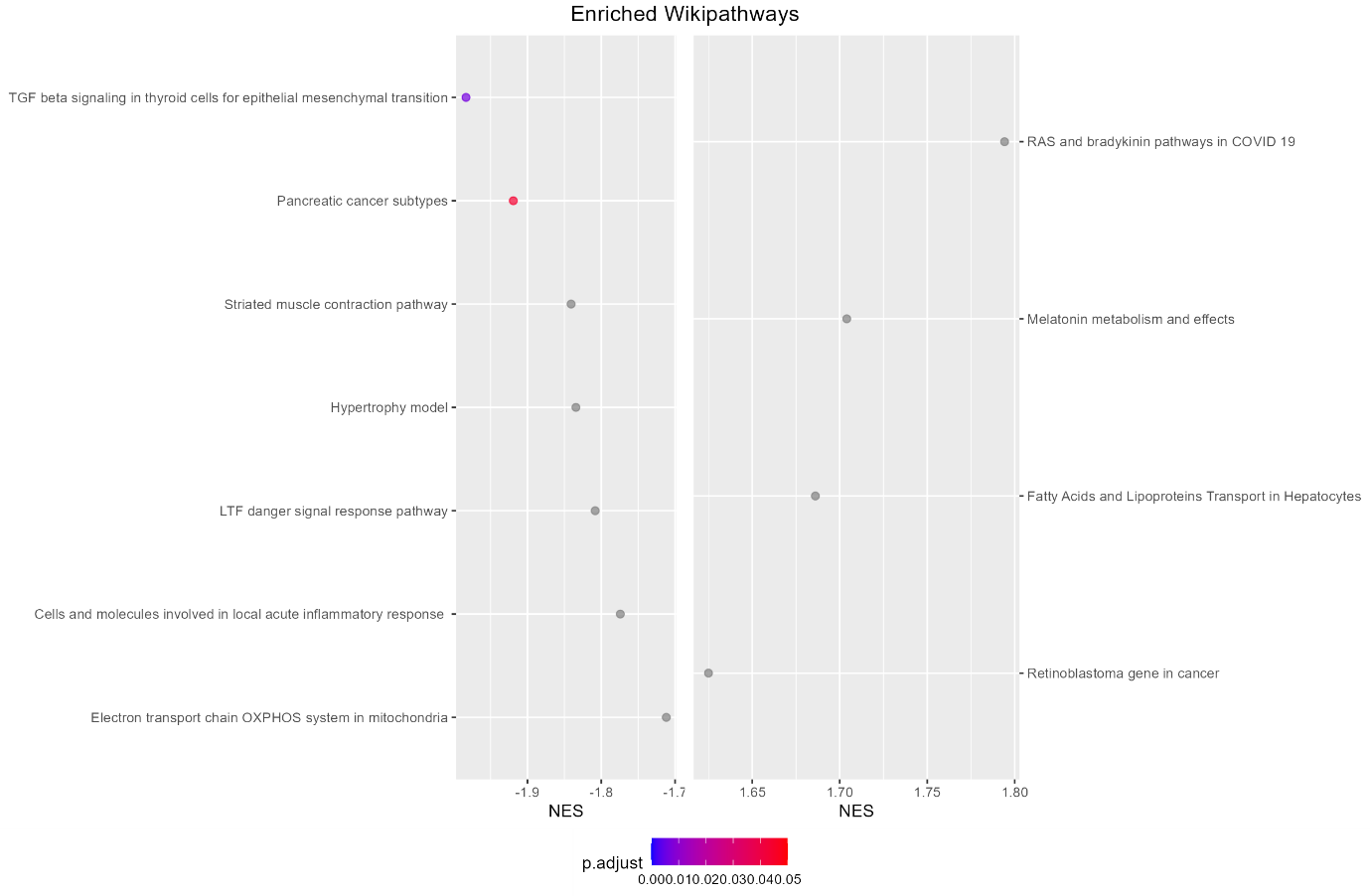

Figure S6. Dot plot of enriched Wikipathways in oex-cells. Y-axis represents enriched gene sets, and X-axis the normalized enrichment score for each gene set, where a negative value indicates more downregulated genes and a positive value more upregulated genes. The color of each dot represents the Benjamini Hochberg adjusted p-value for each gene set.

As we mentioned in the main article, gene set analysis is not the end all be all functional analysis, it is sometimes very insightful to also investigate the individual gene changes. The most notable difference between oex-cells and treated cells is that genes in oex-cells had a larger log2 fold change in both directions. Although these changes seem random based on gene set analysis, some insight could still be made. These changes are exhibited in Figure S7.

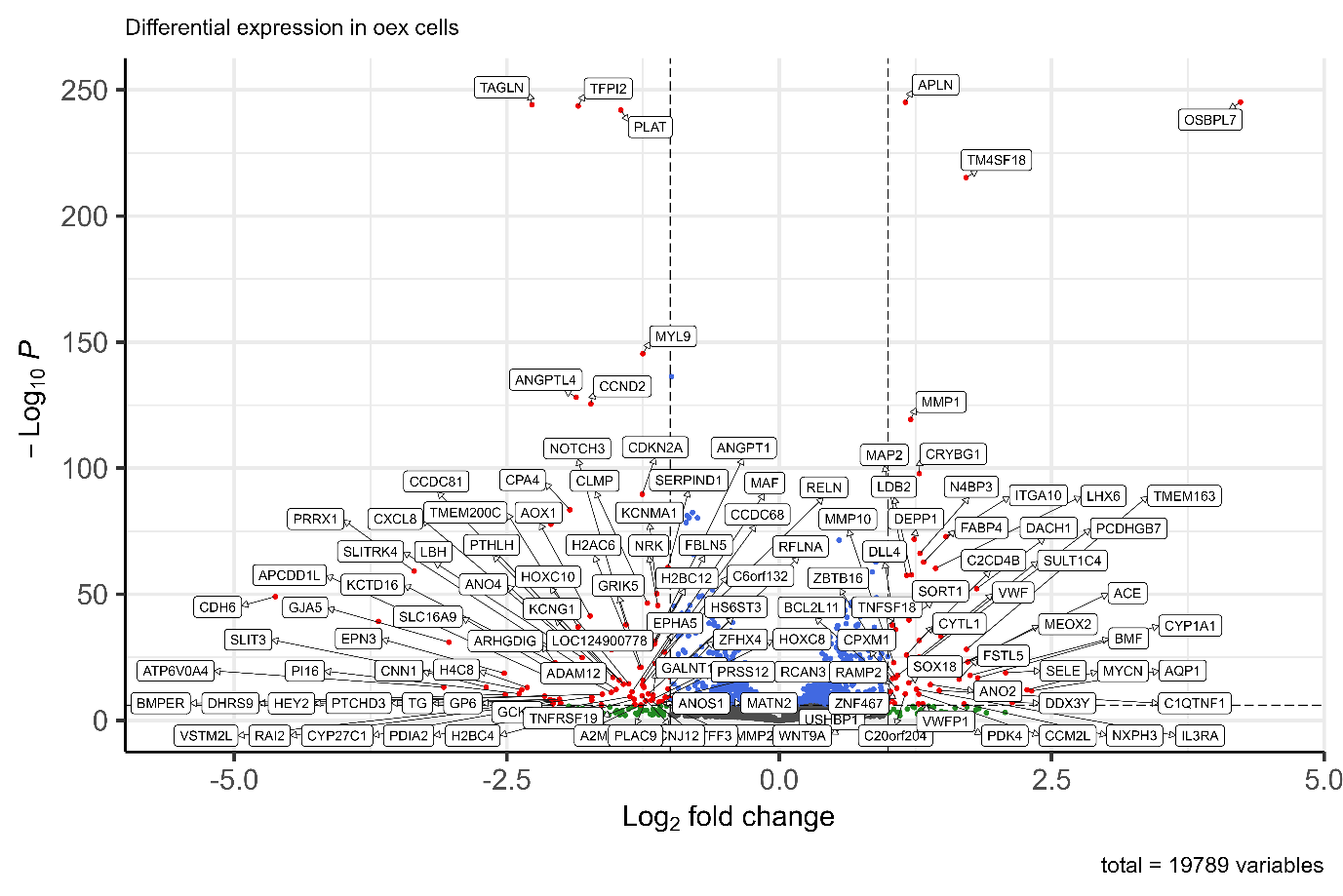

Figure S7. Volcano plot of differential expression results in CpdG treated cells. X-axis represents log2 fold change and Y-axis -log10 adjusted p-value. Labeled red dots show genes that have an absolute log2 fold change of 1 or higher and are significantly altered, blue dots have an absolute log2 fold change which is lower than 1 and are significantly altered, grey dots are genes that have not been significantly altered.

We can see that OSBPL7 has had the larger positive fold change and with a large significance which as a good quality control metric for our overexpression construct in general. There are some similarities between the inhibited cells and oex-cells in the overexpressed genes, mainly: CYP1A1 ITGA10, SELE and SERPIND1 at the downregulated genes.

3.4 Manipulated HUVECs have reduced angiogenic capacity.

Oex-cells were plated on a matrix which induces angiogenesis in HUVECs, to measure how the over expression of ORP7 affects angiogenesis. These results are shown in Figure S8, where multiple angiogenesis metrics have been plotted in oex-cells. All Y-axes represent different angiogenic metrics whereas all x-axes show each group measured. In all statistics calculated oexMYC cells were the reference group.

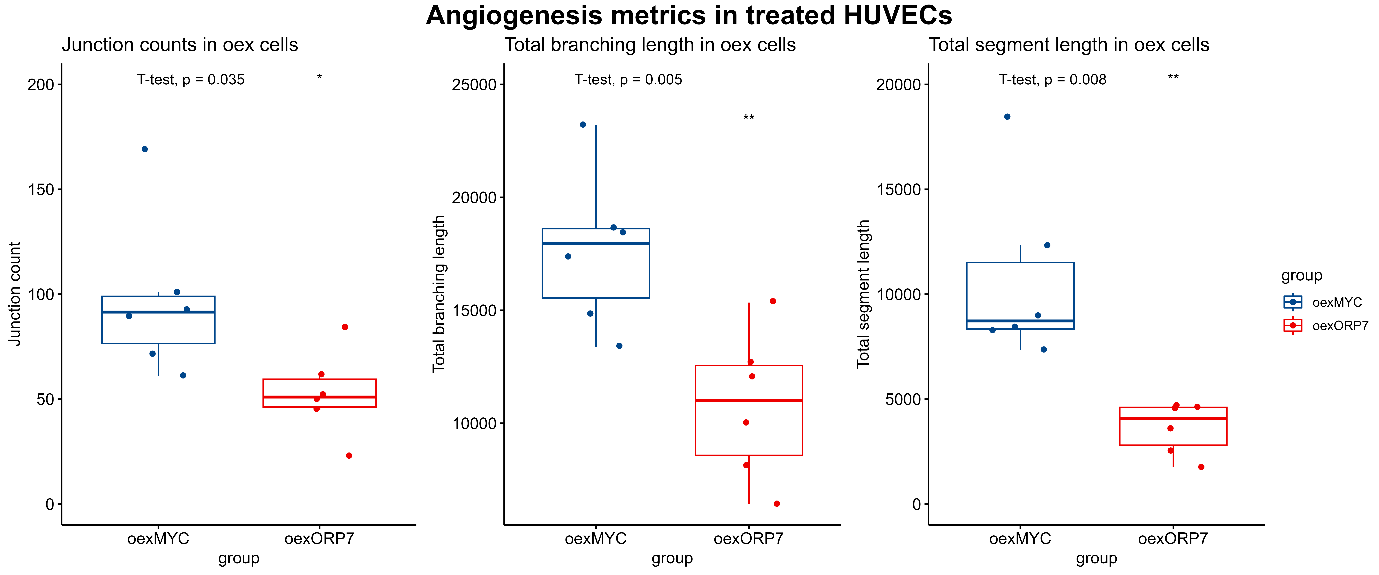

Figure S8. Bar plots of angiogenesis metrics. In all plots the Y-axis represents the each angiogenic metric and X-axis the group measured. Blue represents the oexMYC cell control and red indicates inhibitor treated. As shown, slight reductions in all angiogenetic metrics oexcells are clearly visible.

As exhibited in Figure S8. overexpression of ORP7 reduced angiogenetic capacity in all metrics measured and a similar manner to inhibition of ORP7 but does not block it entirely.

3.5 Lipidome of manipulated HUVECs is significantly altered.

We also measured similar lipidomic metrics from overexpressing cells as from treated cells. In contrast to the treated cells we were able to detect a small reduction in free cholesterol levels in oex-cells, but otherwise these results are the same.

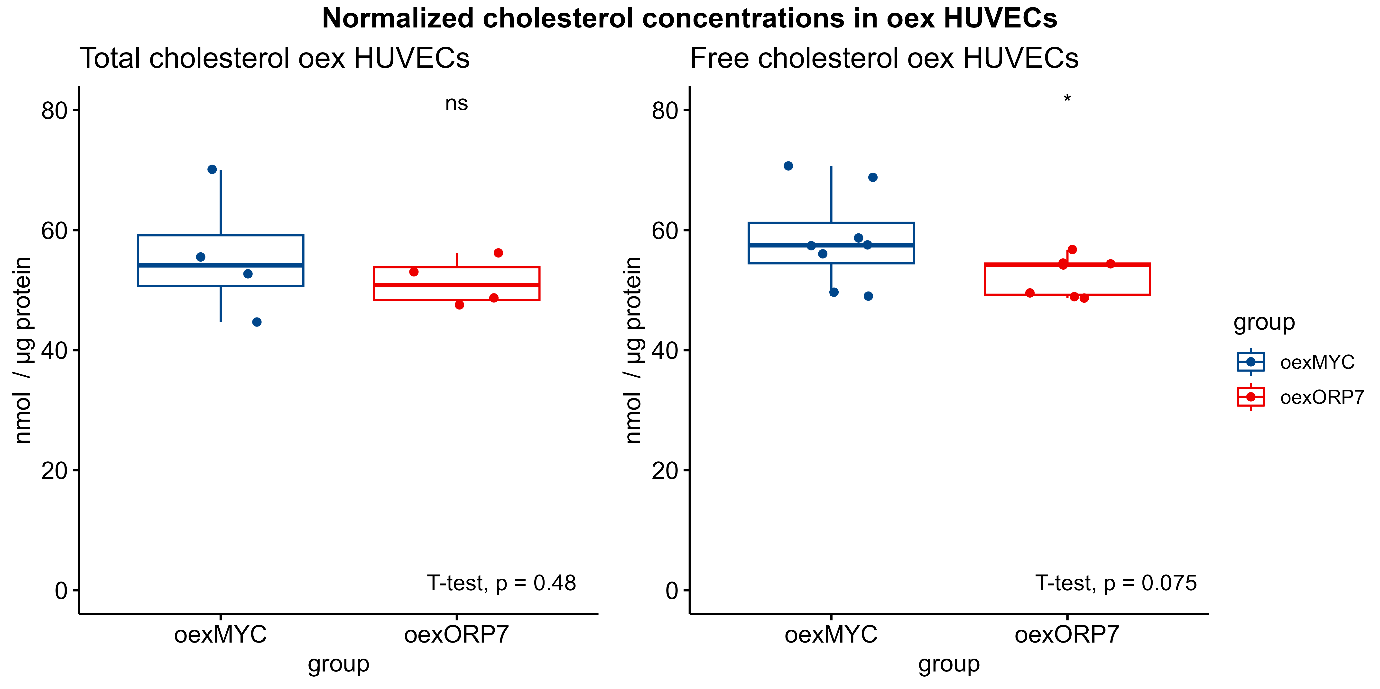

Figure S9. Box plots of total and free cholesterol levels, where the Y-axis represents protein normalized cholesterol concentrations and X-axis each group measured. Blue color represents the oexMYC control and red depicts oex-cells. A significant but slight decrease in mean free cholesterol levels is visible. P-values were determined using Student’s T-test.

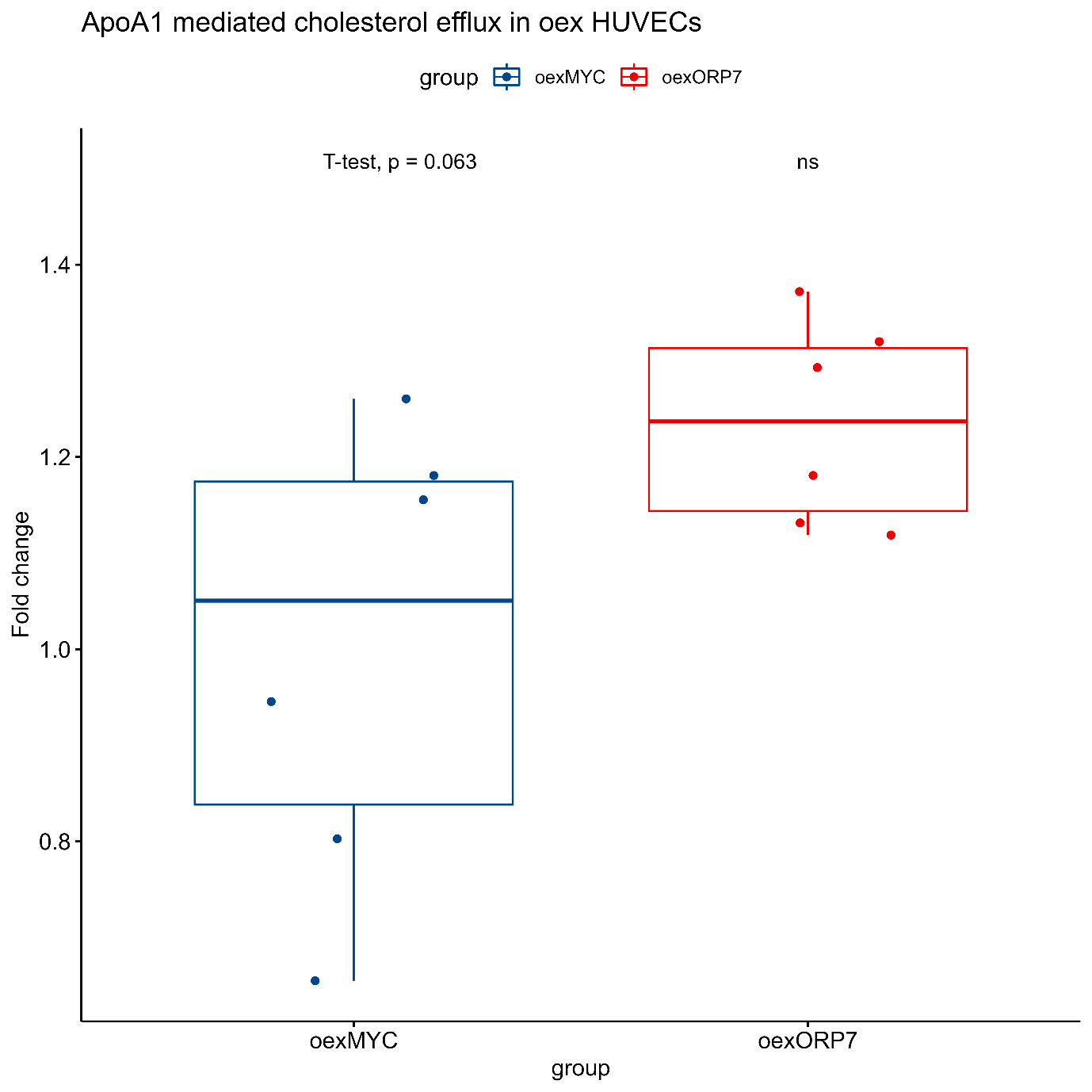

Figure S10. Box plots of cholesterol efflux fold change oex-cells. Y-axis represents the fold change compared to oexMYC control and the X-axis each group measured. A slight but non-significant increase in ApoA1 mediated cholesterol efflux is visible. P-values were determined using Student’s T-test and ANOVA.

ApoA1 mediated cholesterol efflux showed a slight increase in oex-cells but this change was statistically not significant.

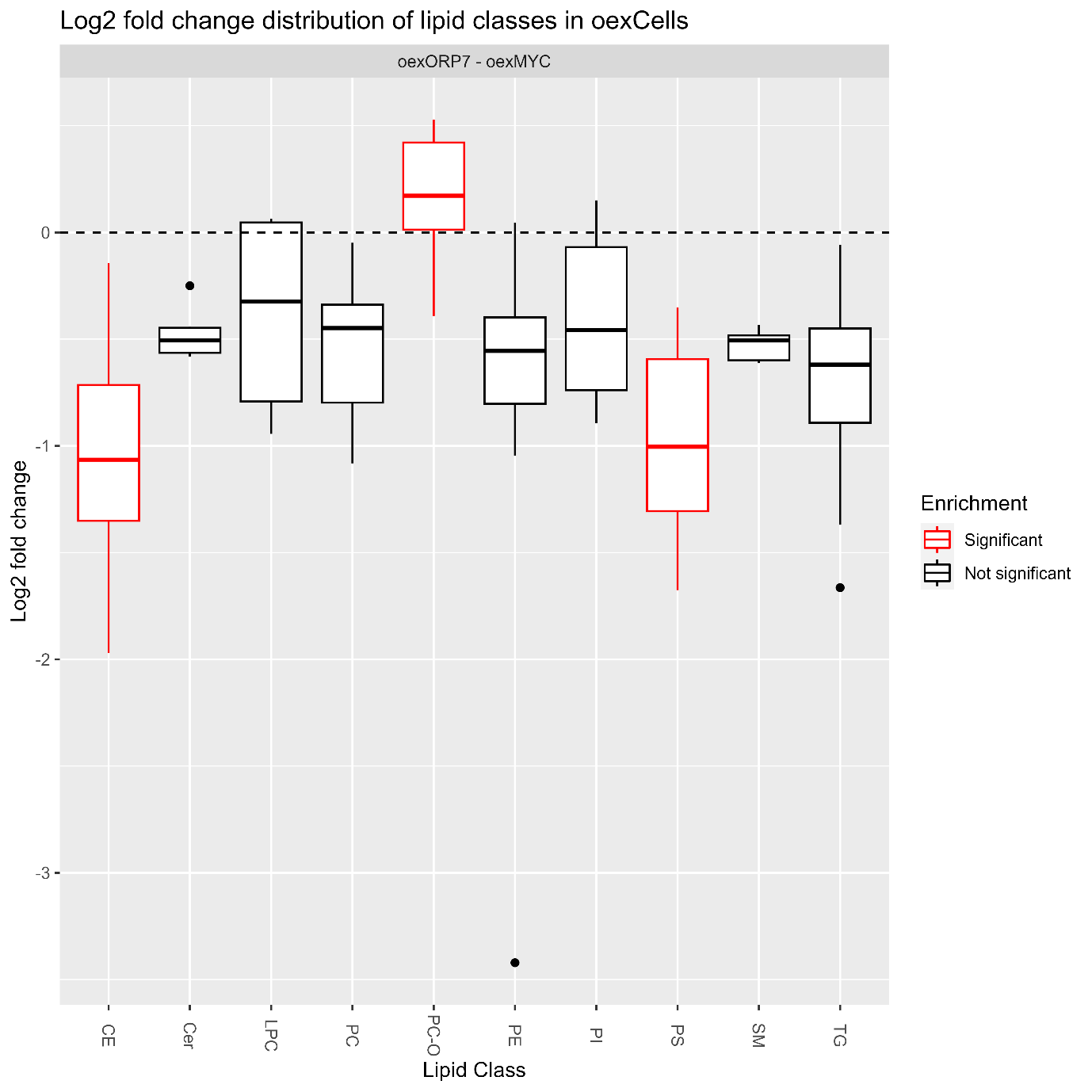

Figure S11. Box plot of log2 fold change distribution of different lipid classes in oex HUVECs. The Y-axis represents log2 fold change and X-axis each lipid class, significantly altered classes are shown in red. Oex-cells show significant changes in CEs,PC-O and PS.

Lipidomic analysis of oex-cells showed a similar decrease except more muted in CEs and a larger drop in PS to inhibited cells, as well as an increase in PC-Os in oex-cells.

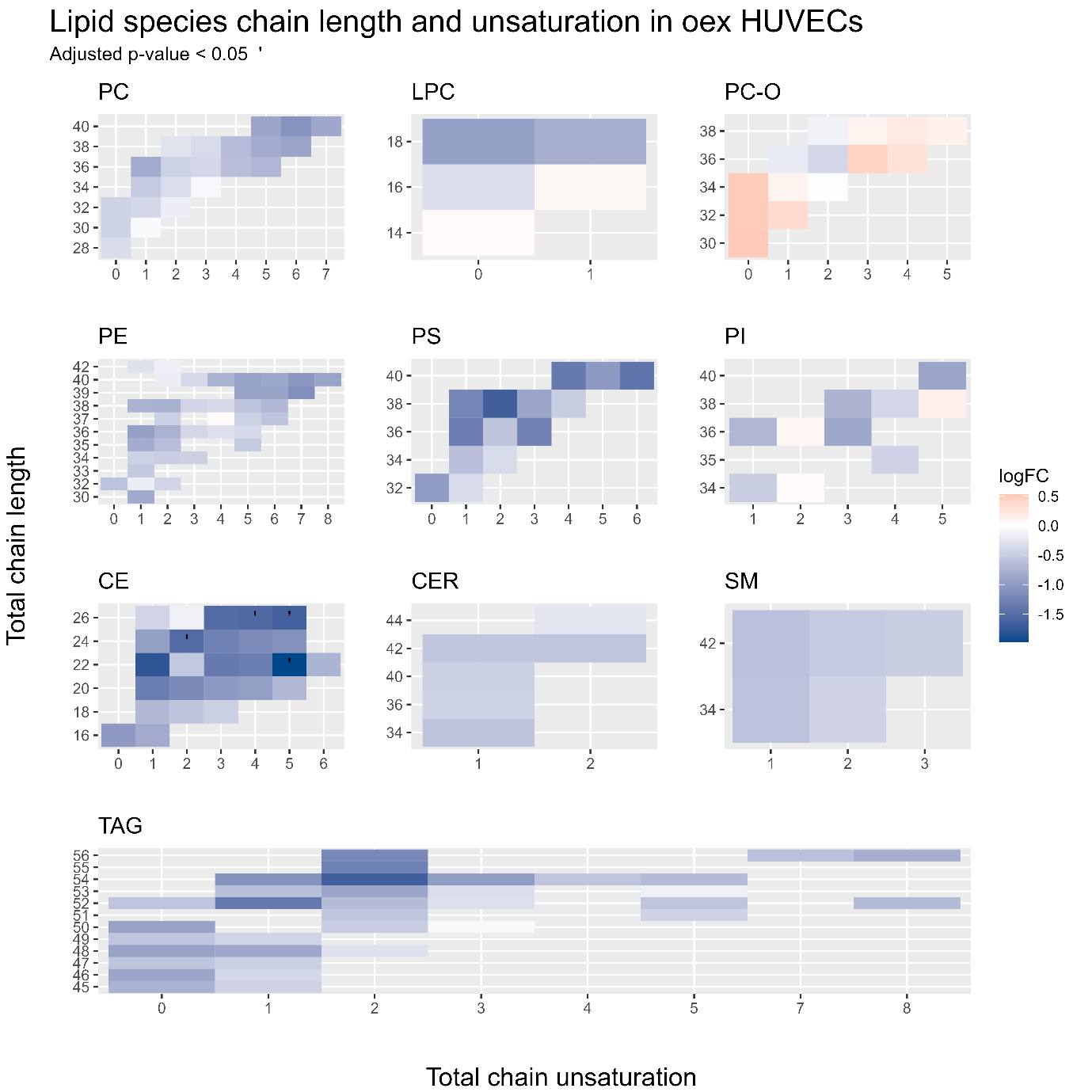

Figure S12. Tile plot of log2 fold changes in oex-cells. Each facet shows a different lipid class, where the Y-axis represents the total chain length, the X-axis total chain unsaturation and each tile depicts a different lipid species. Each tile is colored according to the log2 fold change of each species, where orange represents an increase in and blue a decrease, each statistically significantly altered species has been marked with a dot on the tile.

Even though CEs, PS and PC-Os showed significant changes as a group, Figure S12 clearly shows that the changes in individual lipids are not significant in any other class except for CEs, where alterations are concentrated to only a few lipid species.

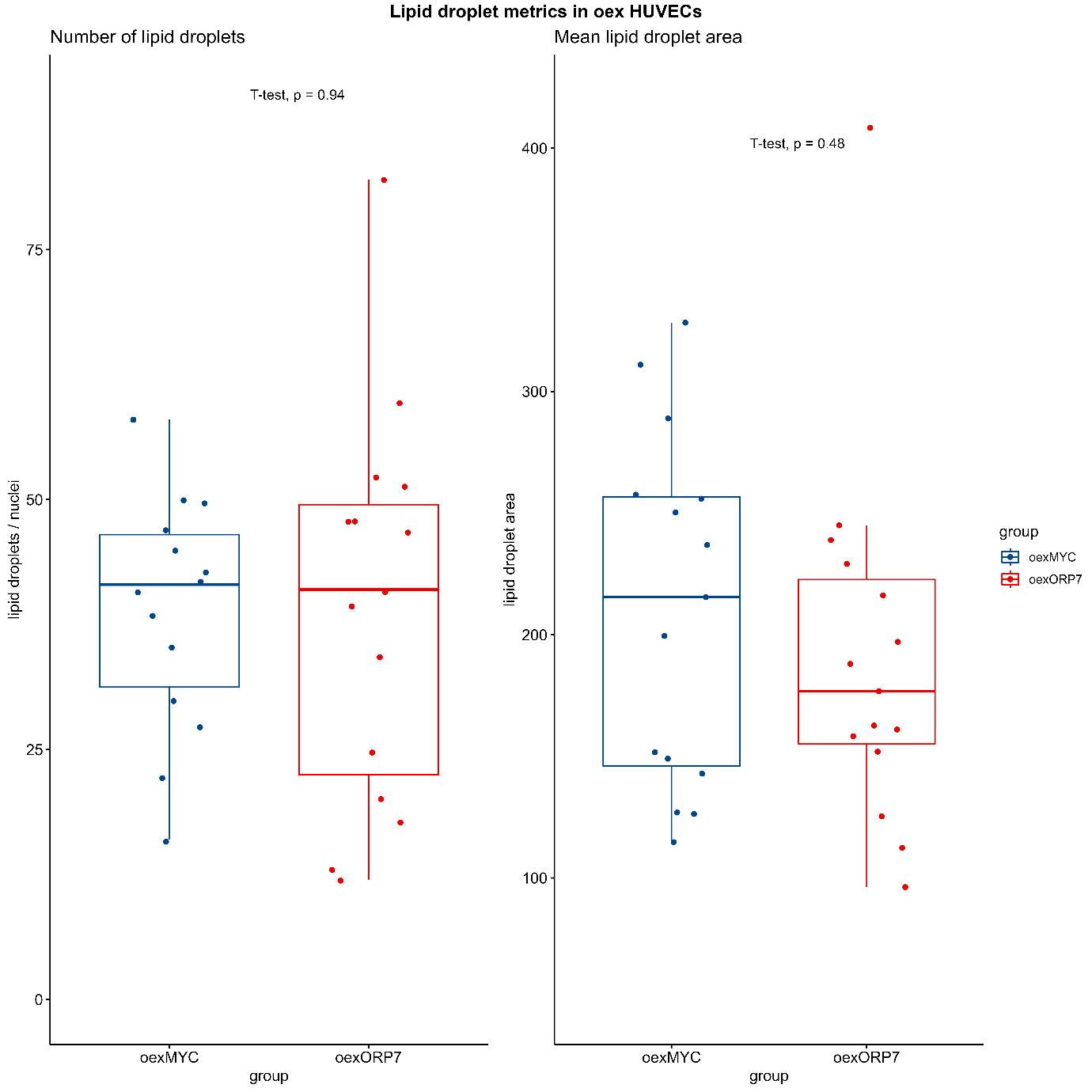

Figure S13, A box plot exhibiting lipid droplet metrics in oex-cells. Left and right facet show mean lipid droplet amount and mean lipid droplet area respectively. OexMYC control is shown in blue. No significant changes can be seen in either metric.

Lipid droplets showed no significant changes, in either the amount of lipid droplets or the mean area of lipid droplets.

Respiratory electron transport, ATP synthesis by chemiosmotic coupling, and heat production by uncoupling proteins.

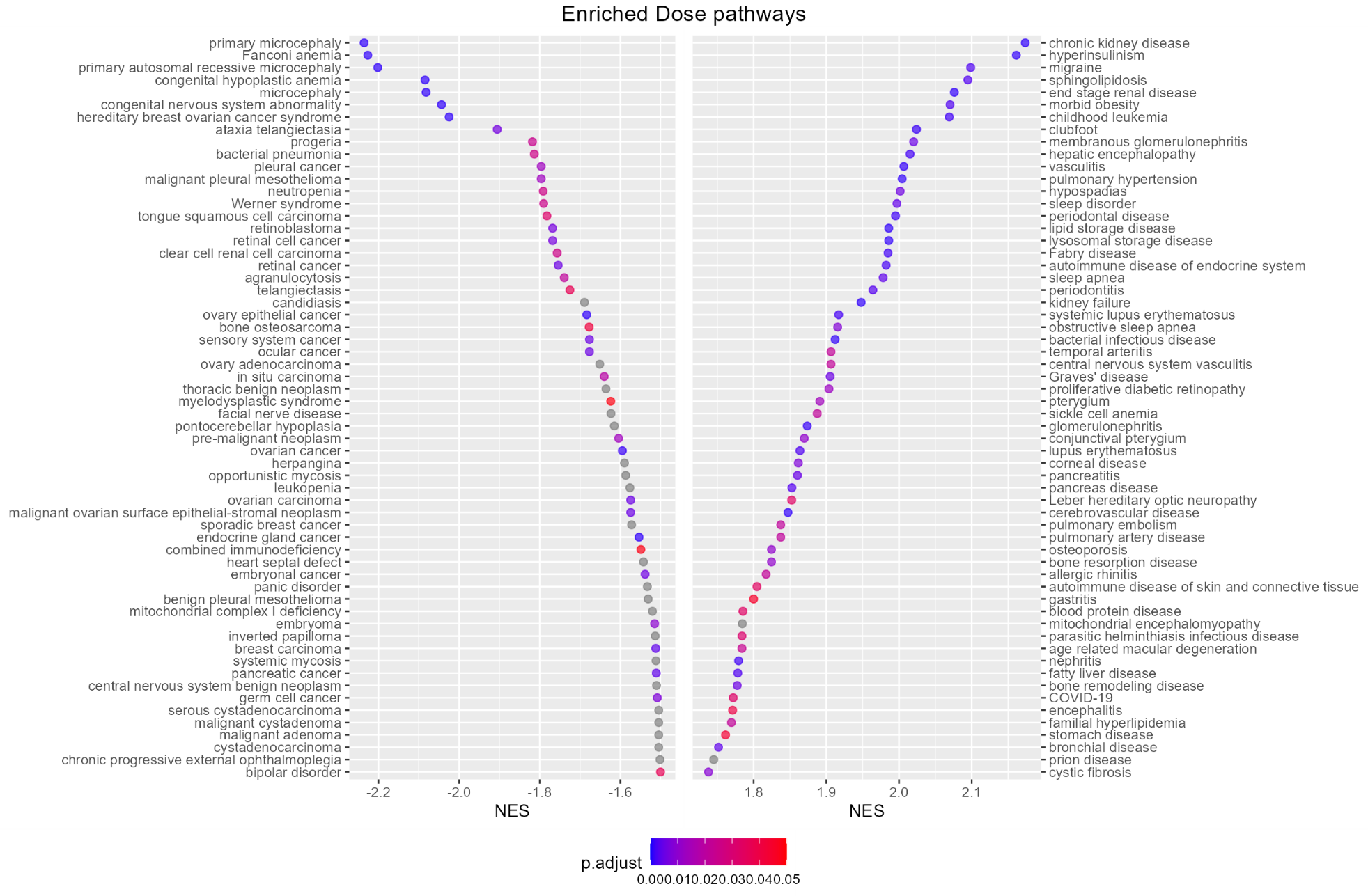
GSEA results on inhibitor treated cells.

Figure S14

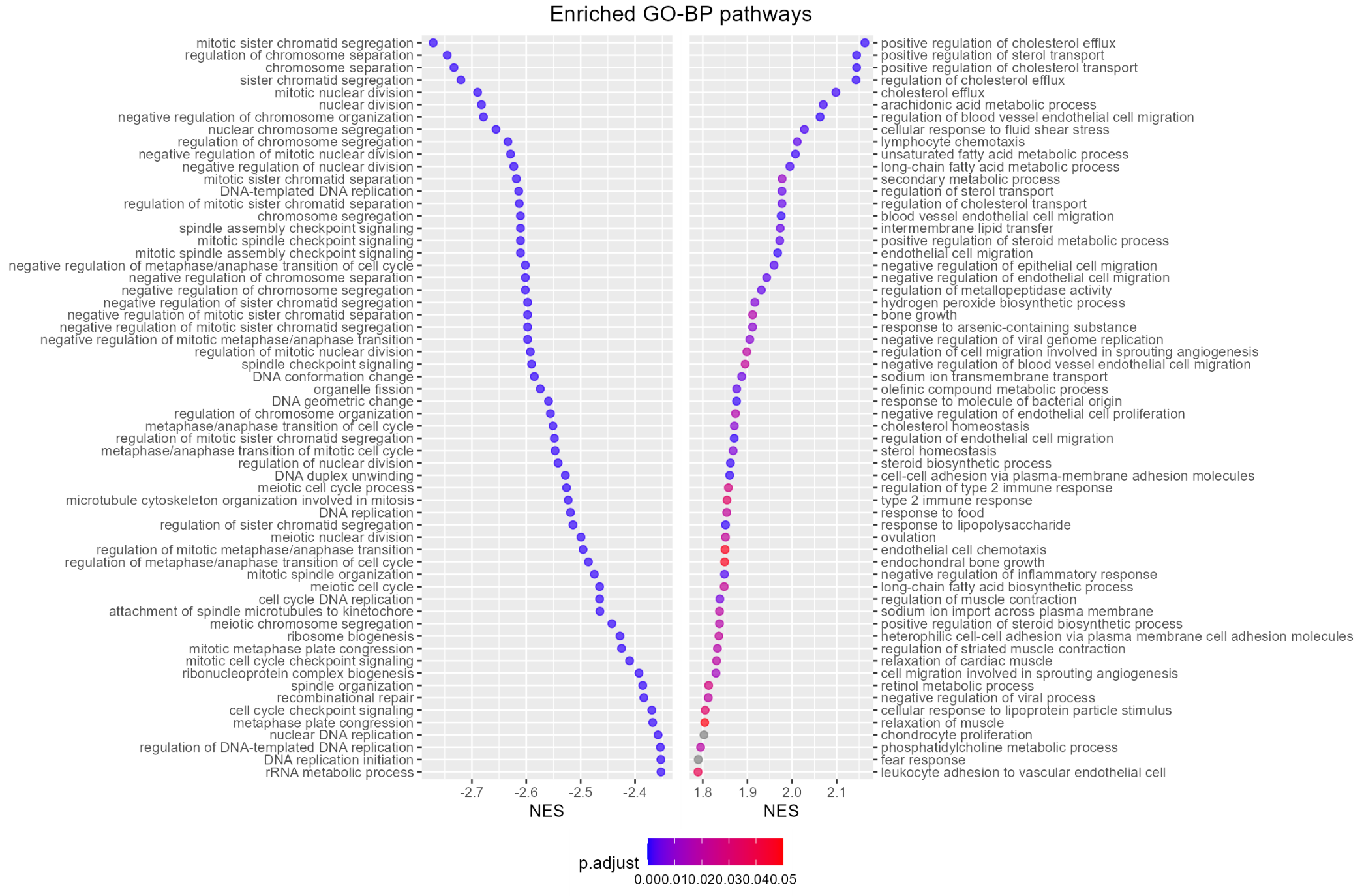
 Figure S15

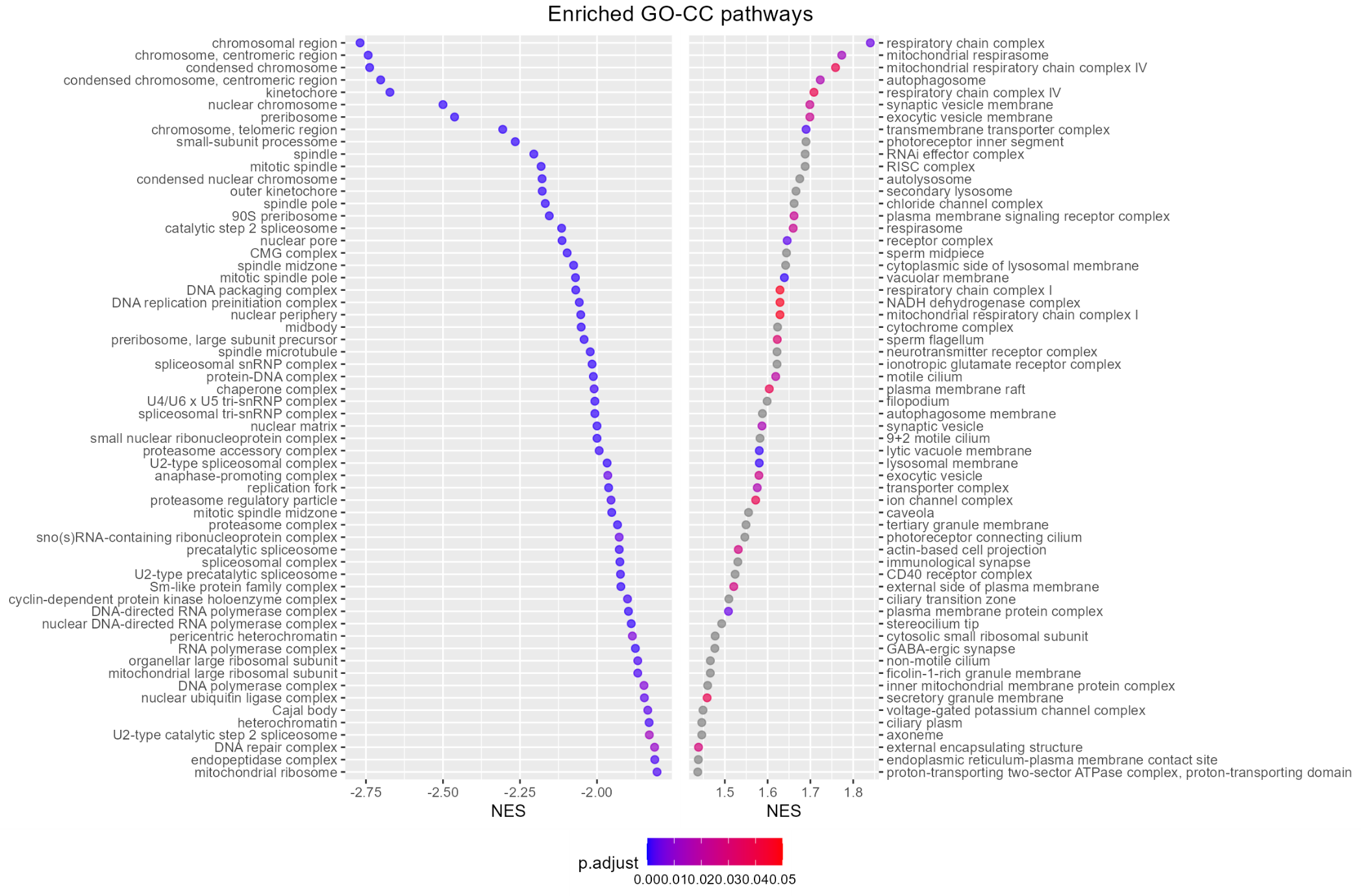
Figure S16

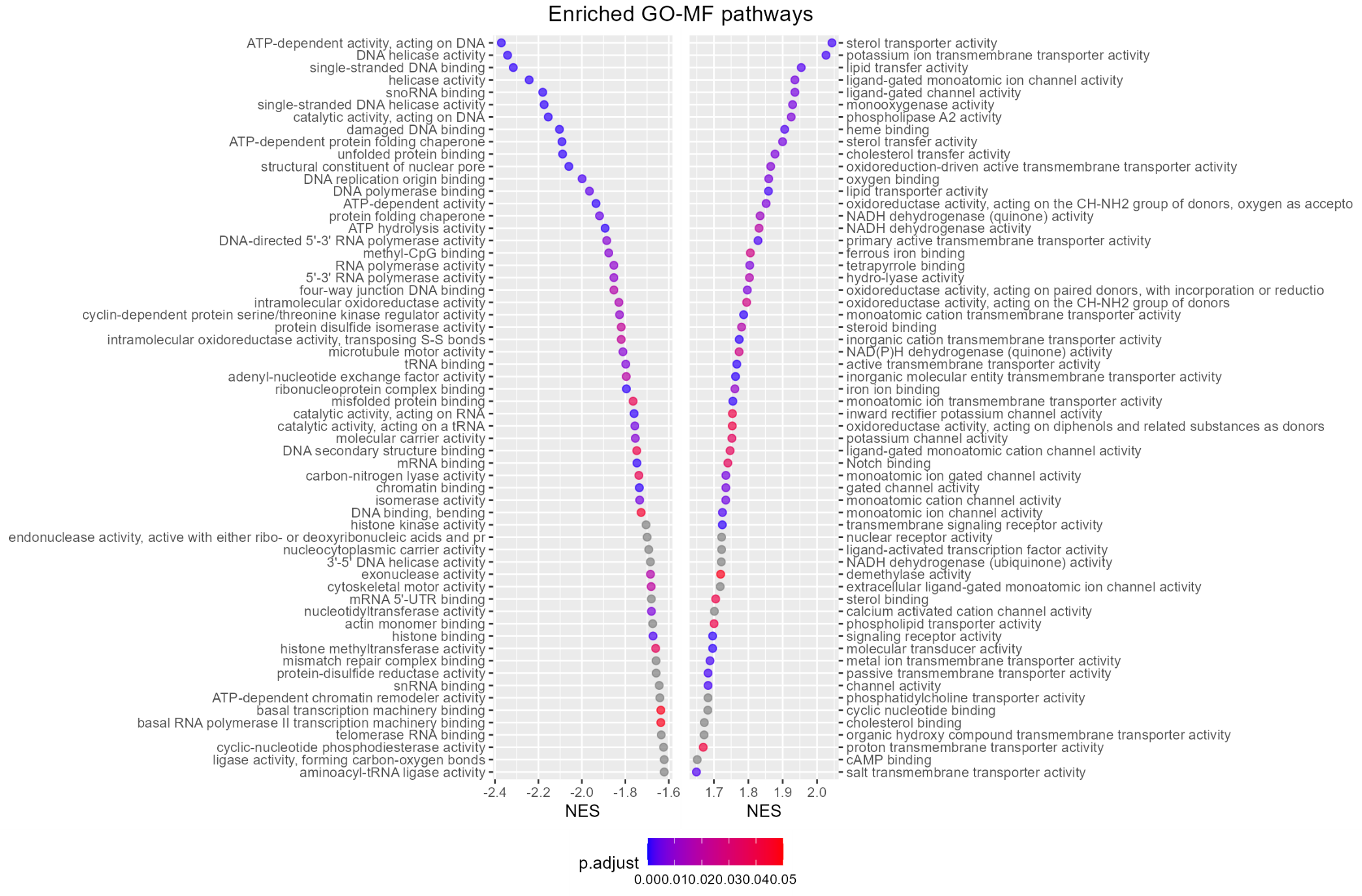
Figure S17

Figure S18
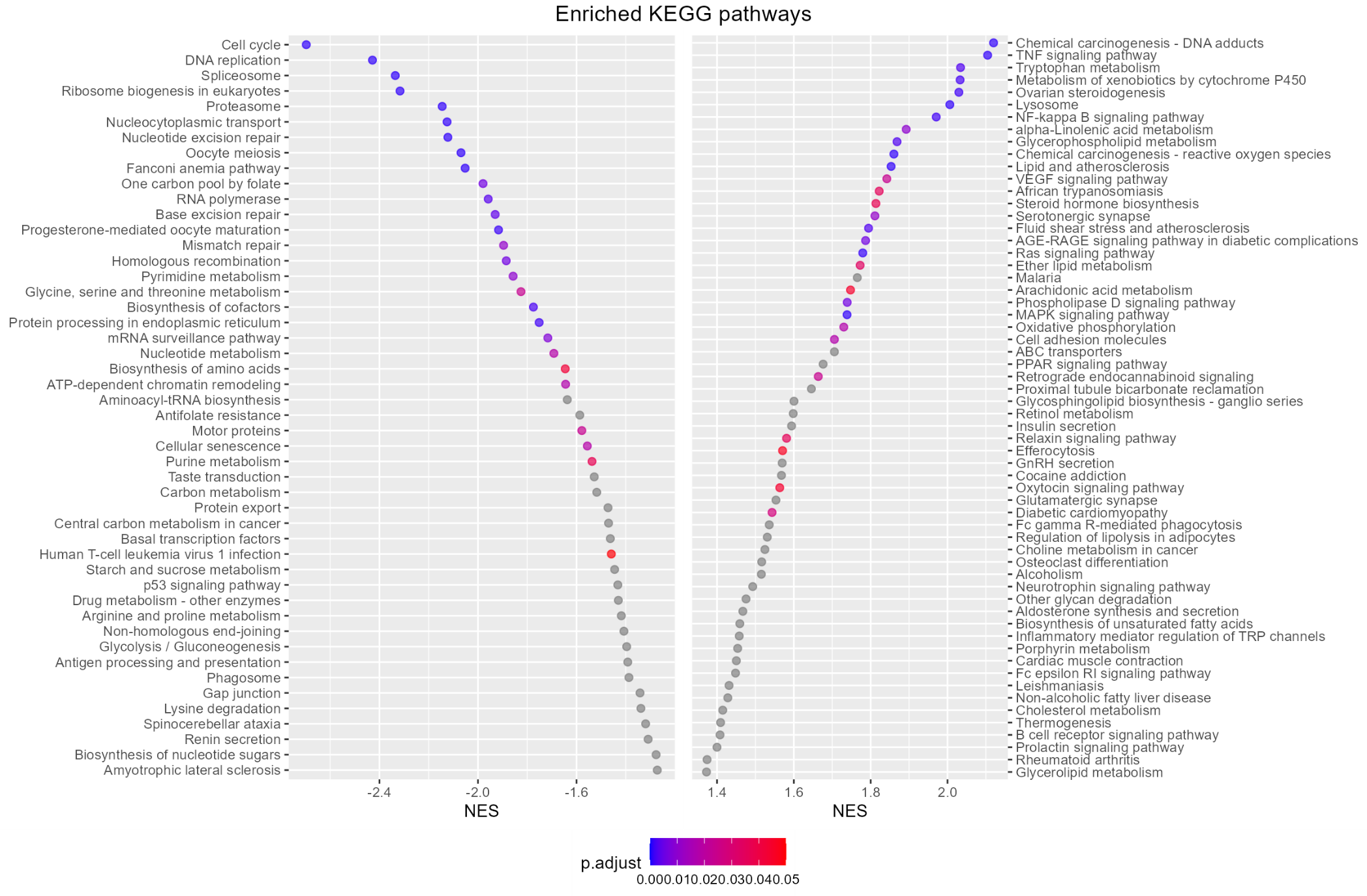

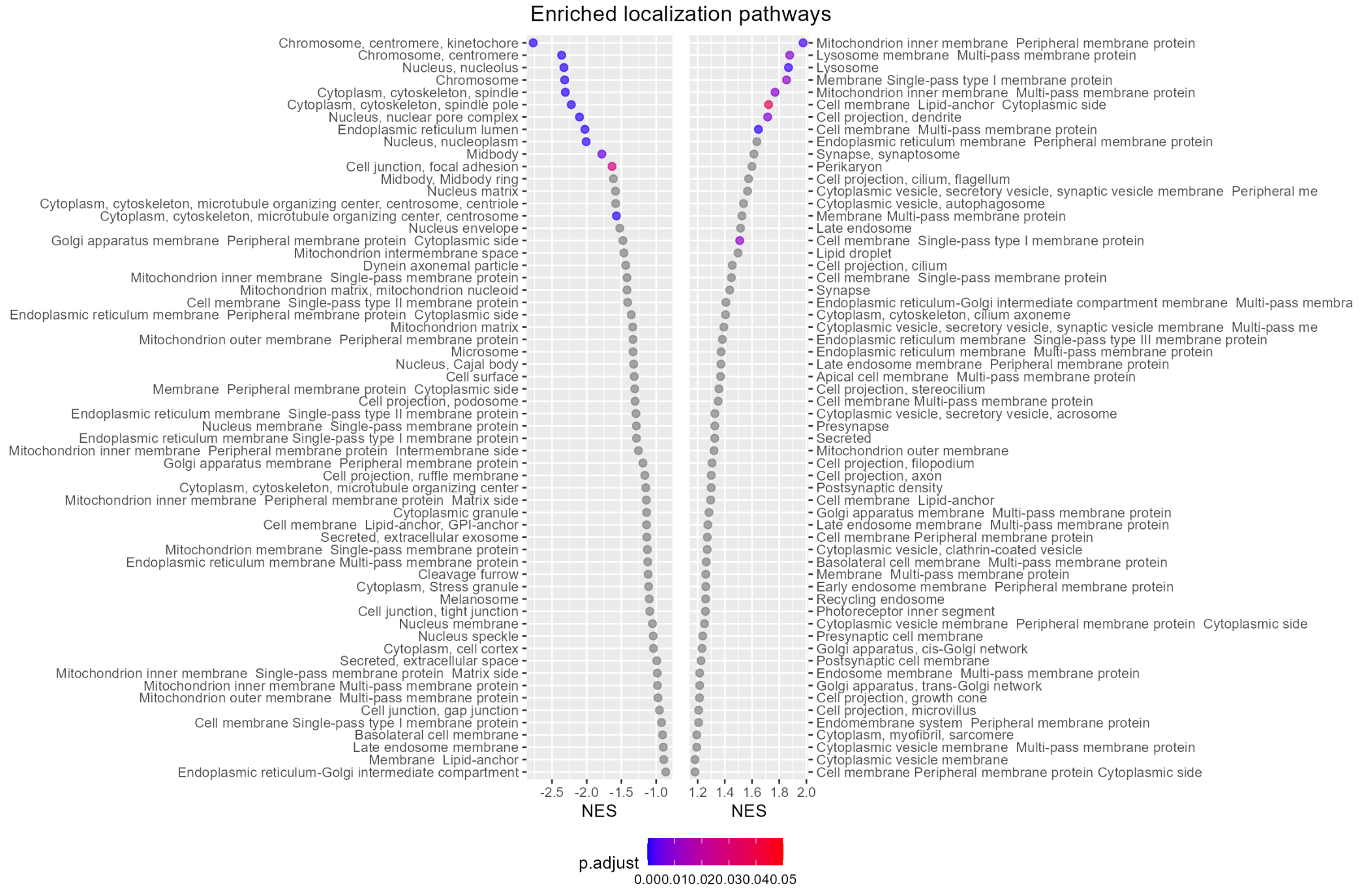
 Figure S19

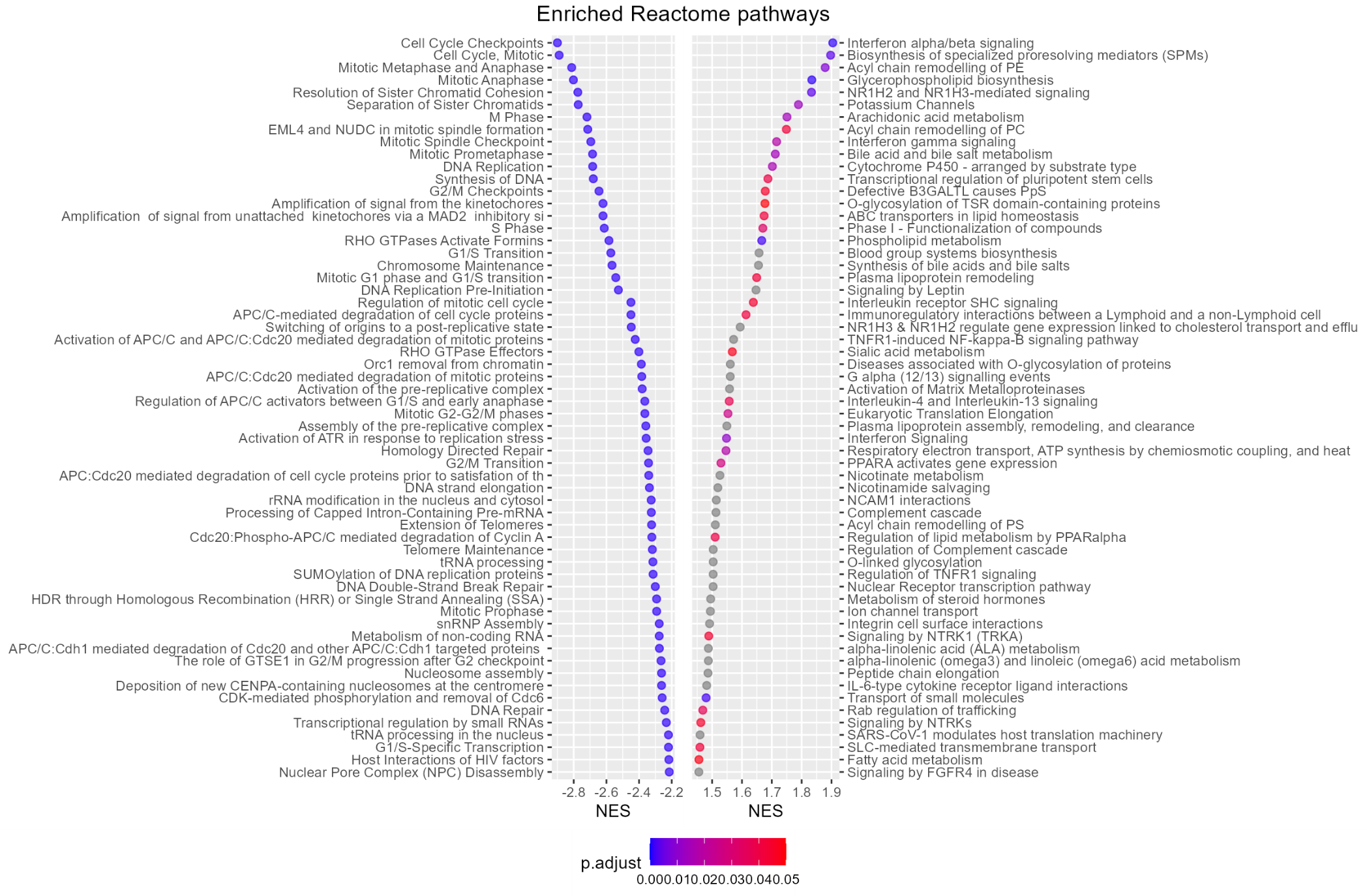
Figure S20

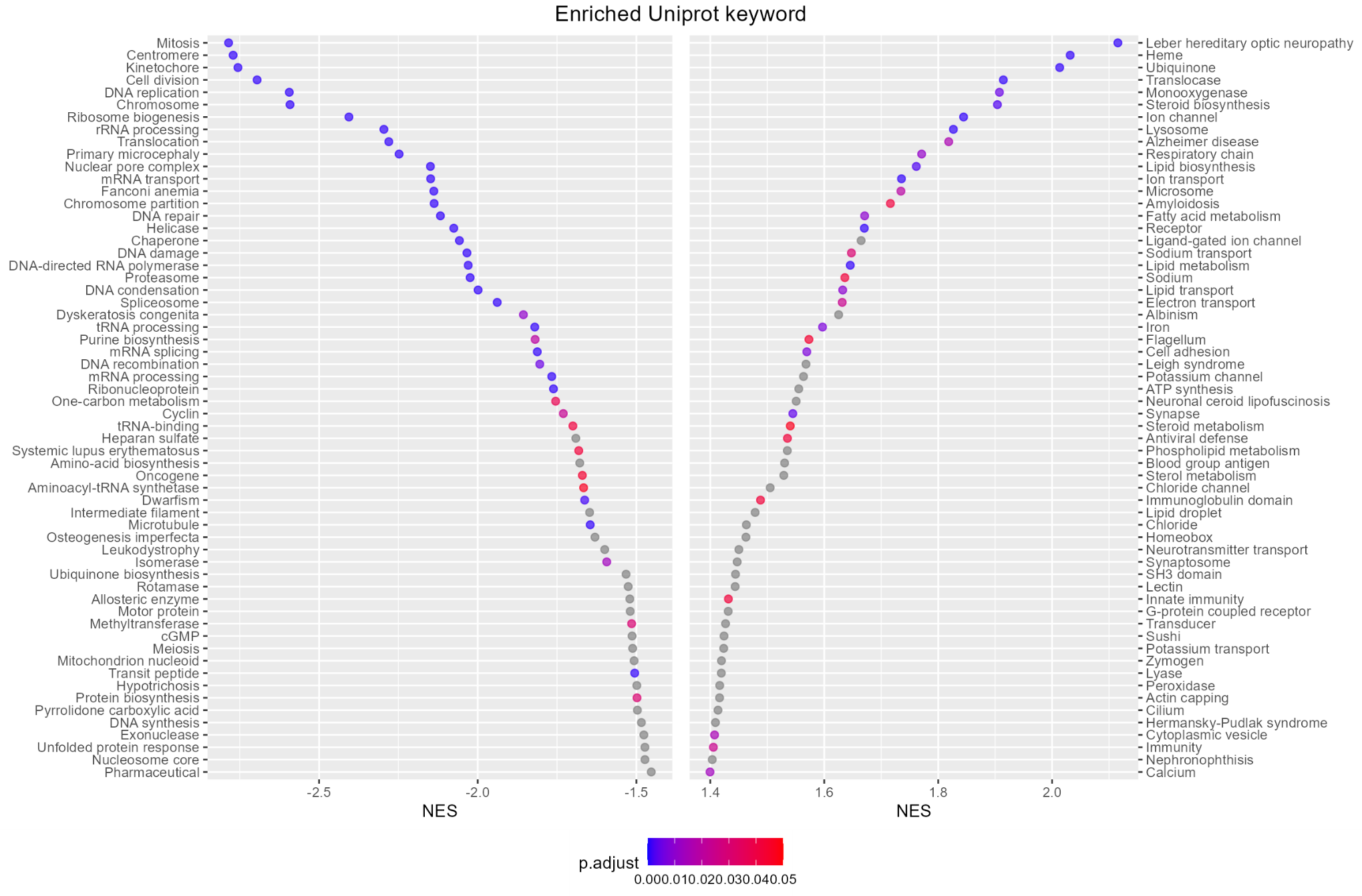
Figure S21

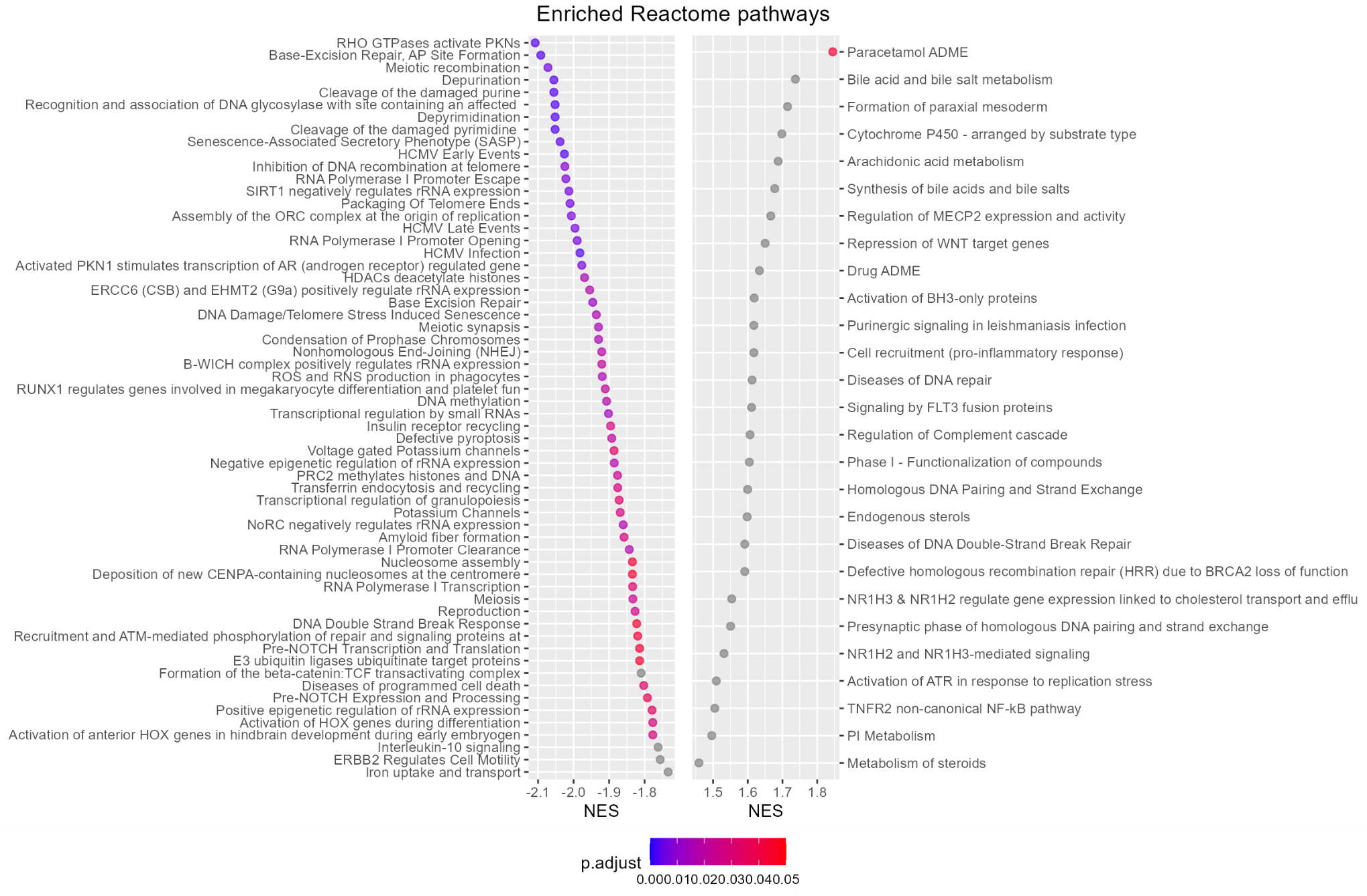
GSEA results on oexCells

Figure S22

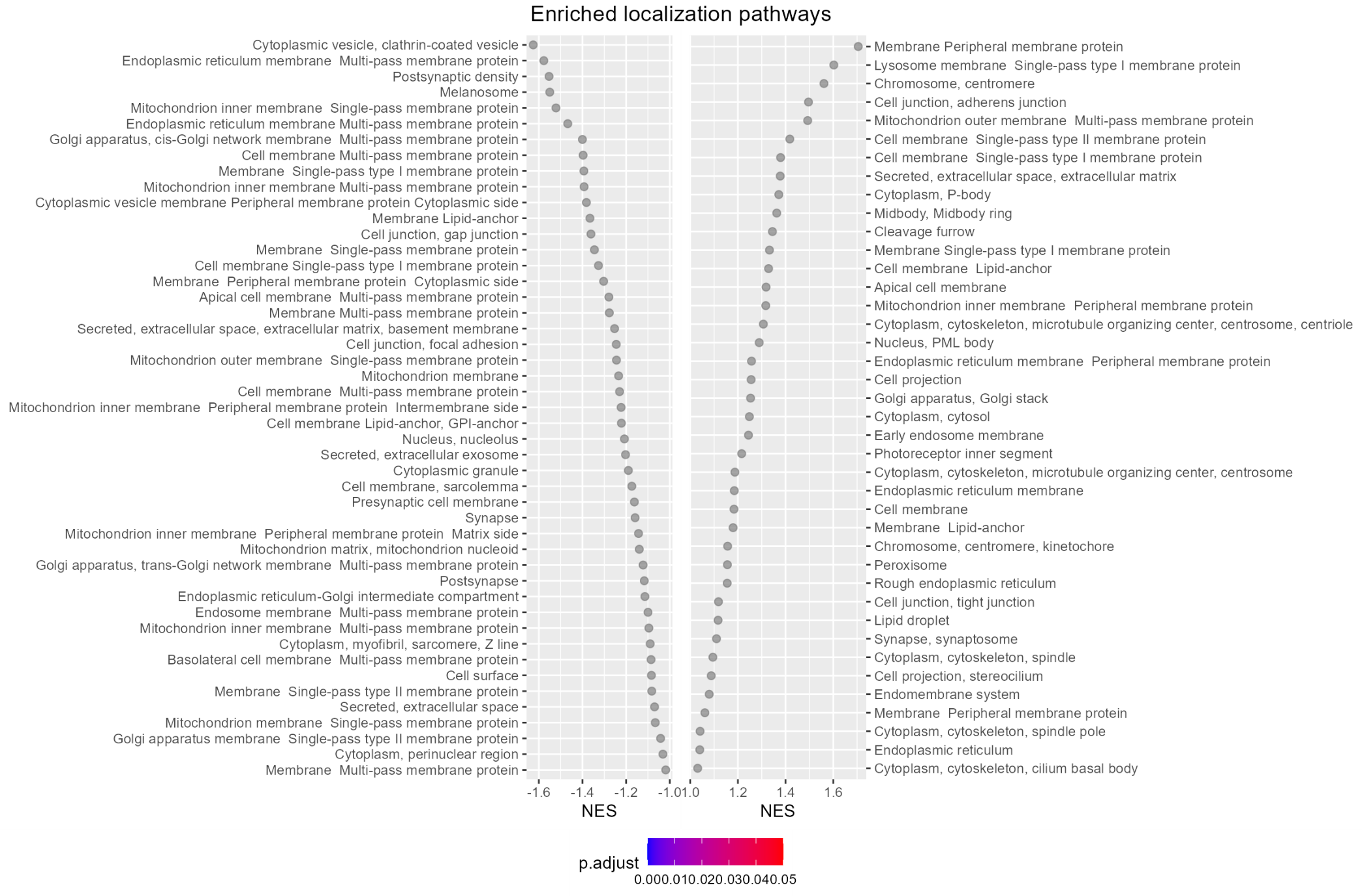
Figure S23

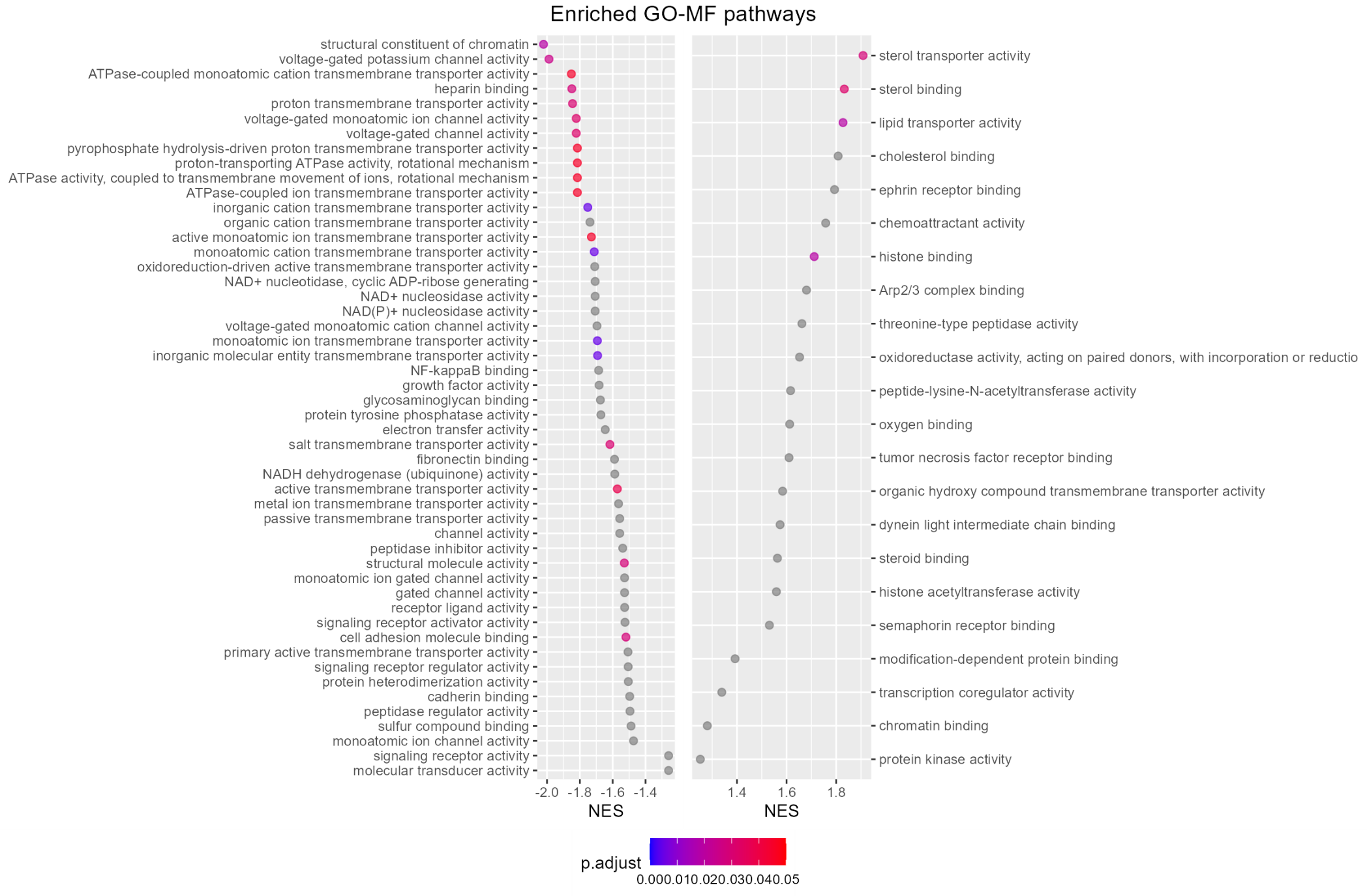
Figure S24

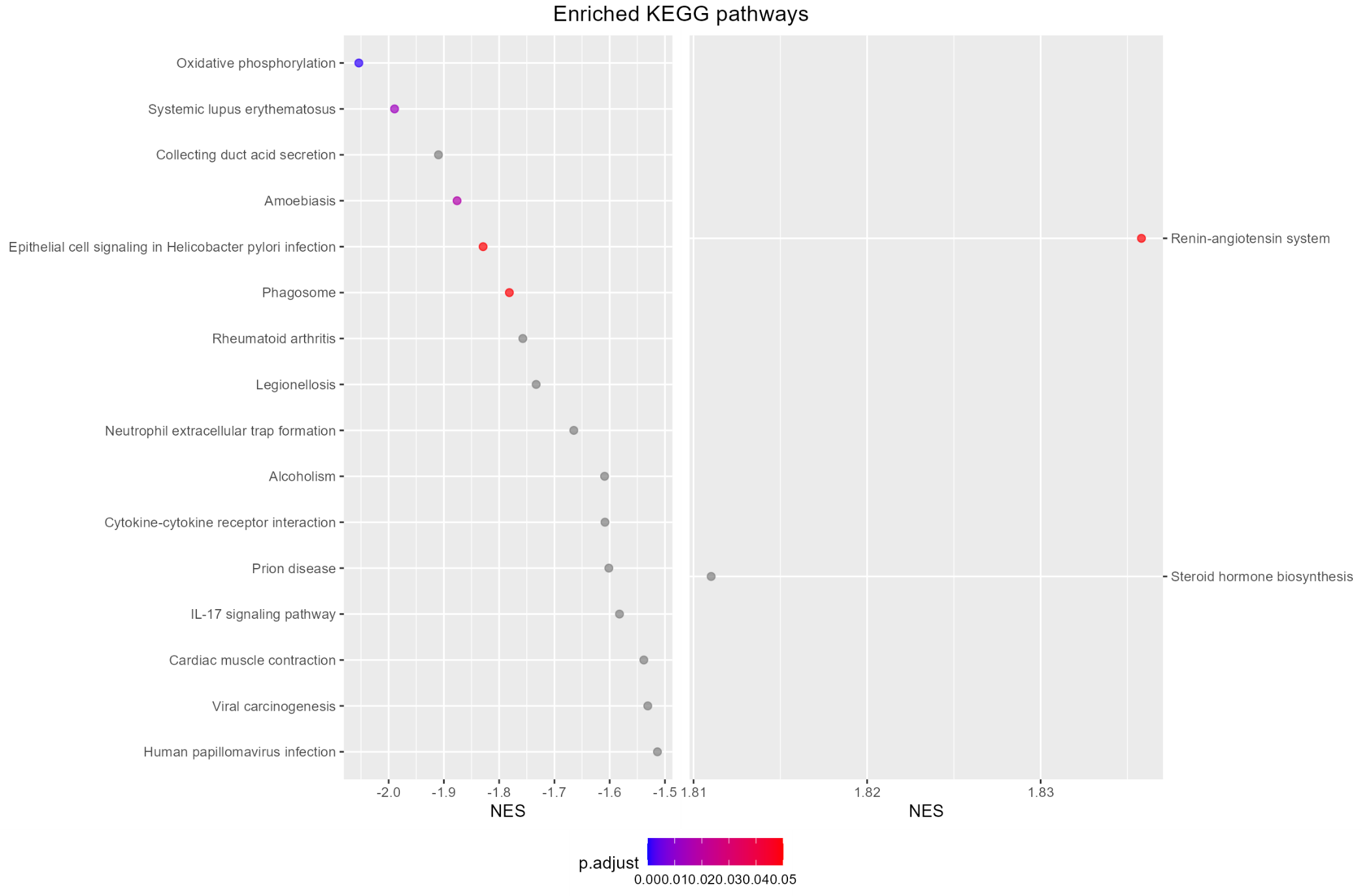
Figure S25

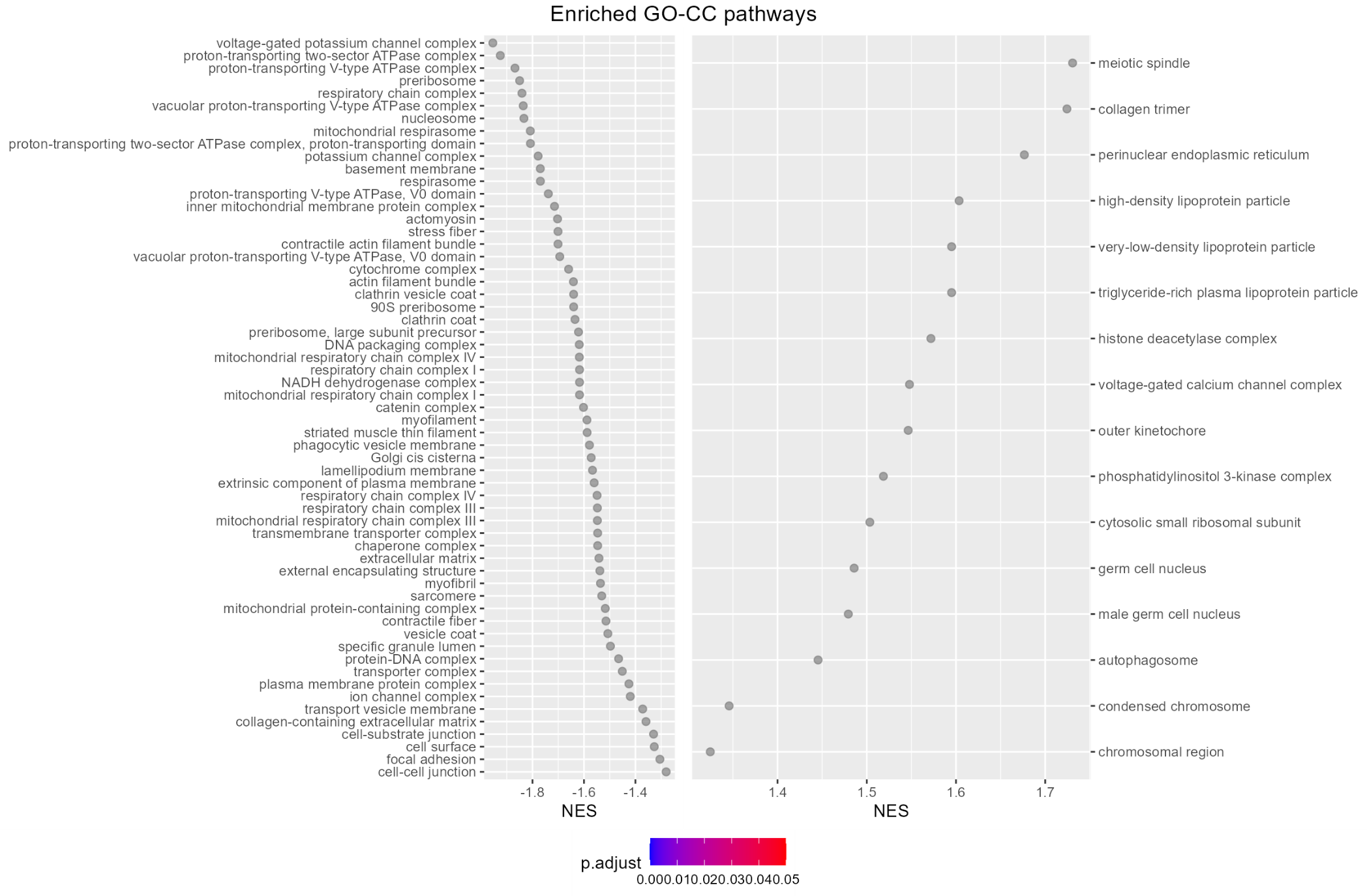
Figure S26

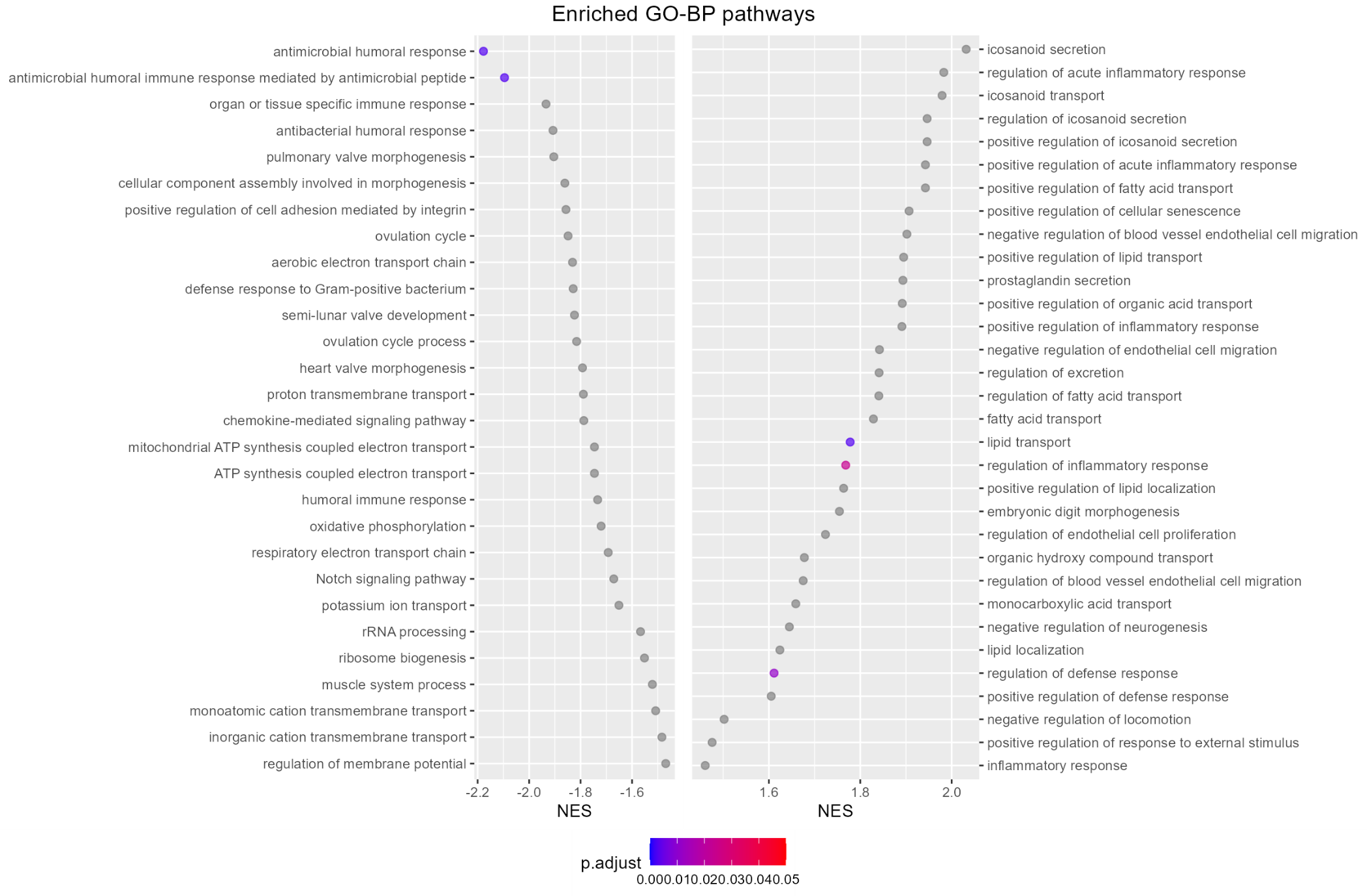
Figure S27

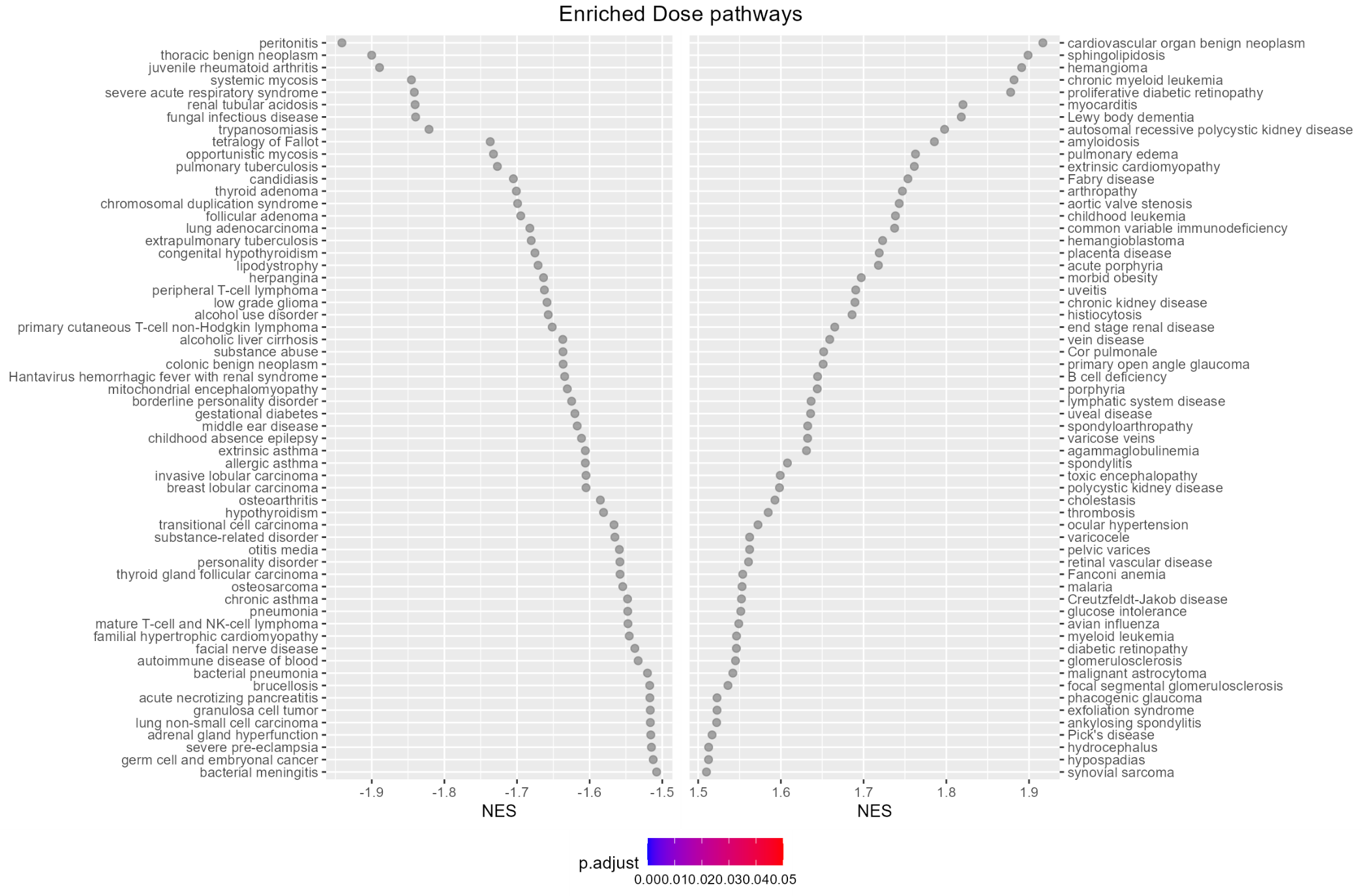
Figure S28

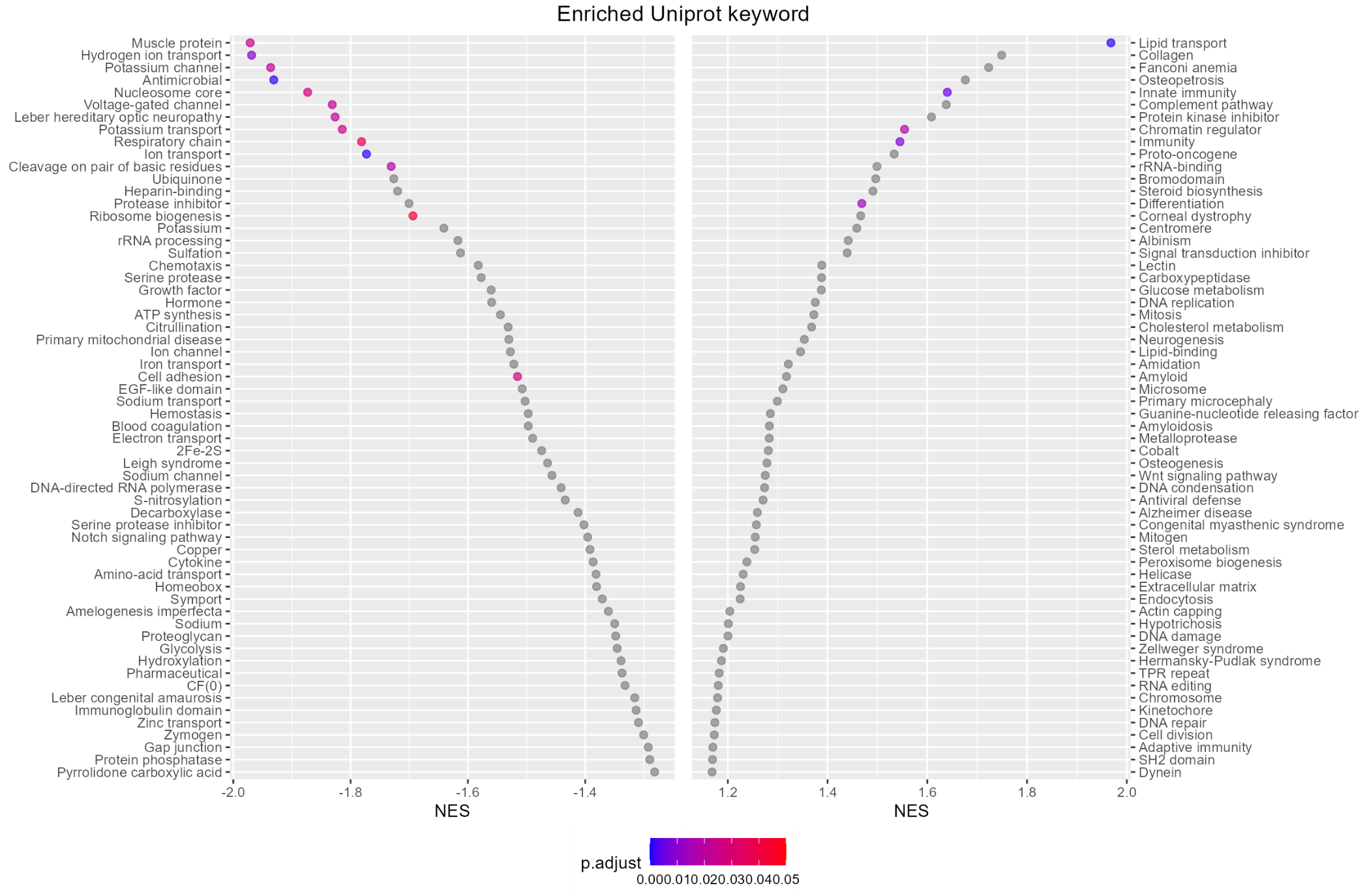
Figure S29
